# Dynamic Clonal Hematopoiesis and Functional T-cell Immunity in a Super-centenarian

**DOI:** 10.1101/788752

**Authors:** Erik B. van den Akker, Stavros Makrodimitris, Marc Hulsman, Martijn H. Brugman, Tanja Nikolic, Ted Bradley, Quinten Waisfisz, Frank Baas, Marja E. Jakobs, Daphne de Jong, P. Eline Slagboom, Frank J.T. Staal, Marcel J.T. Reinders, Henne Holstege

## Abstract

The aged hematopoietic system is characterized by decreased immuno-competence and by a reduced number of hematopoietic stem cells (HSCs) that actively generates new blood cell (age-related clonal hematopoiesis, ARCH). While both aspects are commonly associated with an increased risk of aging-related diseases, it is currently unknown to what extent these aspects co-occur during exceptional longevity. Here, we investigated these aspects in blood cells of an immuno-hematopoietically normal female who reached 111 years. Blood samples were collected across a 9-year period at ages 103, 110 and 111 years. We applied several genetic sequencing approaches to investigate clonality in peripheral blood samples and sorted cell subsets. Immuno-competence was characterized using flow cytometry, T-cell receptor excision circle (TREC) assays, and in vitro proliferation assays. We identified a single *DNMT3A*-mutated HSC clone with a complex subclonal architecture and observed ongoing subclonal dynamics within the 9-year timeframe of our sampling. The mutated HSC generated 78-87% myeloid cells, 6-7% of the B-cells, 6% of CD8^+^ T-cells, and notably 22% of the CD4^+^ T-cells. Intriguingly, we found that T-cells were capable of robust proliferation when challenged *in vitro*. Moreover, we observed a surprisingly high TREC content, indicative of recent generation of naive T-cells. Concluding, we observed long-term stability of extreme ARCH with ongoing clonal dynamics combined with functional T-cell immunity. Our results indicate that extreme ARCH does not compromise immuno-competence and that a clonally expanded CD4+ T-cell subset may serve as a potential hallmark of the supercentenarian immune system.

**Key points:**

1. Longitudinal blood sampling from a female aged 103-111 revealed a dynamic clonal hematopoiesis contributing to myeloid and lymphoid subsets
2. Despite the highly advanced age and extreme clonal hematopoiesis we observed functional T-cell immunity

## Introduction

Age-related Clonal hematopoiesis (ARCH) is an inevitable physiological consequence of ageing^1^, conferring a tenfold increased risk for the development of myeloid dysplasia or malignancies^2,3^. In addition to hematological disease, large prospective epidemiological studies have also established associations between ARCH and prevalent type 2 diabetes^4^, prevalent chronic obstructive pulmonary disease^1^, incident cardiovascular disease^5^, and vascular or all-cause mortality^1-3,5^. While these associations suggest a potential role for ARCH not only in leukemia, but also in a broad spectrum of age-related low-grade inflammatory syndromes^6^, its heterogeneity with respect to these clinical outcomes has also been noted^7-10^. For instance, ARCH has been postulated to constitute an early phase of myelodysplasia and leukemia^7^, yet irrespective of ARCH status, the absolute risk to progress to one of these clinical entities remains overtly small. In addition, while ARCH becomes increasingly prevalent with age^2,3,11^, and can become very extreme in the exceptionally old^12^, its association with all-cause mortality seems to wane in these oldest old^9^. Collectively, these findings illustrate that we are far from understanding the molecular pathophysiological transitions that ARCH may initiate during ageing.

ARCH arises when an ageing hematopoietic stem cell (HSC) acquires a somatic mutation that confers a competitive growth advantage, leading to its gradual expansion^13^. ARCH-associated mutations typically target genes previously linked with myeloid aberrancies, most prominently in genes encoding epigenetic regulators such as *DNMT3A, TET2* and *ASXL1*^2,3,13^. Nevertheless, previous reports indicate that a significant part of ARCH occurs in the absence of such known drivers (ARCH-UD)^1,3^. These studies exploited the fact that HSCs accumulate somatic mutations over the course of a lifetime^14^, thus effectively tagging each individual HSC with a unique ‘genetic barcode’^1^. We previously applied this paradigm to trace the clonal origins of peripheral blood cells taken from a 115-year-old female, which revealed that up to 65% of her peripheral blood was derived from a single expanded HSC^12^. While this study exemplified that extreme ARCH does not preclude attaining an exceptional age, it also raises questions on the characteristics of this particular type of clonal hematopoiesis that sets it apart from other forms that eventually do lead to disease.

Studies on the ageing immune system and its relation to age-related disease have traditionally focused on the lymphoid branch, also referred to as ‘immuno-senescence’, which is predominantly characterized by the age-related loss of functionality of the T-cell compartment. Hallmarks of immuno-senescence involve the age-associated changes in circulating CD4+ and CD8+ T-cell populations, including the lack of naïve T-cells, increased numbers of memory T-cells, decreased complexity in the antigen-recognizing repertoire, and decreased cell proliferation upon encountering an antigen^15^. Nearly all of these changes can be attributed to a reduced production of T-cells in the thymus^16^, which gradually ceases to function after puberty and is typically fully involuted at advanced age. Thymic proliferation can be assessed by quantifying the concentration of T-cell receptor excision circles (TRECs) in peripheral blood or in sorted immune subsets^17-19^. TREC content has been shown to decline sharply after middle age and often becomes undetectable at old age (>85 years)^20^. Nevertheless, human studies on the ageing immune system typically do not include both these assessments of changes in T-cell proliferation or in hematopoietic clonality (ARCH). Consequentially, it remains elusive to what extent immuno-senescence and ARCH co-occur in the ageing human immune system.

We address this question by performing a detailed study on peripheral blood and its sorted subsets drawn from an immuno-hematologically normal individual at age 103, 110 and 111 years. In accordance with our previous findings in a 115-year old individual, deep sequencing revealed extreme ARCH: over 75% of the peripheral blood cells shared their clonal origin. We subsequently employed this unique study sample to characterize various aspects of extreme ARCH, including its subclonal architecture, its temporal stability, and its potential restriction to particular immune lineages. In parallel, we investigated various aspects of immuno-senescence using flow-cytrometry, TREC assays and *in vitro* proliferation experiments. Collectively, this study represents a comprehensive exploration of the ageing immune system at the extreme end of human lifespan.

## Methods

Full details of the methods used are given in the **Supplementary Methods**.

### Study design and data collection

Subject of this study was W111, a Dutch female who lived for 111 years and 10 months. W111 was enrolled as a participant of the Leiden Longevity Study^21^ and seven years later as a participant of the 100-plus Study^22^. 100-plus Study and Leiden Longevity Study have been approved by their respective local Medical Ethical Committees and all participants, including the current subject, gave informed consent for study participation.

### Clinical diagnosis of hematological status

To rule out myeloid dysplasia, we investigated whether the DNA derived from a peripheral blood sample collected at age 110 had accumulated a mutation in one or more of the 54 genes frequently mutated in myeloid neoplasia. For this we used the TruSight Myeloid panel (Illumina) and re-sequenced on a MiSeq instrument (Illumina) at >500x read depth (**Supplementary Methods 1**). Furthermore, the presence of clonal BCR or TCR recombinations in peripheral blood may point to suspect lymphoproliferations, or a recent or sustained antigenic stimulation. The diversity in both T and B-cell receptor repertoire was investigated using IdentiClone™ PCR assays followed by quantification using an ABI Capillary Electrophoresis Instrument.

### Detection of somatic mutations in blood

To detect somatic mutations in W111 blood, we analyzed DNA derived from peripheral blood and a skin biopsy collected on the same day using Whole Genome Sequencing (WGS) using SOLiD sequencing technology (Life Technologies) at 80x mean depth (**Supplementary Methods 2**). Single Nucleotide Variations (SNVs) were called in both samples using GATK Haplotypecaller^23^ and somatic mutations were identified using *in-house* software Sommix (**Supplemental Methods 3**).

### Targeted amplicon Sequencing

To validate the somatic origin of the identified mutations we used Ion Torrent amplicon re-sequencing at 6000x average read-depth (Proton Ampliseq sequencing, Thermo Fisher Scientific, **Table S1, Supplementary Methods 4**), contrasting blood with brain-cortex. Variants were called using Torrent Variant Caller (TVC) version 5.0.0 (Life Technologies).

### Annotation of somatic variants

Somatic variants were annotated with UCSC’s Variant Annotation Integrator and compared to previously reported candidate driver definitions (**Supplementary Methods 5**,**Table S2**). Analyses of their tri-nucleotide context and resemblance to previously published and curated mutational signatures (COSMIC, V2 - March 2015) was performed with the R package MutationalPatterns^24^.

### Subclonal architecture

Somatic mutations were clustered by their Variant Allele Frequencies (VAFs) using SciClone version 1.1.0^25^ (**Table S3**). Resulting cluster annotations and VAFs were subsequently analyzed with SCHISM version 1.1.2^26^, to infer a phylogenetic tree describing the most probable order in which subclonal expansion events (clusters of SNVs) occurred. The percentage of cells carrying a set of mutations marking a specific subclonal expansion event was computed using the median VAF across all assigned mutations. To obtain the percentage of cells that belong to a particular subclone, the median VAF across the mutations marking a specific expansion event were corrected for the median VAF across the mutations marking the immediate child subclonal expansion event(s) in the tree, e.g. A’ = A – B.

### Immuno-phenotyping

Extensive immuno-phenotyping was performed on peripheral blood using flow cytometry, T-cell receptor excision circle (TREC) assays), and mixed leukocyte reactions (**Supplementary Methods 6-8**).

## Results

Blood samples from W111 were collected at three time points, age 103 (timepoint 0), 110 (timepoint 1), and 111 (timepoint 2) respectively (**Figure 1**), and included peripheral blood (PB), its flow sorted subsets: granulocytes (G), monocytes (M), T-cells (T), CD4^+^ T-cells (T4), CD8^+^ T-cells (T8) and B-cells (B). Additionally, a skin biopsy (S) was collected at age 110, and the subject agreed to post-mortem brain donation, allowing the investigation of brain-cortex (C). Her blood showed no signs of cytopenia, dysplasia, or other clinical signs of haematological malignancies.

**Fig 1.**
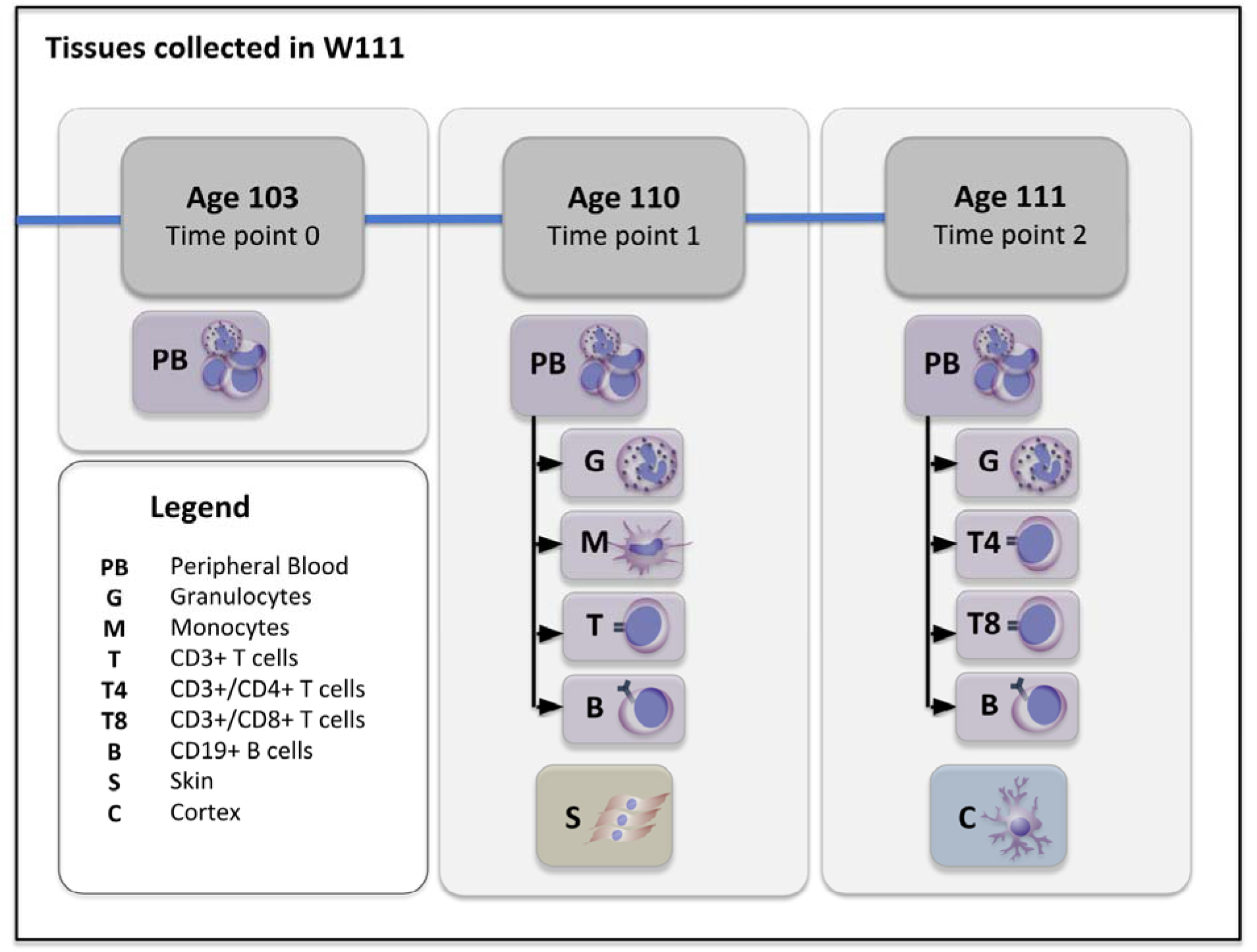
Study design: An overview of the tissues collected at three different time points.

### Sequencing peripheral blood reveals extreme age-related clonal hematopoiesis

To identify somatic mutations that had accumulated in the hematopoietic stem cell compartment of W111, we compared WGS of DNA derived from the peripheral blood collected at age 110 years and 3 months (PB1) with WGS of DNA derived from the skin biopsy collected on the same day (**Methods**). We identified 650 putative single nucleotide somatic variations (SNVs) (**Figure 2A, Table S1**). Concurrent with previous work on the non-leukemic accumulation of somatic mutations in peripheral blood^12^ or in cultures of single expanded hematopoietic stem cells^14,27,28^, the vast majority of these mutations were non-coding (**Figure 2B**), did not exhibit any positional preferences in the genome (**Figure S1**), and were dominated by C>T and T>C transitions (**Figure 2D**). Moreover, the tri-nucleotide sequence context of the identified mutations (**Figure 2E, Figure S2**) largely overlapped with established ‘clock-like’ mutational signatures that represent relatively benign ageing processes^28,29^ (**Figure S3**).

**Figure 2:**
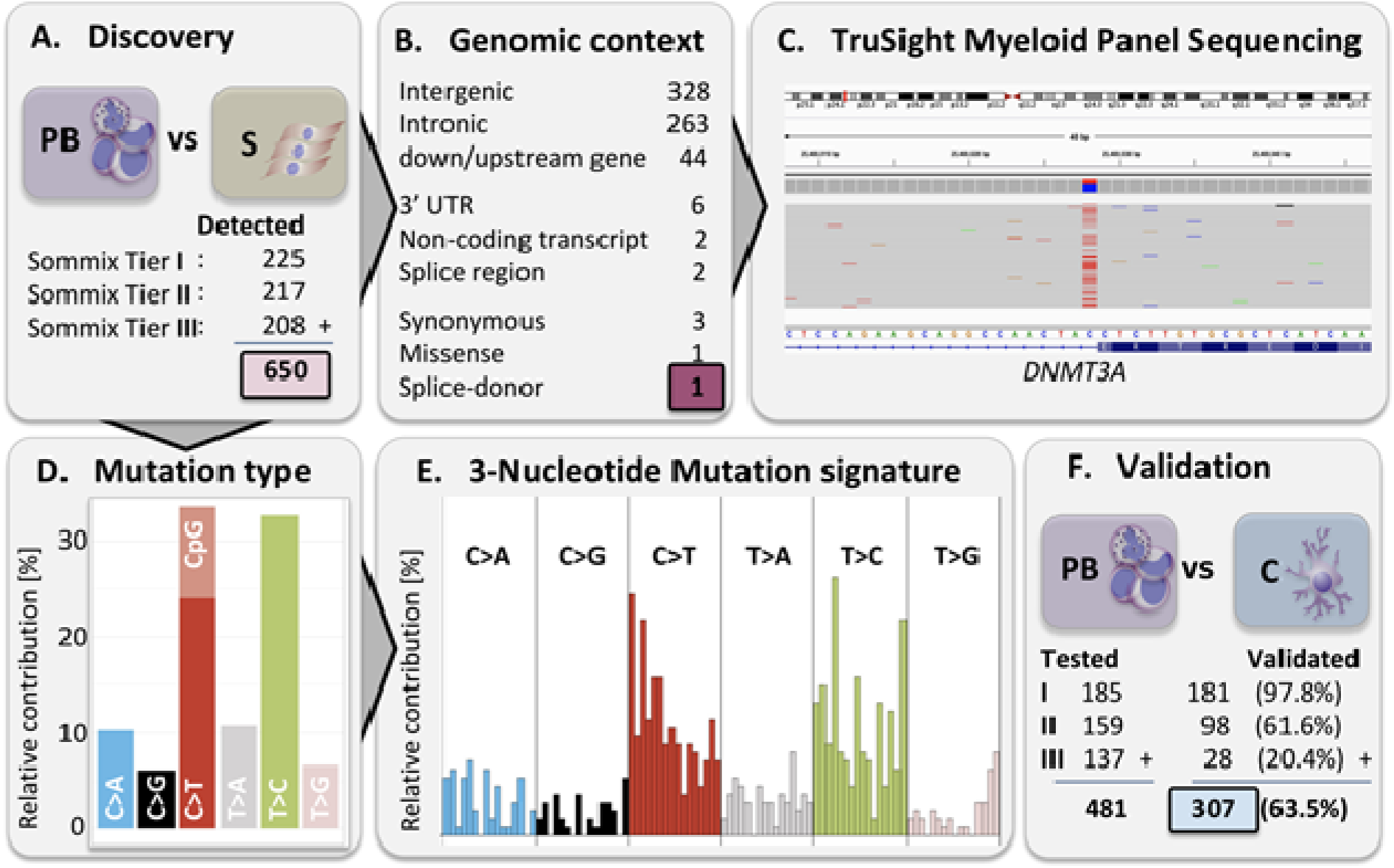
Cataloguing somatic mutations in peripheral blood drawn at age 110. [**A**] Whole genome sequencing of peripheral blood (PB) versus skin (S) and subsequent analysis with Sommix (**Methods**) identified 650 putative somatic mutations assigned to three confidence tiers (Tier I most confident). [**B**] Identified mutations mostly reside in non-coding genomic locations (UCSC’s Variant Annotation Integrator, hg19, refgene definitions) [**C**] IGV plot of the validated splice-donor site mutation in *DNMT3A* (NM_022552.4, chr2:25,469,028 C>T, c.1429+1 G>A), altering the G nucleotide of the highly conserved GT intronic sequence^52^. Amplicon sequencing performed at 961x and indicated a VAF of 37.8% for the mutant allele. [**D**] Identified mutations stratified to nucleotide changes show a high frequency of C>T and T>C changes. Part of the C>T nucleotide changes coincide with a CpG site, potentially also affecting DNA methylation. [E] 3-Nucleotide sequence context of the identified mutations exhibits a high resemblance to the clock-like mutational signatures 1 and 5 (**Supplemental Figure S2**). [**F**] Validation of the identified somatic mutations using amplicon sequencing in peripheral blood (PB) versus Cortex (C), split per confidence tier.

Next, we screened the 650 identified SNVs for potential candidate driver mutations using the mutation definitions compiled by Jaiswal *et al*.^5^ (**Methods**, definitions in **Table S2**) for 16 genes frequently mutated in the blood of healthy elderly individuals^12^. We identified a splice-donor site mutation in intron 11 of DNA (cytosine-5)-methyltransferase 3A (*DNMT3A*, NM_022552.4, chr2:25,469,028 C>T, c.1429+1 G>A). Targeted resequencing (**Methods and Supplementary Methods 4**) confirmed this variant at 961x read depth, with an estimated variant allele frequency (VAF) of 0.378 (**Figure 2C**). This suggests that 75.6% of the peripheral blood cells are derived from a single clone carrying a *DNMT3A* splice-donor mutation.

### Longitudinal analysis reveals a complex and dynamic subclonal architecture

To obtain a set of somatic mutations that can serve as genetic markers for clonal tracing, we performed validation experiments for the 650 identified putative SNVs (**Figure 2F**). Custom target re-sequencing panels were successfully designed for 474 out of 650 (72.9%) identified somatic mutations. Using brain-cortex as a control tissue, we confirm the somatic origin of 20.1% of the mutations with the lowest evidence for being somatic (Tier 3), 61.6 % of the mutation with intermediate certainty Tier 2), and 97.8% of the mutations with the highest evidence of being somatic (Tier 1) (**Figure 2**). Overall, we were able to confirm 307 (64.8%) mutations to be of somatic origin.

Amplicon sequencing in peripheral blood samples collected at ages 103 and 111 revealed that all 307 mutations identified at age 110 years were detectable (**Figure 3A**). While VAFs generally increased between ages 103 and 110 years, VAFs remained equal or decreased between ages 110 and 111 years. Nevertheless, VAFs were highly inter-correlated between timepoints (Pearson’s r_103->110_ = 0.983 and r_110->111_ = 0.988, **Figure S4**), and VAF density distributions looked highly similar across time points.

**Figure 3:**
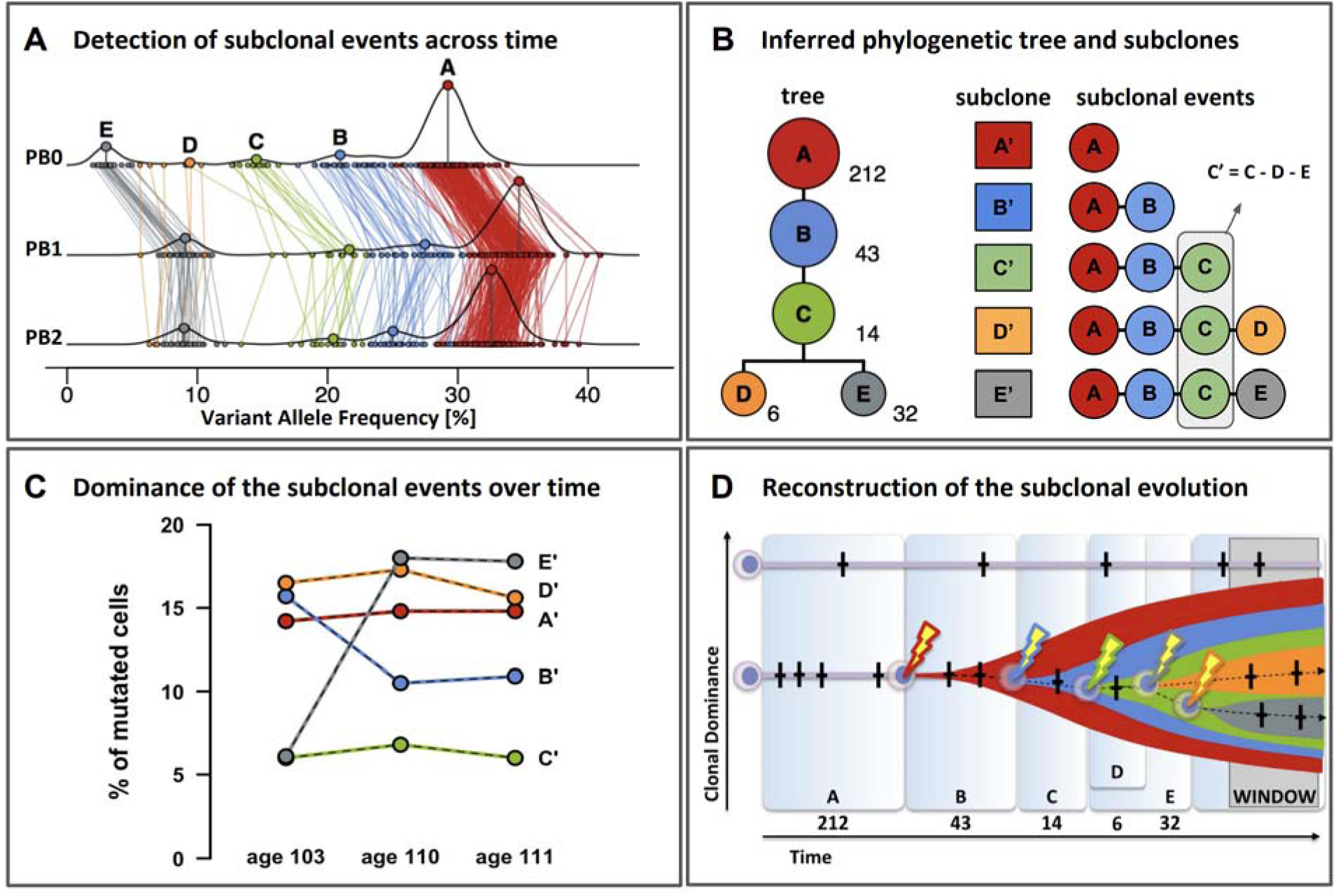
Deep sequencing of longitudinal samples reveals the clonal architecture within the peripheral blood of elderly subject with age-related clonal hematopoiesis. [**A**] Variant allele frequencies of the 307 confirmed somatic mutations at ages 103, 110 and 111. Lines connect the same mutations measured at different ages. Mutations were assigned to five independent clonal events (A-E) using SciClone^25^ and colored accordingly. [**B**] Left: Analysis with SCHISM revealed the most likely underlying subclonal architecture. The number of somatic mutations supporting each subclonal event are listed next to the clones. Right: To estimate the contributions of each subclone to peripheral blood, we need to correct for the interdependencies introduced by the shared clonal descendance. [**C**] Median VAFs after subtraction of the median VAF of the descendant clonal event indicates that clone E displays the most variation. [**D**] Reconstruction of subclonal evolution. Time frames A-E correspond to the periods in which passenger mutations (crosses) were accumulated until a clonal event driving expansion (bolt) was encountered. Widths of the time frames are roughly proportional to the number of mutations detected for each event. The y-axis reflects the relative contribution of a HSC to overall peripheral blood production. ‘WINDOW’ refers to our window of observation ranging from age 103 to 111, a 9-year period characterized by the expansion of clonal event E.

Density distributions of VAFs feature multiple peaks at all three timepoints, suggestive of an underlying subclonal architecture^14^. In accordance, a SciClone analysis^25^ (**Methods**) on VAF data from all three time points, assigned the 307 confirmed somatic mutations to five independent clonal events (A-E, **Figure 3A**). Subsequent modelling with SCHISM^26^ (**Methods**) indicated that these five clonal events most likely occurred consecutively within a single clonal lineage. This clonal lineage terminates into two independent sister-clones D and E, which were derived from a shared ancestral subclone carrying mutations associated with clonal events A-C (**Figure 3B, left**).

When analyzing the temporal changes in clonal dominance, i.e. the changes in VAFs between ages 103, 110 and 111 years, we need to take into account that the somatic variants are inherited along the subclonal architecture **(Figure 3A).** All somatic mutations in clone A are present in its clonal descendants B-E, and all somatic mutations in B are present in C-E, but not A. After adjusting for these interdependencies **(Figure 3B right, Methods)**, we observed that changes in dominance of subclones A-C are largely explained by changes in dominance of subclone E (**Figure 3C**), and notably not by subclone D. In fact, while clonal event D exhibits a near equal contribution of approximately 16.5% of the cells to peripheral blood at age 103, 110 and 111 years, clonal event E nearly tripled its clonal contribution from approximately 6.1% at age 103 to 17.9% of the peripheral blood cells at ages 110 and 111. Meanwhile, clone B becomes less dominant, as its contribution to peripheral blood decreases from approximately 15.7% at age 103 to 10.7% at ages 110 and 111.

### The founding clone and its descendants are multipotent and differentially contribute to sorted immune subsets

Next, we investigated to what extent the somatic mutations were present in the various major cell subsets of peripheral blood sampled at age 110 (PB1) and age 111 (PB2). We used the amplicon panel of 307 somatic mutations to re-sequence DNA derived from FACS-sorted immune subsets (**Figure 1**). A comparison of median VAFs between different cell subsets collected at age 110 (**Figure 4A**) indicated a significant higher clonal contribution to the myeloid branch, i.e. 87.4% of the granulocytes (G1_VAF_ = 0.437) and 77.8% of the monocytes (M1_VAF_ = 0.389), compared to total peripheral blood, i.e. 67.0% of the cells (PB1_VAF_ = 0.335). Interestingly, somatic mutations were also observed in DNA derived from cell subsets of the lymphoid lineage: ∼10.6% of the T-cells (T1_VAF_ = 0.053), and ∼7.4% of the B-cells (B1_VAF_ = 0.037) carried mutations, demonstrating both the multipotency and myeloid bias of the mutated clone.

**Figure 4:**
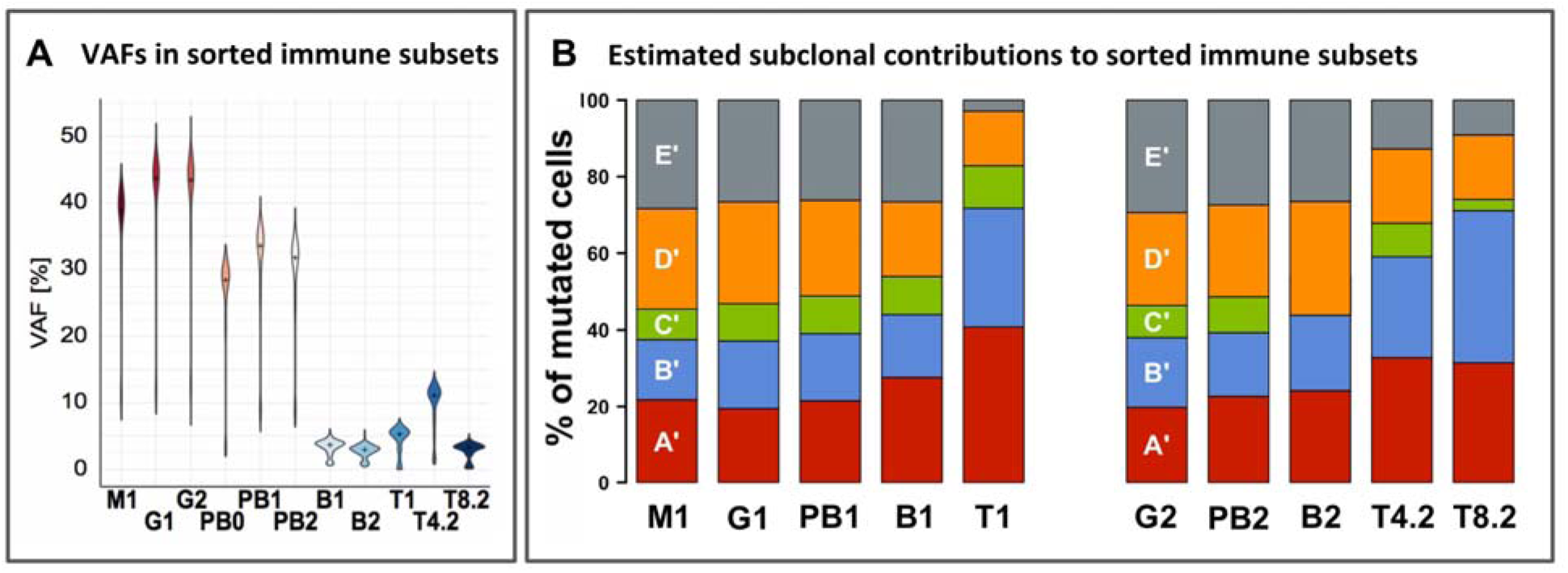
Contributions of clonal events to the major subsets in peripheral blood. [**A**] Violin plots of VAFs [%] in peripheral blood and its sorted subsets. M: Monocytes; G: Granulocytes; PB: Peripheral Blood; B: B-cells; T: T-cells; T4: CD4^+^ T-cells; T8: CD8^+^ T-cells. Numbers signify time points: 0: age 103; 1: age 110; 2: age 111. [**B**] Fraction of mutated cells per sorted cell subset derived from each subclone. Stacked bar plots per subset add up to 100%.

Re-sequencing within the blood sample collected at age 111 not only confirmed both the myeloid bias and the multipotency (**Figure 4A**), but also pointed to a bias within the lymphoid branch. Median VAFs were significantly higher in CD4^+^ T-cells (22.2% of the cells, T4.2 VAF = 0.111) compared to CD8^+^ T-cells (6.4% of the cells, T8.2_VAF_ = 0.032, *p*<0.001, Wilcoxon) and B-cells (6.0% of the cells, B2_VAF_ = 0.030, *p*<0.001, Wilcoxon). Because *DNMT3A* mutations are not restricted to myeloid leukemias^30^, but also have been reported in T-cell lymphomas^31^ and T-cell leukemias^32^, we verified the absence of T-cell malignancies by testing for clonal T-cell receptor gene recombinations^33^ (**Figure S5**).

While the founding clone (A) and its descendants (B, C, D and E) contributed to similar fractions of myeloid and peripheral blood cells, we observed a consistent deviation across the T-cell subsets. At age 110 and age 111 **(Figure 4B**, left and right), clone E contributed relatively little to sorted T-cells compared to myeloid cells or total peripheral blood cells. Interestingly, this deviation was more prominent at age 110 compared to age 111, suggesting that subclone E increased its contribution to T-cells substantially between ages 110 and 111 years. In addition, according to our calculations, subclone C did not contribute a detectable fraction to B cells at 111 years.

### W111 has an immuno-competent naïve CD4^+^ T-cell compartment

Flow cytometry analyses of peripheral blood taken at age 110 showed increased fractions of senescent CD4^+^ and CD8^+^ T-cells relative to middle aged controls, as apparent by the expression of CD57 (**Figure 5A, Figure S6**). Moreover, we observed a myeloid shift, (i.e. high myeloid to B lymphocyte ratios), particularly due to lowered B-cell levels at age 111 years (**Figure 5B**). Hence, W111’s peripheral blood shows clear signs of an aged immune system.

**Figure 5:**
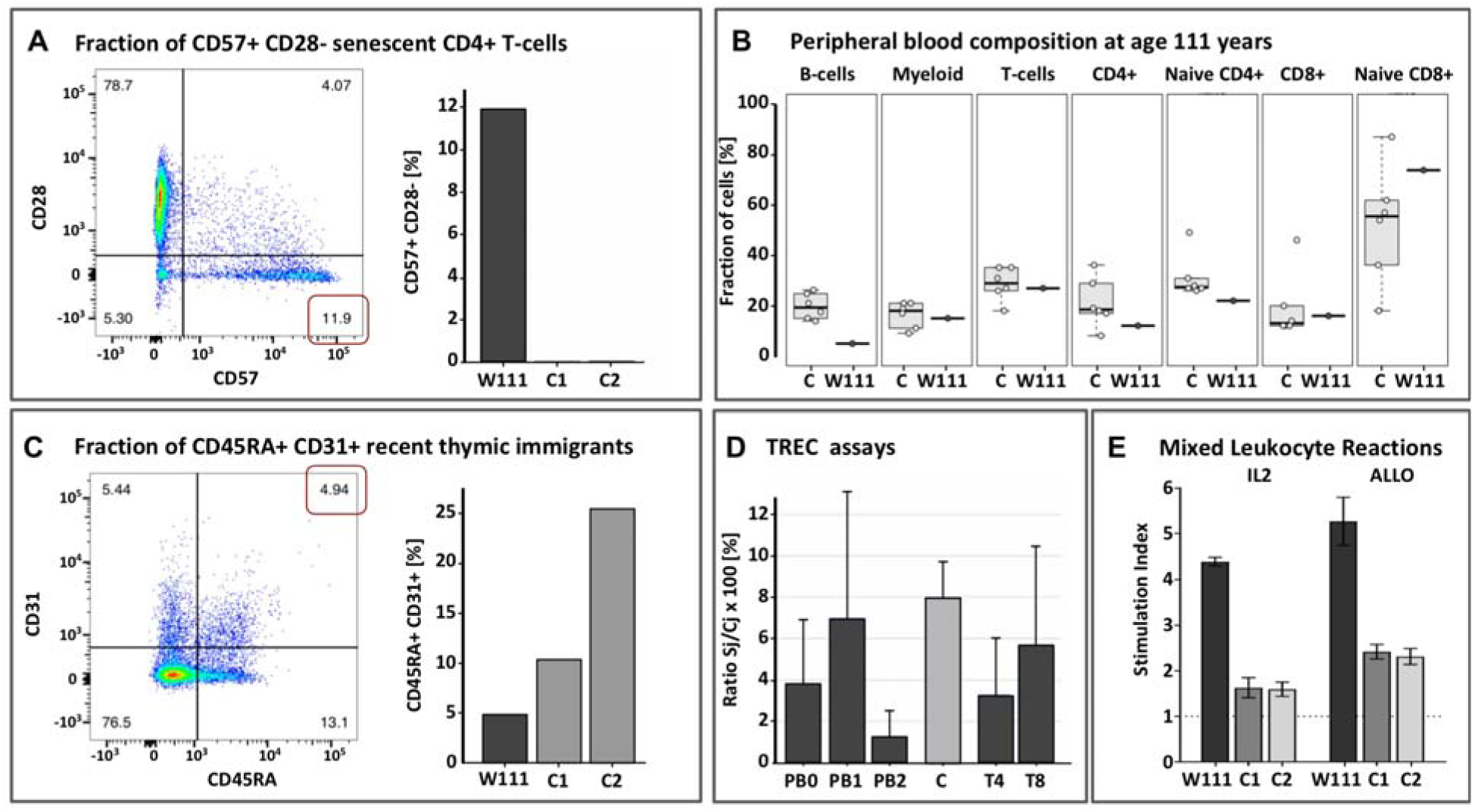
Immune characterization of W111. [**A**] Left: Sorting of CD57^+^ CD28^-^ senescent CD4^+^ T-cells at age 110 years; Right: W111 compared to middle-aged controls C1 and C2. [**B**] Proportions of sorted immune subsets (B-cells, Myeloid, T-cells, CD4^+^ T-cells, naive CD4^+^ T-cells, CD8^+^ T-cells, naive CD8^+^ T-cells) in peripheral blood of six middle-aged female controls (left) in W111 at age 110 years (right). [**C**] Left: Sorting of CD45RA^+^ CD31^+^ recent thymic immigrants at age 110 years; Right: W111 compared to middle-aged controls C1 and C2. [**D**] Percentage of TREC positive cells in material derived from W111 or a middle-aged female control C. PB: Peripheral Blood cells; T4: CD4^+^ T-cells; T8: CD8^+^ T-cells. Numbers signify time points: 0: age 103; 1: age 110; 2: age 111. [**E**] Stimulation Indices computed for IL2/TCR-dependent (IL2) and the allogenic (ALLO) assay of cultured T-cells of W111 versus two middle-aged female controls C1 and C2.

In contrast, we also observed that, at ages 110 and 111 years, the fraction of naive CD4^+^ T-cells was only slightly decreased and that the fraction of naive CD8^+^ T-cells was comparable relative to middle aged controls (**Figure 5B, Figures S7**). Moreover, at age 110 years, we observed that nearly 5% of the CD4^+^ T-cells expressed both CD45RA and CD31, indicative of recent thymic emigrants (**Figure 5C left, Figure S6**). Although this level is much lower than observed in the middle-aged controls **(Figure 5C right)**, this suggests that there are still recent thymic emigrants in W111’s peripheral blood^34,35^. In accordance, the assessment of the replication history of T-cells by T-cell receptor excision circles (TRECs) assays indicated that both CD4^+^ and CD8^+^ T-cells had undergone a similar number of divisions compared to middle-aged healthy controls (**Methods, Figure 5D**), with TREC contents of 3-6%^20^.

In parallel, we investigated the capacity of W111’s T-cells to mount an immune response. Flow cytometry analyses of cells collected at age 110 indicated that both the CD4^+^ and CD8^+^ T-cell subsets contained considerable fractions of *in vivo* activated cells, evidenced by their high CD25 expression and CD69 expression (**Figure S7**). This suggests ongoing or recent T-cell directed immune responses. To further validate this observation, we performed two types of *in vitro* proliferation assays (**Methods**). In both the IL2/TCR-dependent and the allogeneic mixed-lymphocyte assays, T-cells collected from W111 outperformed those taken from middle-aged controls on a per cell basis (**Figure 5E**).

## Discussion

We explored the properties of extreme age-related clonal hematopoiesis (ARCH) and consequences on immuno-competence in an immuno-hematologically normal female, whose blood was longitudinally sampled at age 103, 110 and 111 years. We observed that >75% of her peripheral blood was derived from a single *DNMT3A*-mutated hematopoietic stem cell (HSC) clone. We observed that this HSC clone had a complex subclonal architecture, in which one founding clone A produced descendant subclones that successively expanded over time. Clonal evolution most likely started prior to our study and subclonal competition between sister-lineages was still ongoing during the 9-year period of observation, as evidenced by the increased expansion of new subclone E. Our results further indicated that while the large majority of clonal output was myeloid biased, the clone also contributed to lymphopoiesis, in particular to CD4+ T-cells. Surprisingly, we observed that, despite extreme ARCH, this extremely aged individual was able to mount a functional T-cell response, evidenced by *in vitro* proliferation and T-cell receptor excision circle assays.

ARCH can either be established by identification of a candidate driver mutation, typically residing in *DNMT3A, TET2* or *ASXL1*^2,3,13^, or by identification of a disproportionately large number of somatic mutations, which are presumed to be passenger mutations^1,3^. With 650 identified somatic mutations, including a mutation in *DNMT3A*, the ARCH identified complied with both criteria. The identified mutation in *DNMT3A* disrupts a canonical splice-donor site and was previously reported in patients with hematopoietic or lymphoid malignancies (COSM5945645)^36-38^. Since this mutation was observed in the founding clone, it is well possible that this somatic mutation drove the observed clonal expansion. Importantly, examination of peripheral blood revealed neither cytopenias nor dysplastic morphologies, and sequencing-based diagnostic tests for suspect myeloid or lymphoid leukemias were negative^33^. Collectively, these findings fit the inclusion criteria for Clonal Hematopoiesis of indeterminate Potential (ChiP)^7^, also referred to as ARCH^39^, to describe immuno-hematologically normal elderly individuals carrying a pre-leukemic mutation with unclear clinical implications.

While the initiating clonal expansion may be attributable to a *DNMT3A* splice donor mutation, no additional somatic mutations were identified that could explain any of the successive subclonal expansions. This result might be explained by undetected or incomplete knowledge of driver mutations^1^, but could also point to non-genetic mechanisms, e.g. so-called ‘epi-mutations’^40^. An explanation for this may lie in the fact that *DNMT3A* is a key epigenetic regulator of myelopoiesis^41^ and responsible for the maintenance of DNA methylation in HSCs^42^. A mutation in *DNMT3A* may lead to a gradual loss of epigenetic control, allowing ‘epimutations’ to arise, which may iteratively improve the replicatory fitness of an HSC^43^. Alternatively, the effect of the *DNMT3A* mutation may lead to an enhanced HSC-proliferation upon bone marrow stress, such as inflammation^44^ or environmental stimuli^45^. In such a scenario, the clonal architecture would represent the history of re-activation of otherwise quiescent HSCs^46^.

In line with previous reports in individuals with ARCH^47,48^, we observed that the *DNMT3A*-mutated stem cell contributed to the large majority of the myeloid cells (78-87%) and only to a small proportion of the T-cells (11%) and B-cells (6-7%). Moreover, the mutated stem cell contributed to a significantly larger proportion of CD4+ T-cells (22%) than to CD8+ T-cells (6%). We also observed differences between the subclonal contributions to specific cell subsets. Specifically, we noted that subclones A and B generated a disproportionally high fraction of T-cells, while subclone E generated a disproportionally low fraction. Collectively, These observations could be explained by a progressively myeloid bias in the offspring generated by the newer subclones. An alternative explanation, as previously proposed by Buscarlet *et al*.^47^, may be found in the differential longevity of specific cell subsets. T-cells, in particular, are known to live years to tens of years, to uphold long-term immunity for antigens against which they were raised. Therefore, T-cells generated by older clones, possibly years prior to sampling, may lead to a relative increased contribution of older clones to T-cells. Interestingly, we observed that between age 110 and age 111, the contribution of subclone E changed to become more comparable with myeloid cells and peripheral blood cells. In addition, the contribution of subclone C to B cells is lost between age 110 and age 111. Together, these changes suggest strong and ongoing dynamics in the subclonal contribution to specific cell subsets.

Ageing of the T-cell compartment, i.e. immune senescence has been postulated as a major factor underlying a reduced life expectancy^49,50^. The thymus gradually ceases to function after puberty and is fully involuted and inactive at advanced ages. The concentration T-cell receptor excision circles (TRECs) in peripheral blood, indicative of T-cell proliferation, decreases correspondingly and is often undetectable after 85 years^20^. Given this background, our observations in W111 are remarkable. TREC contents in W111’s peripheral blood and sorted CD4^+^ and CD8^+^ T-cells at age 110 and 111 years were comparable to that of middle-aged healthy controls. Additionally, in *in vitro* proliferation assays, we observed that W111 had preserved the capability of mounting vigorous naive responses which suggests that W111 still disposes of functional T-cell immunity. In parallel, we observe a marked contribution of the stem cell clone to the CD4^+^ T-cell subset which raises the possibility that the exceptional functional T-cell immunity could be attributed to the lymphoid progeny of the *DNMT3A* mutated stem cell. In aggregate, this leads us to speculate that maintained T-cell immunity, as a prerequisite for exceptional longevity, is partly derived from a *DNMT3A*-dependent clonal expansion. Interestingly, the suggestion that a clonally expanded CD4^+^ T-cell subset may serve as a potential hallmark of the supercentenarian immune system is in line with recent findings by Hashimoto *et al*.^51^ who observed clonally expanded CD4^+^ T-cells in 7 supercentenarians. While Hashimoto *et al.* attributed this clonal T-cell expansion to a sustained antigenic stimulation, our findings suggest ARCH as a competing explanation.

We acknowledge that this study describes observations in only a single healthy supercentenarian which precludes any inference of causality between the co-occurrence of an extreme *DNMT3A*-associated ARCH and an unexpectedly functional T-cell immunity. Nevertheless, the presented findings warrant future research into our findings which suggest that a functional T-cell immunity is essential for achieving an exceptional age in good health.

## Acknowledgements

The authors thank W111 for her enthusiasm in study participation, for allowing repeated sample collection and agreeing to post mortem brain donation. Our acknowledgements also go out to her family, for their continuous support of this study. The 100-plus Study was supported by Stichting Alzheimer Nederland (WE09.2014-03), Stichting Diorapthe (VSM.14.04.14.02), Stichting VUmc Fonds and the Horstingstuit Foundation. The Leiden Longevity Study has received funding from the European Union’s Seventh Framework Programme (FP7/2007-2011) under grant agreement number 259679. This study was supported by the Netherlands Consortium for Healthy Ageing (grant 050-060-810), in the framework of the Netherlands Genomics Initiative, Netherlands Organization for Scientific Research (NWO); by BBMRI-NL, a Research Infrastructure financed by the Dutch government (NWO 184.021.007). EvdA is funded by a personal grant of the Dutch Research Council (NWO; VENI: 09150161810095)

## Author contributions

Samples were collected by: H.H. and P.E.S.P. Experiments were performed by: M.H.B., T.N., T.B., M.E.J., and Q.W. Results were analyzed and figures were made by: E.A., S.M., and M.H. Experimental design was supported by: F.B., D.J., F.J.T.S. M.J.T.R. and H.H. H.H. conceived and financed the study. The manuscript was written by E.A. and H.H.

---

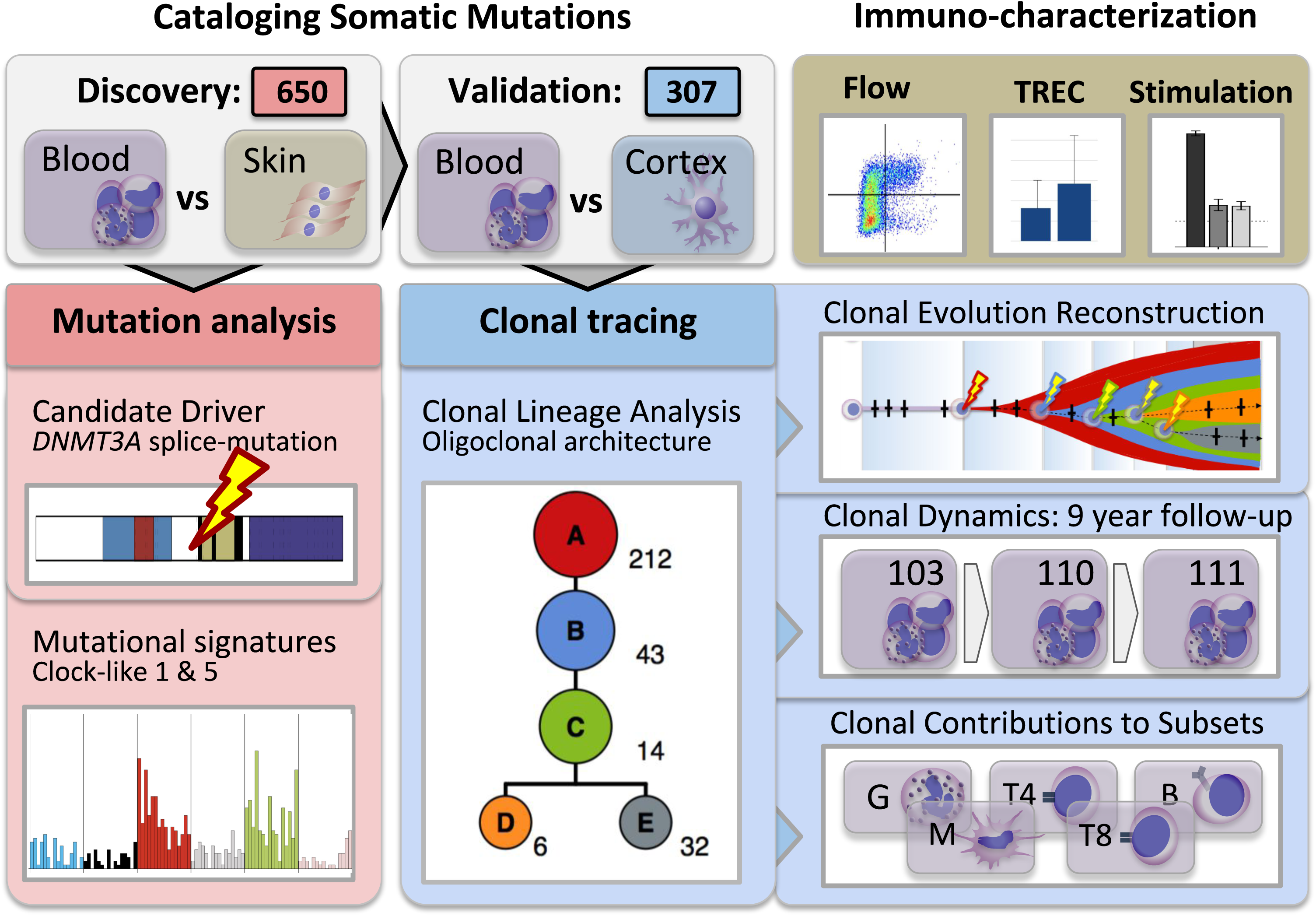

