## Supplemental Material for "Dynamic Clonal Hematopoiesis and Functional T-cell Immunity in a Super-centenarian"

to

### TABLE OF CONTENTS


**SUPPLEMENTARY METHODS** 3

Methods S01: TruSight Myeloid Panel - MiSeq 3
Methods S02: Whole genome sequencing 3
Methods S03: Sommix 3
Methods S04: Amplicon re-sequencing 7
Methods S05: Identification of candidate driver mutations 8
Methods S06: Immuno-phenotyping by flow cytometry 8
Methods S07: T-cell Receptor Excision Circle (TREC) assays 8
Methods S08: Mixed Leukocyte Reactions (MLR) 9

**SUPPLEMENTARY FIGURES** 10

Figure S01: Rainfall plot of 650 putative SNVs 10
Figure S02: Tri-nucleotide context of 650 putative SNVs 11
Figure S03: Mutational signatures contributing to 650 putative SNVs 12
Figure S04: Pairwise scatter plots of 307 SNVs at PB0, PB1 & PB2 13
Figure S05: Testing for suspect clonal BCR/TCR gene recombinations 16
Figure S06: T-cell characterization at age 110 by flow cytometry 20
Figure S07: T-cell characterization at age 111 by flow cytometry 29

**SUPPLEMENTARY TABLES** 30

Table S1: 650 Putative Somatic Mutations 30
Table S2: Variant Allele Fractions of 307 Validated Somatic Mutations 44
Table S3: Candidate driver definitions 51

**REFERENCES** 53

References 53

### Supplementary Methods

#### SM1: TruSight Myeloid Panel - MiSeq

54 Genes recurrently mutated in myeloid malignancies were re-sequenced using Illumina's TruSight Myeloid Panel (v1.4) on a MiSeq. Obtained data was analysed using Illumina's Isis (v2.6.2.3). Assayed genes included: *ABL1, CEBPA, HRAS, MYD88, SF3B1, ASXL1, CSF3R, IDH1, NOTCH1, SMC1A, ATRX, CUX1, IDH2, NPM1, SMC3, BCOR, DNMT3A, IKZF1, NRAS, SRSF2, BCORL1, ETV6/TEL, JAK2, PDGFRA, STAG2, BRAF, EZH2, JAK3, PHF6, TET2, CALR, FBXW7, KDM6A, PTEN, TP53, CBL, FLT3, KIT, PTPN11, U2AF1, CBLB, GATA1, KRAS, RAD21, WT1, CBLC, GATA2, MLL, RUNX1, ZRSR2, CDKN2A, GNAS, MPL, SETBP1.*

#### SM2: Whole genome sequencing

Whole genome sequencing was performed in peripheral blood and skin obtained at age 110 years. Library preparations were performed according to the Fragment Library Preparation 5500 series SOLiD protocol (4460960A) and executed using an automated routine on the Biomek FXp workstation. The resulting library was prepared for sequencing according to SOLiD protocols EZB_Emuls_gsg_4441486E, EZB_AMP_GSG_4443494E, and EZB_enrich_gsg_4443496E. Sequencing was conducted using the SOLiD 5500XL with paired end sequences, 75bp forward en 35bp reverse, according to the 5500 user guide 4456991L protocol. Resulting data was aligned with Lifescope (v2.5), after which variants were called using GATK (v2.7.5), involving the following steps: Local read depths were determined using GATK's DepthOfCoverage; local realignment around known indels was performed calling GATK's RealignerTargetCreator and subsequently IndelRealigner. Mate information was then recovered using Picard Tools (v1.105). Base call recalibration was performed by running BaseRecalibrator after which variants were called using HaplotypeCaller. Variants were annotated using KGGSeq (v0.4).

#### SM3: Sommix

##### Rationale and Design

To detect somatic mutations in blood using skin as a control we had to correct for the contamination of skin cells with blood. For this purpose, we developed Sommix. Input data for Sommix are mapped reads from a blood and a skin sample as well as the expected contamination fraction *c*. The output of the algorithm is the most probable combination of genotypes for the blood and skin samples at every genomic position.

To narrow the search space, we did not assess the whole genome with Sommix, but only the positions where GATK found a possible indication of a variant in either blood or skin. BAM files were realigned and recalibrated, after which the GATK Haplotypecaller (version 3.1) was used to obtain variant calls. A low confidence threshold of 10 for emitting a variant was used, resulting in a total of 4,554,470 possible variant sites. All these sites, irrespective of their call status or quality, were reinvestigated through Sommix.

We denote $\boldsymbol{G} = \{\mathrm{AA, AC, AG, AT, CC, CG, CT, GG, GT, TT}\}$ as the set of all possible genotypes and $G_b, G_s \in \boldsymbol{G}$ as the genotype of the blood and the skin sample respectively in a given genomic position. Sommix identifies the most probable pair of $G_b, G_s$ out of the 100 possible combinations. If there are $N_b$ reads covering the specific position in blood and $N_s$ reads in skin, the probability of each genotype pair given the available reads (*D*) is given by:

$$P(G_b, G_s|D) = \frac{P(G_b, G_s) \cdot P(D|G_b, G_s)}{P(D)} = \frac{P(G_b, G_s) \cdot \prod_{i=1}^{N_b} P(r_i|G_b, G_s) \cdot \prod_{j=1}^{N_s} P(r_j|G_b, G_s)}{P(D)} \quad (1)$$

where $P(G_b, G_s)$ is the prior probability of the genotype pair $G_b, G_s$ and $P(D)$ is a normalizing factor. Since $P(D)$ is the same for all 100 genotype combinations, it does not influence the final result. Therefore, Sommix only computes the numerator of Eq. 1.

#### Computation of Priors

Let $g$, $s$ be the prior probability of observing a germ line and a somatic mutation respectively in any position.
Let R be the reference base and $V_1$, $V_2$, $V_3$ each one of the remaining three bases.
The prior for a genotype pair is

$$P(G_b, G_s) = P(G_s)\, P(G_b|G_s) \quad (2)$$

where the prior of a germ line mutation is set to:

$$P(G_s) = \begin{cases} 1 - g, & G_s = RR \\ \frac{2g}{3}, & G_s = RV_i \\ \frac{g}{3}, & G_s = V_iV_j \end{cases} \quad (3)$$

For loci in dbSNP (build 141), $g$ is set to 0.2, while for locations not in dbSNP, it is set to 0.001.
The prior of a somatic mutation is set to:

$$P(G_b|G_s) = \begin{cases} 1 - s, & G_b = G_s \\ s, & if\ G_b\ and\ G_s\ have\ 1\ allele\ in\ common \\ 0, & otherwise \end{cases} \quad (4)$$

The prior $s$ is set to 1e-07 (corresponding to an expectation of approximately 300 positions in the whole genome with a somatic mutation).

#### Somatic genotype fraction estimate

Sommix calculates for every genotype pair $G_b, G_s$ the fraction to which a somatic genotype is present in a position, which we denote as $V(G_b, G_s)$. This value is the weighted average of the estimate of this fraction in blood and skin. If $G_b = G_s$, then $V(G_b, G_s) = 0$.

For a heterozygous somatic mutation:

$$V^{blood}(G_b, G_s) = 2\frac{\#reads\ with\ somatic\ allele\ in\ blood}{\#reads\ with\ reference\ allele\ in\ blood + \#reads\ with\ somatic\ allele\ in\ blood} \quad (5)$$

$$V^{skin}(G_b, G_s) = 2\frac{\#reads\ with\ somatic\ allele\ in\ skin}{\#reads\ with\ reference\ allele\ in\ skin + \#reads\ with\ somatic\ allele\ in\ skins} \quad (6)$$

Other more complex somatic events, such as homozygous somatic mutations and/or the presence of germ line mutations are handled similarly.

To obtain comparable somatic genotype fractions, we convert the estimate $V^{skin}(G_b, G_s)$ to $V^{blood*skin}(G_b, G_s)$ through use of the expected amount of contamination of blood cells in the skin sample:

$$V^{blood*skin}(G_b, G_s) = \frac{V^{skin}(G_b, G_s)}{c} \quad (7)$$

Then the weighted average is calculated as follows:

$$V(G_b, G_s) = \frac{\#reads\ in\ skin \cdot c \cdot V^{blood*skin}(G_b, G_s) + \#reads\ in\ blood \cdot V^{blood}(G_b, G_s)}{\#reads\ in\ skin \cdot c + \#reads\ in\ blood} \quad (8)$$

This reduces the weight of the skin sample by the expected contamination fraction.

#### Likelihood of a read given a genotype pair

The likelihood of each base given a genotype depends on whether it is present in the genotype and on whether it is a reference base or not. It also depends on the base-calling quality (*BQ*) and mapping quality (*MQ*) in the position in question for a given read. Let $e_{bc}$ be the probability that the base called in a certain position in a given read is erroneous and $e_m$ the probability that a mapping error has occurred in the read.

Base *b* given the genotype *G* = *bb* can be observed in the following ways:

- The sequenced base is *b* and the base-calling and mapping are done correctly.
- The sequenced base is any of the 4 bases and a base-calling error has occurred.
- The sequenced base is *b*, the correct base has been called, but a mapping error has occurred. In that case the likelihood to observe a certain *b* depends on the reference sequence, as an incorrect mapping can only occur for reads that are very similar to the reference sequence at the mapped location. We assume, therefore, that an incorrectly mapped read at each position follows the reference with 0.95 probability, and conversely, includes a non-reference base at 0.05 probability.

If *G* is heterozygous for *b* (*G* = *bx*), then the ways to observe base *b* are similar, with one difference: In the case where both the base-calling and mapping are correct, there is still 0.5 probability of observing each of the two bases.

Finally, if *G* does not contain base *b*, then observing *b* can be either to a base calling or a mapping error. In the case of correct base calling and a mapping error, we still expect a higher chance of observing base *b*, if it is the reference base.

The above is summed up in Eq. 9, where *β* = 0.95 if *b* is the reference base and 0.05 if not.

$$P(b|G) = \begin{cases} \frac{e_{bc}}{4} + (1 - e_{bc})(1 - e_m + \beta e_m), & G = bb \\ \frac{e_{bc}}{4} + (1 - e_{bc})\left(\frac{1 - e_m}{2} + \beta e_m\right), & G = bx \\ \frac{e_{bc}}{4} + (1 - e_{bc})\beta e_m, & b \notin G \end{cases} \quad (9)$$

The probability of observing base $b^s$ in skin is:

$$P(r_i = b^s | G_b, G_s) = (1 - V(G_b, G_s) \cdot c) \cdot P(b^s|G_s) + V(G_b, G_s) \cdot c \cdot P(b^s|G_b) \quad (10)$$

The probability of observing base $b^b$ in blood is:

$$P(r_i = b^b | G_b, G_s) = \left(1 - V(G_b, G_s)\right) \cdot P(b^b|G_s) + V(G_b, G_s) \cdot P(b^b|G_b) \quad (11)$$

#### Heterozygosity Correction

In case of germ line heterozygosity, the expected allele frequency is 0.5 for both alleles. However, in practice, amplification biases and mapping affinities frequently lead to quite different allele frequencies. Not including this in the scoring framework would lead to false positive calls, as germ line heterozygous loci with deviating allele frequency patterns could also be interpreted as a mixture of a germ line and a somatic genotype.

To correct for this, we also estimate the heterozygosity allele frequency for germ line variants directly based on the reads.

If no somatic event occurred, then the expected germ line heterozygous VAF in skin is the average of the calculated VAFs in blood and skin weighted by the number of reads in each sample.

In case of a somatic event, we subtract the part of the reads that can be attributed to the somatic genotype fraction, using the following correction factors:

$$cor_b = 1 - \frac{V(G_b, G_s)}{1 + V(G_b, G_s)} \quad (12)$$

$$cor_s = 1 - \frac{V(G_b, G_s) \cdot c}{1 + V(G_b, G_s) \cdot c} \quad (13)$$

If the somatic genotype is homozygous (and the germ line heterozygous), the estimated heterozygous allele frequency is now estimated as:

$$VAF_{corrected} = \frac{cor_s \cdot \#reads\ with\ som.allele\ in\ skin + cor_b \cdot \#reads\ with\ som.allele\ in\ skin}{cor_s \cdot \#reads\ with\ som.allele\ in\ skin + cor_b \cdot \#reads\ with\ som.allele\ in\ blood + \#reads\ with\ ref.allele\ in\ skin + \#reads\ with\ ref.allele\ in\ blood} \quad (14)$$

More complex cases that involve more than one somatic mutation are handled similarly.

Based on these corrections, in the cases when the germ line genotype is heterozygous, the likelihood of each of the two bases given the genotype needs to be revisited. More specifically, we assumed earlier that in a heterozygous site without base-calling and mapping errors there is 0.5 chance that each of the bases might be observed. We showed here that this is not the case, so this likelihood equation for the germ line was modified as follows:

$$(b|G) = \frac{e_{bc}}{4} + (1 - e_{bc})((1 - e_m) \cdot VAF_{corrected} + \beta e_m), \quad G = bx\ in\ germline \quad (15)$$

#### Mappability correction

To identify locations that might have high mismapping ratios, we downloaded the 50bp mappability track from UCSC (Raney et al. 2011). The mappability for each locus is encoded as 1.0 divided by the number of 50bp genome sections that, with up to two mismatches, can map to this locus. We include this in the mapping error probability calculations, by adapting this probability as follows:

$$e_m = \max(e_m, 1.0 - mappability) \quad (16)$$

#### Depth correction

Alternatively, an anomalously high read depth can also identify mismapped reads. This might be an indication of repeat regions that are mapped to the same locus on the reference genome. Given an expected read depth of 80x, we calculated a mapping quality based on depth as follows:

$$e_{depth} = 1 - \frac{depth \cdot 2.0 \cdot 1.5}{depth\ in\ skin + depth\ in\ blood} \quad (17)$$

Similarly, as for the mappability correction, we included this in the mapping quality:

$$e_m = \max(e_m, 1.0 - \textit{mappability}) \qquad (18)$$

##### Sommix Scores

The optimal genotypes pair is the one that maximizes $\log(P(G_b, G_s|D))$. The Sommix score is calculated as the logarithm of the likelihood ratio between the two most probable genotypes. This score is 0 when two genotypes pairs are equally likely and infinity when only one genotypes pair has non-zero likelihood (e.g. if there are only perfect reads for the base A in both blood and skin in a certain locus, only the pair AA-AA would have non-zero likelihood). Loci with a homozygous reference genotype in skin and a heterozygous genotype in blood that had a positive Sommix score were selected for further analysis.

##### Additional filtering

To obtain a high quality set of calls, three additional filtering steps were performed:

*Strand Bias*

To identify positions with high strand bias, we used the hypergeometric test to test whether strand bias was present. Every cell of the contingency table contains the number of reads that supports the corresponding allele and map to the corresponding strand in both blood and skin. We used a conservative threshold, were loci with a p-value lower than 0.1 were excluded.

*Tri-allelic variants*

Conservatively, loci with 3 alleles were not considered, as these are more likely to constitute mismapping reads rather than a true complex somatic event.

*Concordance with GATK's called variant types*

Putative blood mutations were excluded from further analysis in positions where Sommix detected a somatic single nucleotide substitution (SNV) whereas GATK detected a somatic indel.

###### Division into Confidence Tiers

Application of Sommix yielded 929 bi-allelic somatic single nucleotide mutations in blood. Subsequent filtering removed 262 with a strand bias, and 17 with a discordant variant type called by GATK. The remaining 650 putative single nucleotide somatic mutations were classified into three nearly equal-sized levels of confidence: Tier 1 Sommix score > 1.7; Tier 2 Sommix score > 0.65; Tier 3 Sommix score > 0.

#### SM4: Amplicon re-sequencing

Amplicon re-sequencing was performed to 1] validate the putative somatic mutations identified by whole genome SOLiD sequencing in peripheral blood and skin 2] to assess and compare the VAFs of validated somatic SNVs in peripheral blood at age 103, 110 and 111 3] to assess and compare the VAFs of validated somatic SNVs in sorted immune subsets.

For this purpose, amplicon design software (Ion AmpliSeq Designer V4.2) was employed to automatically design 100bp insert size amplicons covering the putative somatic variants. Ampliseq libraries were prepared according to the Ion AmpliSeq™ Library Preparation protocol (MAN0006735, Revision A.0). Sequencing was performed according to Ion PI™ IC 200 Kit protocol (MAN0010078, Revision B.0). Obtained data was analysed with Torrent Suite (V4.4.2) to verify whether the targeted genomic areas were sufficiently covered, using the following parameters:

```
BeadFind Args: justBeadFind –beadfind-minlivesnr 3 –region-size=216,224
                 –total-timeout 600
Analysis Args: Analysis –from-beadfind –clonal-filter-bkgmodel true
                 –region-size=216,224 –bkg-bfmask-update false
```

```
                        –gpuWorkLoad 1 –total-timeout 600
                        –gopt /opt/ion/config/goptp1.1.17ampliseqexome.param.json
Pre-BaseCaller Args for calibration: BaseCaller –barcode-filter 0.01
                        –barcode-filter-minreads 10 –keypass-filter on
                        –phasing-residual-filter=2.0 –num-unfiltered 1000
                        –max-phasing-levels 2
Calibration Args: calibrate –skipDroop
BaseCaller Args: BaseCaller –barcode-filter 0.01
                        –barcode-filter-minreads 10 –keypass-filter on
                        –phasing-residual-filter=2.0 –num-unfiltered 1000
                        –barcode-filter-postpone 1
Alignment Args: stage1 map4
IonStats Args:
Analysis Parameters: default
```

#### SM5: Identification of candidate driver mutations

WGS data was also analysed for the presence of candidate driver mutations in 16 genes previously reported to be recurrently mutated in the blood of apparently healthy elderly individuals: *DNMT3A*, *TET2*, *ASXL1*, *TP53*, *JAK2*, *SF3B1*, *GNB1*, *CBL*, *SRSF2*, *GNAS*, *BRCC3*, *CREBBP*, *NRA S*, *RAD21*, *U2AF1*, *PPM1D*[1-4]. For this purpose, SNPs and short indels called by GATK Haplotypecaller were consecutively filtered according to the following five criteria. *i*) Positioned within or near, max 5 nucleotides, of the coding sequence of a transcript annotated to the aforementioned genes (hg19, UCSC refgene definitions, UCSC's Variant Annotation Integrator (https://genome.ucsc.edu/cgi-bin/hgVai). *ii*) A minimal read depth of the mutant allele of 3 and 6 for respectively SNVs and indels. *iii*) A protein altering predicted impact (UCSC Variant Annotation Integrator, NCBI Refseq curated subset). *iv*) Adhering to the gene specific mutational profiles, as previously compiled[5] (**Table S3**). *v*) Called in blood, but not in brain. A *DNMT3A* splice mutation was identified and confirmed by resequencing the peripheral sample blood drawn at 110 years and 3 months using the Illumina TruSightMyeloid panel and MiSeq instrument at 961x and visualized in Integrative Genome Viewer (IGV, Broad Institute, version 2.4.9).

#### SM6: Immuno-phenotyping by flow cytometry

PBMCs were incubated with a titrated cocktail of antibodies for 30 minutes on ice in the dark. Cells were subsequently washed with FACS buffer (PBS supplemented with 2% FCS and Na-azide) prior to flow cytometry analysis using FACS Fortessa-X20 (BD Biosciences). The employed antibody mix consisted of: FITC-anti-CD57 (clone HCD57); PE-anti-CD28 (clone CD28.2); APC-Cy7-anti-CD4 (clone RPA-T4); PE-Cy7-anti-CD197 (CCR7, clone 3D12); Pacific Blue-anti-CD27 (clone M-T271), BV650-anti-CD8 (clone SK1), all were from BD Pharmingen (USA). Alexa Fluor 700-anti-CD45RA (clone HI100); BV605-anti-CD31 (clone WM59) were from Biolegend (San Diego, USA). Phenotypic data were analyzed using FlowJo software (TreeStar).

#### SM7: T-cell Receptor Excision Circle (TREC) assays

T-cell receptor excision circle (TREC) assays are based on the rearrangement of the T-cell receptor occurring early during T-cell development[6]. This rearrangement results in the formation of a coding joint (CJ) which remains stably present in the genomic DNA, and a signal joint (SJ) on the corresponding excision circle. As with every cell division, SJ is diluted, while CJ is stably maintained in the genomic DNA, it is possible to derive the number of divisions the assayed cells have undergone from the difference between CJ and SJ[7]. TRECs analyses were essentially performed as previously described[8]. In short, DNA was extracted from peripheral blood or sorted T-cell subsets, after which two parallel real-time PCRs are performed that quantify the signal joint (SJ) or the coding joint (Cj) relative to an internal control of Albumin.

#### SM8: Mixed Leukocyte Reactions (MLR)

Antigen-dependent T-cell proliferation capacity was determined using a mixed leukocyte reaction (MLR). PBMC's from test samples were cultured with irradiated HLA-mismatched PBMCs (3000 rad). To assess the general capacity of T-cells to proliferate, PBMCs were stimulated with Interleukin-2 (IL-2, 25 IU/ml). Cells were incubated in triplo in the 96-well round bottom plates in IMDM medium conditioned with Glutamine, Pen/Strep and 10% inactivated Human Serum (HS, Sanquin, Amsterdam) for 5 days. Radioactive thymidine (2 µCi/ml) was added in the last 18 hrs of the culture to enable incorporation in the DNA of proliferating cells and the radioactivity in DNA recovered from the cells was measured using a scintillation beta-counter. Obtained data (counts per minute, CPM) were used to calculate stimulation index by dividing the CPM values of the test samples by the CPM of test PBMC cultured in medium alone.

### Supplementary Figures

#### Figure S01: Rainfall plot of 650 putative SNVs

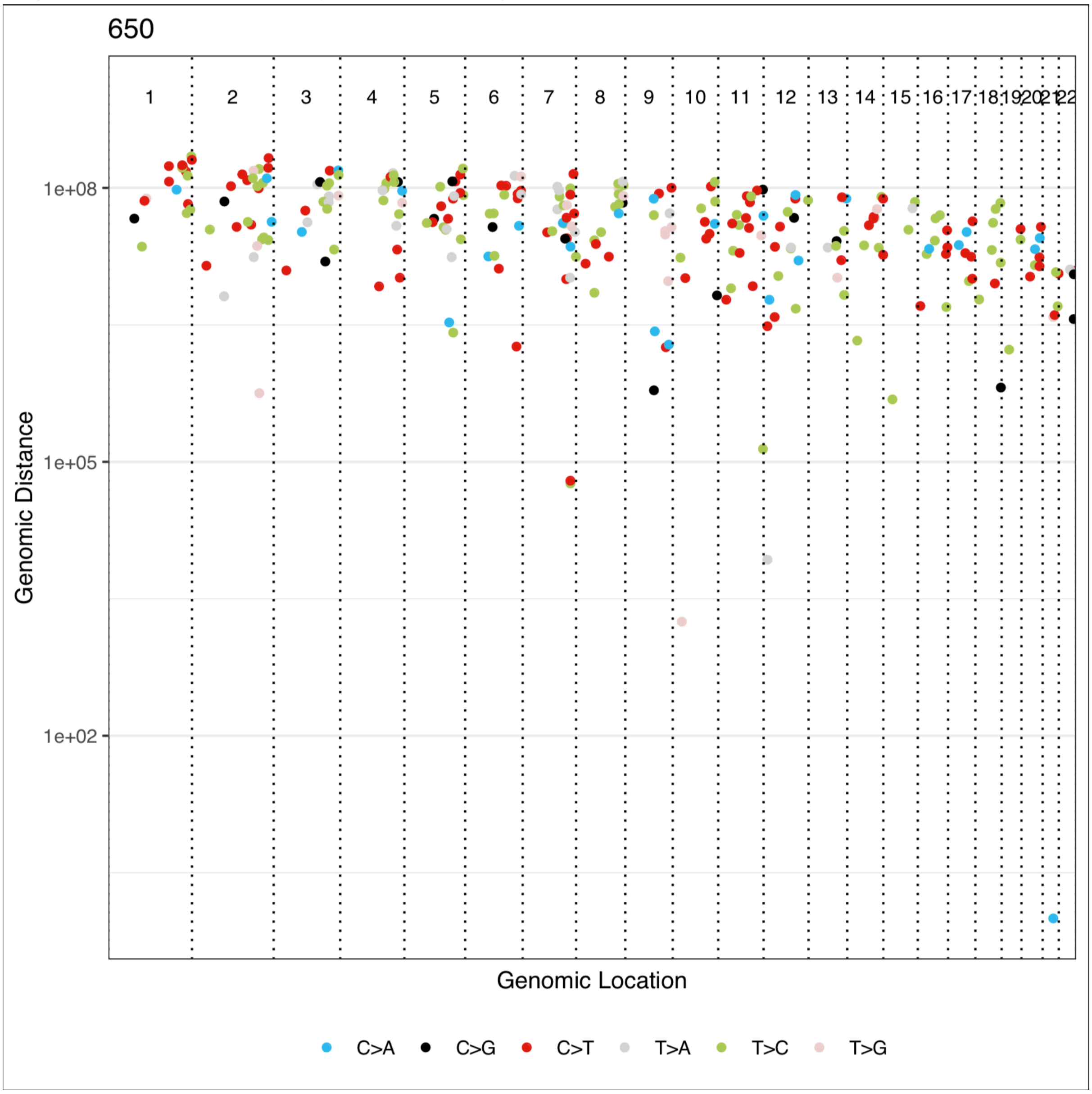


**Figure S01:** Rainfall plot of the 650 putative somatic SNVs as produced by the package MutationalPatterns [Blokzijl 2018]. A rainfall plot visualizes the types of mutations (colors), their genomic positioning (x-axis), and the distance between them (y-axis, log distance). Mutation hotspots would appear as clusters of mutations with lower intermutation distance.

#### Figure S02: Tri-nucleotide context of 650 putative SNVs

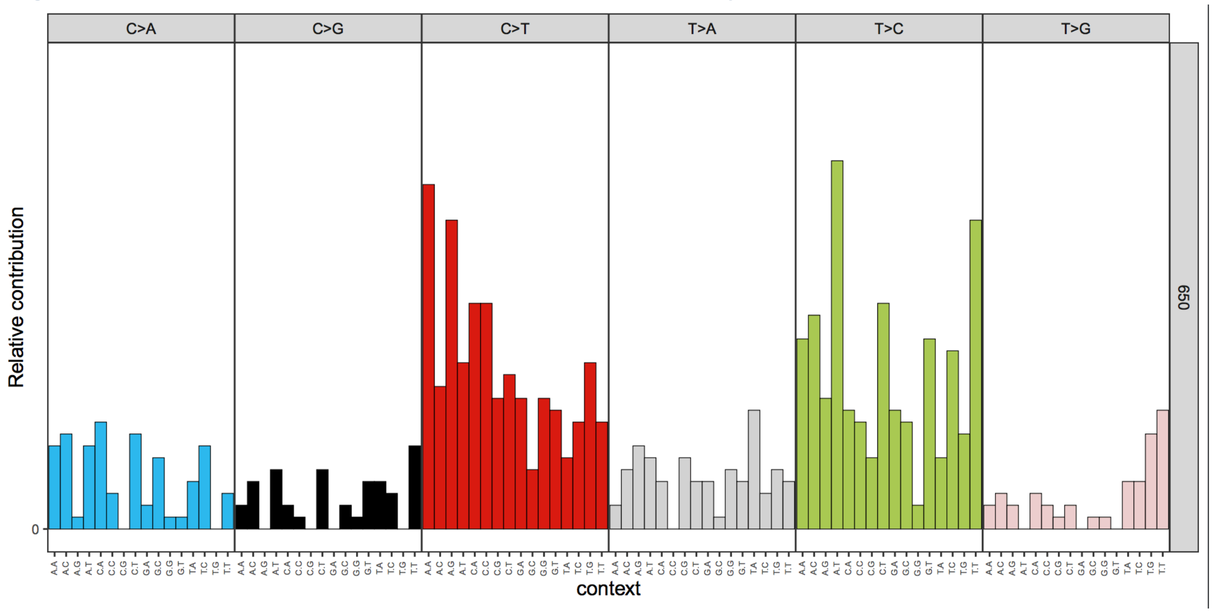


**Figure S02**: Relative contribution of each of the 96 possible trinucleotide combinations to the 650 putative SNVs as produced by the package MutationalPatterns [Blokzijl 2018]. Each category of single nucleotide substitutions (indicated at top) is further divided into 16 groups according to the neighbouring nucleotides (indicated at bottom). Resulting trinucleotide patterns are typically produced in cancer studies and have been associated with particular oncogenic processes [Nik-zainal 2012; Alexandrov 2013]. Interestingly, some of these trinucleotide patterns discovered in a pan-cancer setting have been associated with the age of diagnosis, and seem to represent ageing processes [Osorio 2018; Alexandrov 2015].

### Figure S03: Mutational signatures contributing to 650 putative SNVs

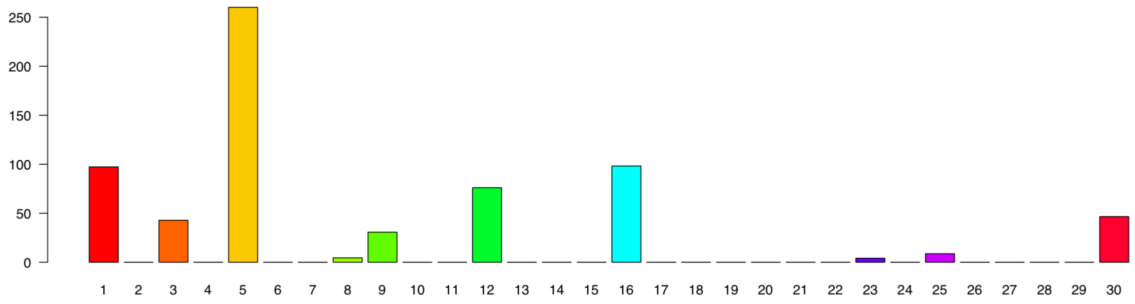


**Figure S03:** The number of mutations predicted to be derived from each of the 30 mutational signatures (COSMIC V2 – March 2015). Predictions are based on a linear combination of mutational signatures that most closely reconstructs the mutational pattern observed in this study (Figure S2), as computed by the package MutationalPatterns [Blokzijl 2018]. Next to signatures 1 and 5, considered to be ageing signatures, also signatures 12 and 16 seem to explain a lot of the observed mutational pattern. Both these signatures are typically observed in liver cancer, however, and typically exhibit a strong transcriptional strand bias. A closer inspection of our mutational profile did not reveal such a transcriptional strand bias (data not shown), hence associations with signatures 12 and 16 were discarded.

### Figure S04: Pairwise scatter plots of 307 SNVs at PB0, PB1 & PB2

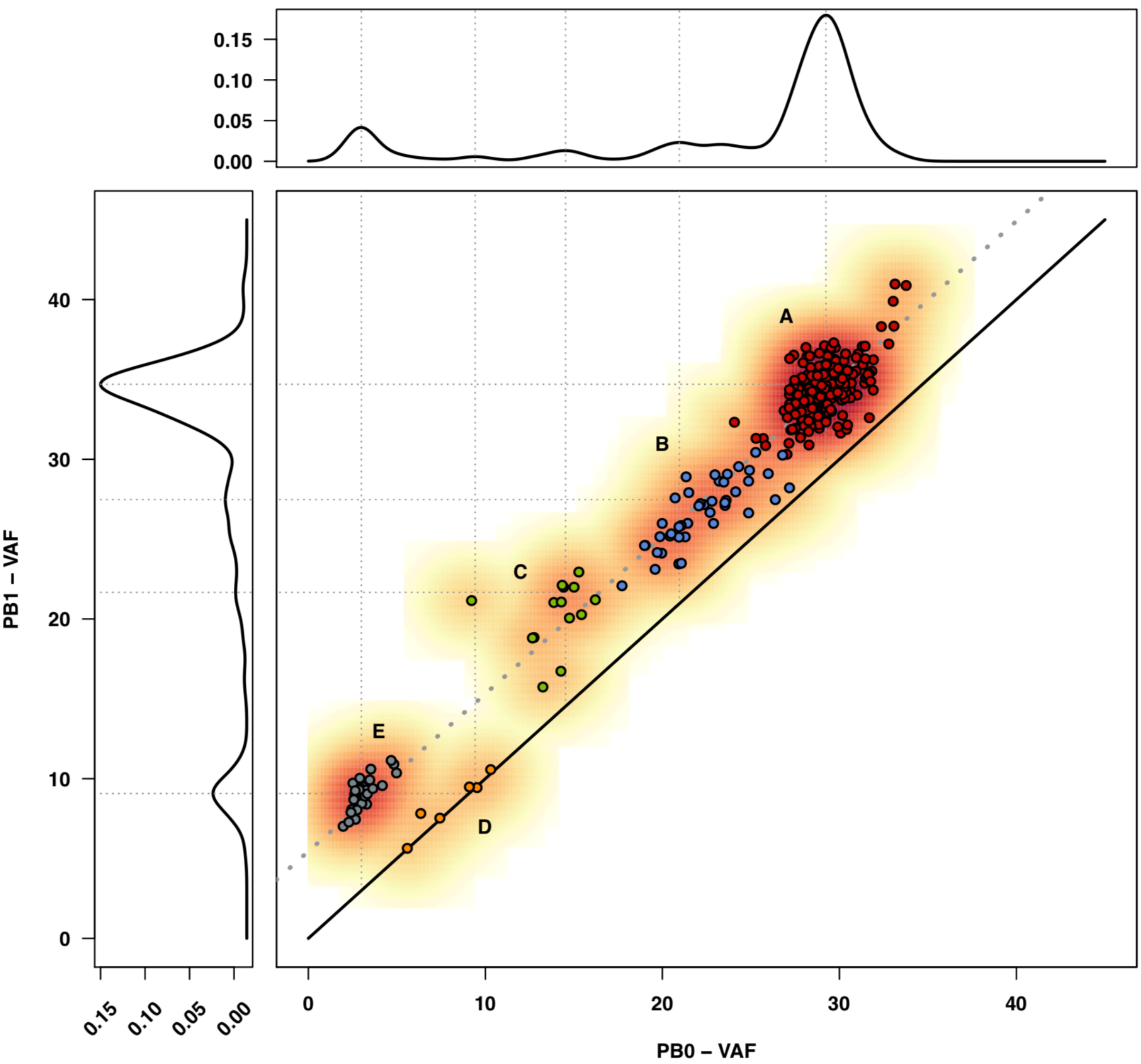


**Figure S04A:** Pairwise scatter plot of Variant Allele Frequencies (VAF) in peripheral blood measured at age 103 (PB0) and 110 (PB1). Each dot indicates a SNV, and is coloured according to subclonal event (A-E). Shades of red represent the local density. Density distributions at the sides represent the density distributions of VAFs at age 110 (y-axis; PB1) and age 103 (x-axis; PB0) respectively.

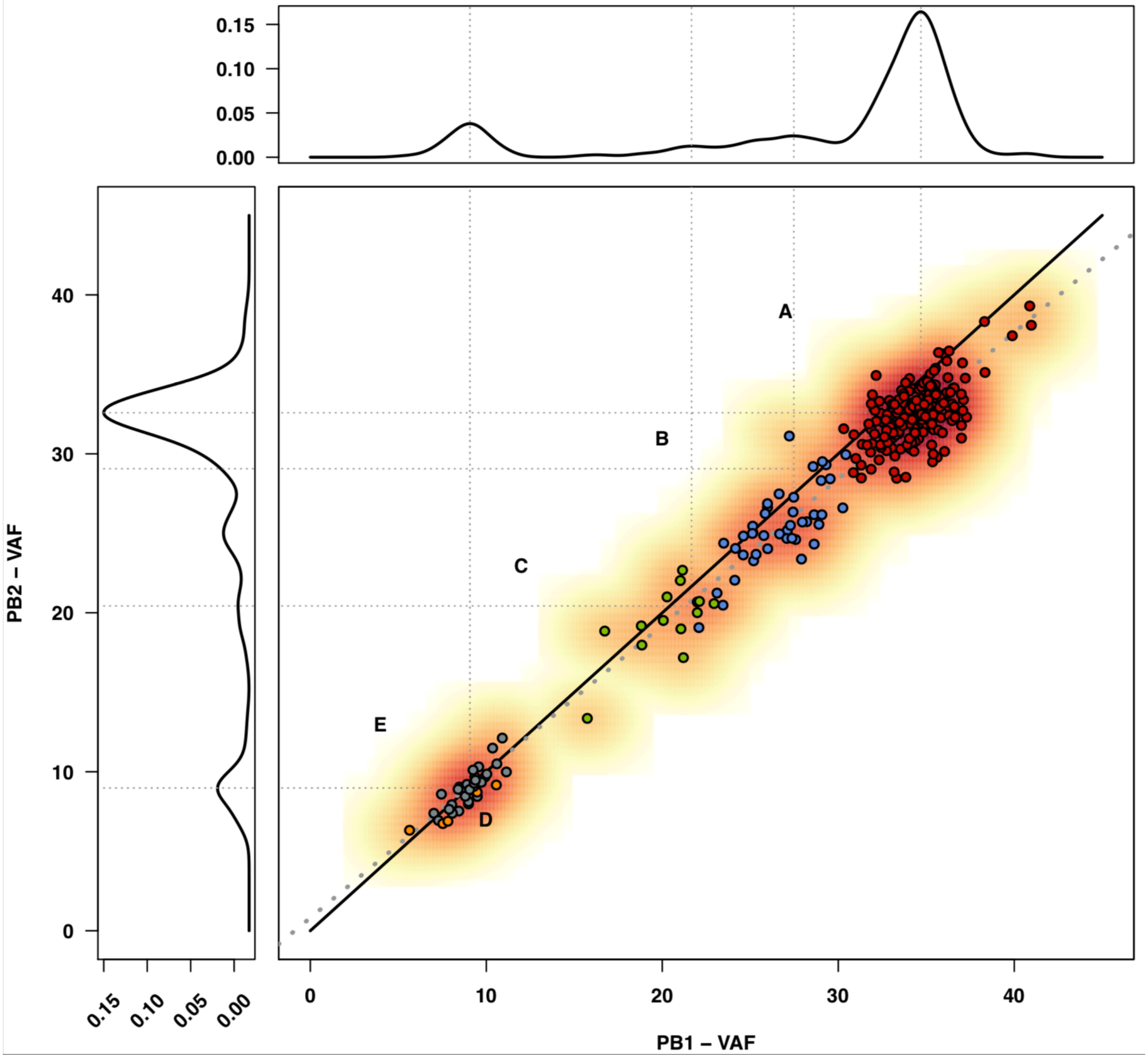


**Figure S04B:** Pairwise scatter plot of Variant Allele Frequencies (VAF) in peripheral blood measured at age 110 (PB1) and 111 (PB2). Each dot indicates a SNV, and is coloured according to subclonal event (A-E). Shades of red represent the local density. Density distributions at the sides represent the density distributions of VAFs at age 110 (y-axis; PB1) and age 111 (x-axis; PB2) respectively.

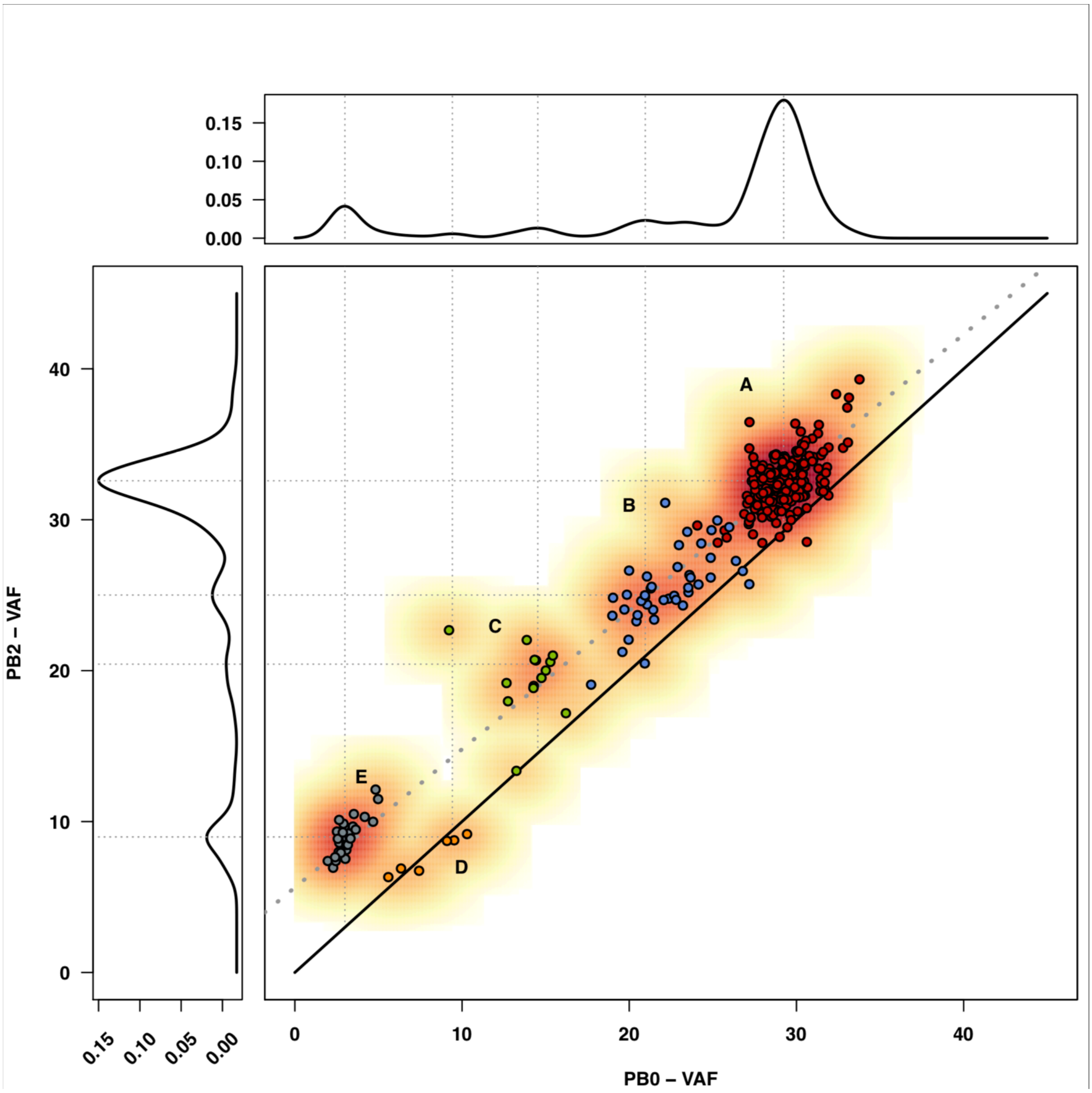


**Figure S04C:** Pairwise scatter plot of Variant Allele Frequencies (VAF) in peripheral blood measured at age 103 (PB0) and 111 (PB2). Each dot indicates a SNV, and is coloured according to subclonal event (A-E). Shades of red represent the local density. Density distributions at the sides represent the density distributions of VAFs at age 103 (y-axis; PB0) and age 111 (x-axis; PB2) respectively.

### Figure S05: Testing for suspect clonal BCR/TCR gene recombinations

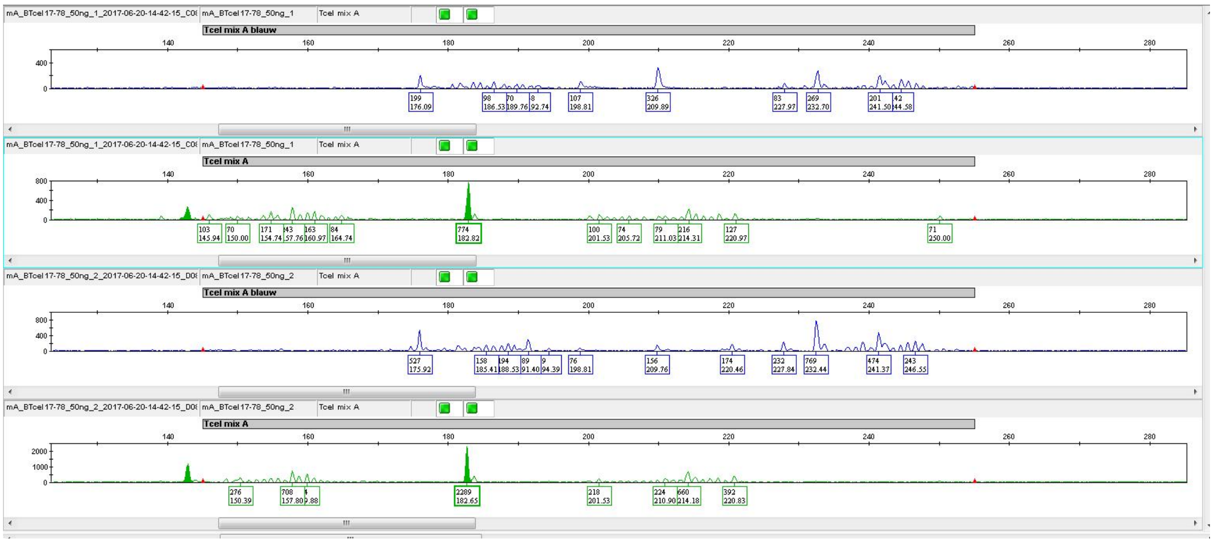

**Figure S06A**: **T-cell mix A**

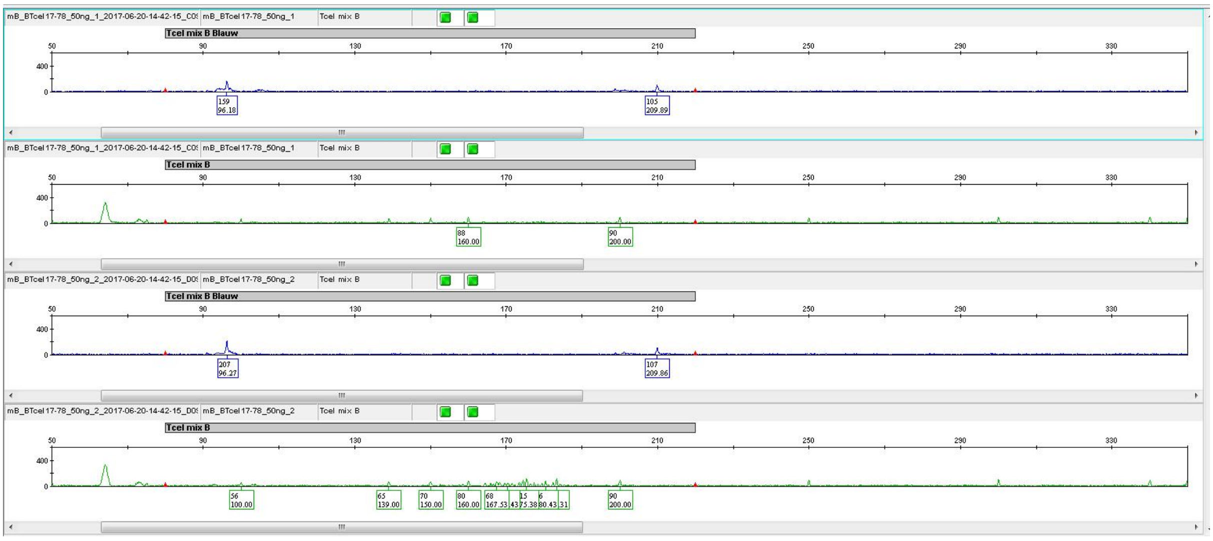

**Figure S06B**: **T-cell mix B**

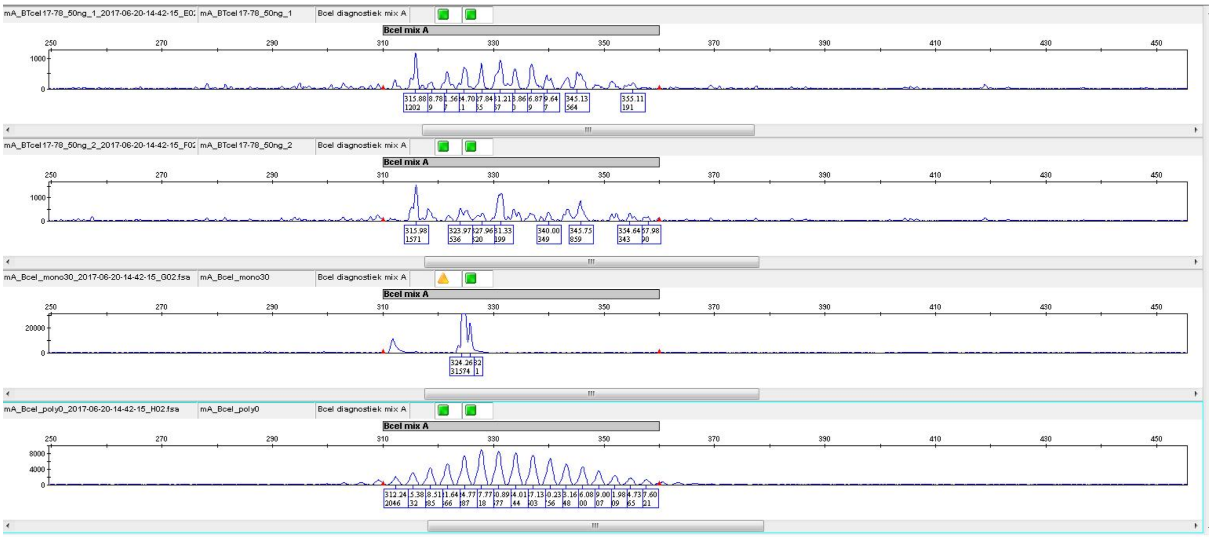


**Figure S06C**: **B-cell mix A**

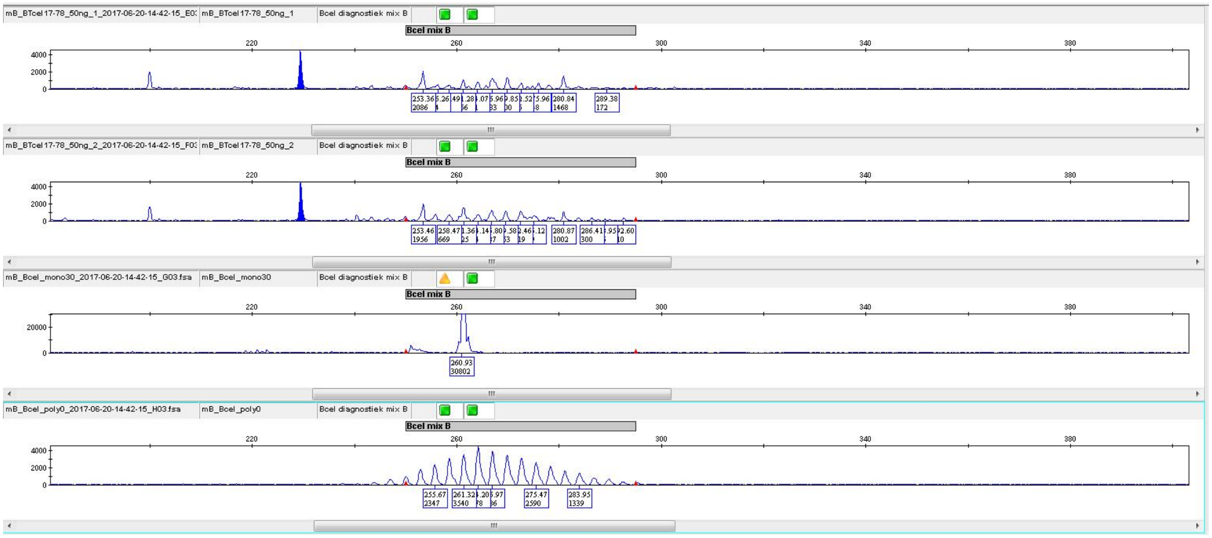


**Figure S06D**: **B-cell mix B**

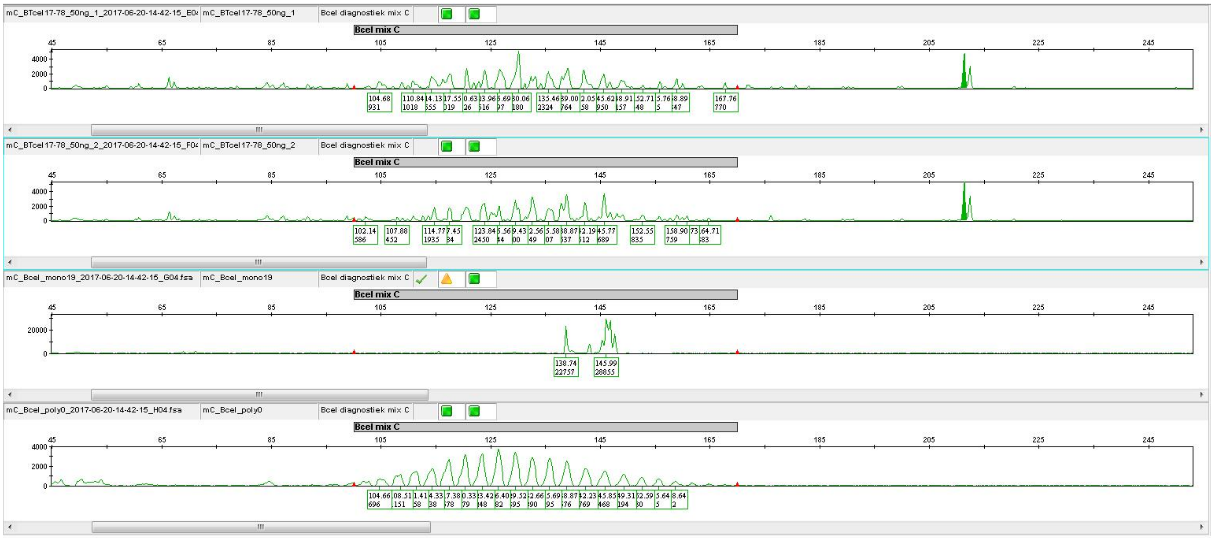

**Figure S06E**: **B-cell mix C**

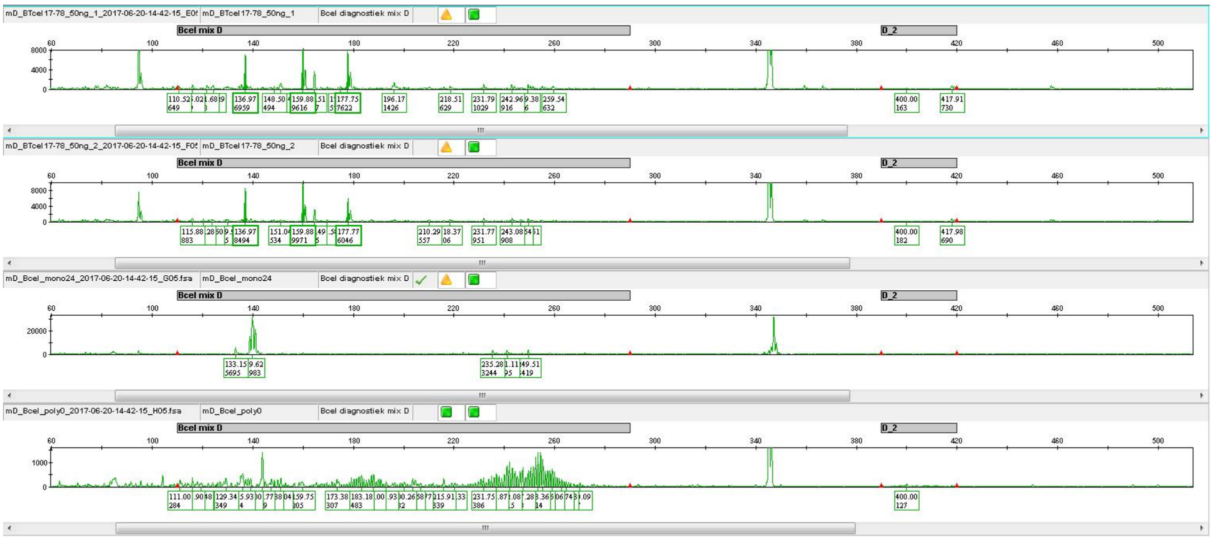

**Figure S06F**: **B-cell mix D**

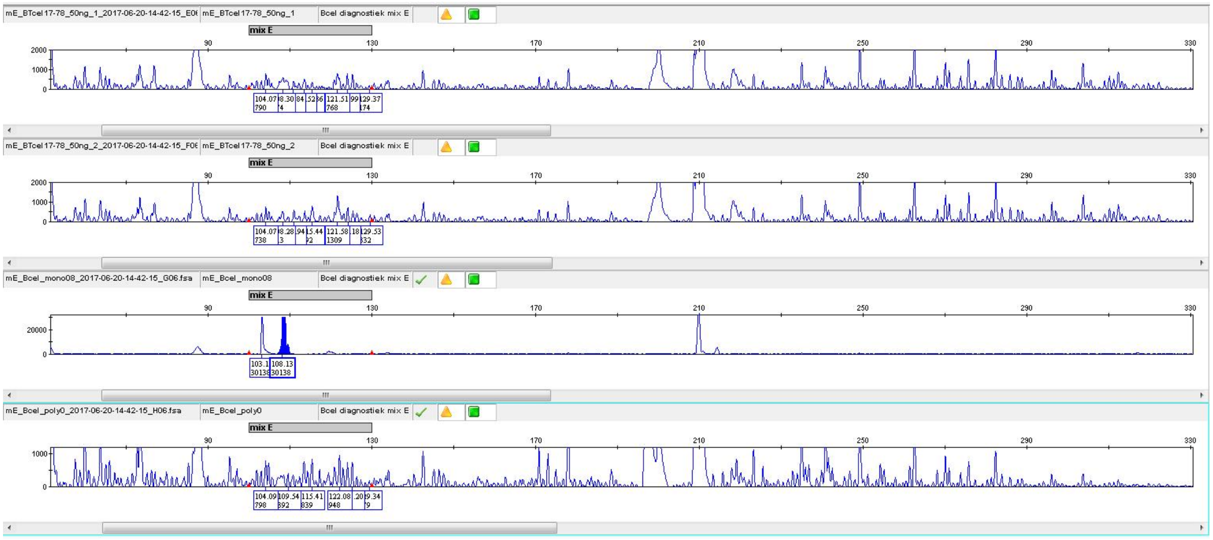


**Figure S06G**: **B-cell mix E**

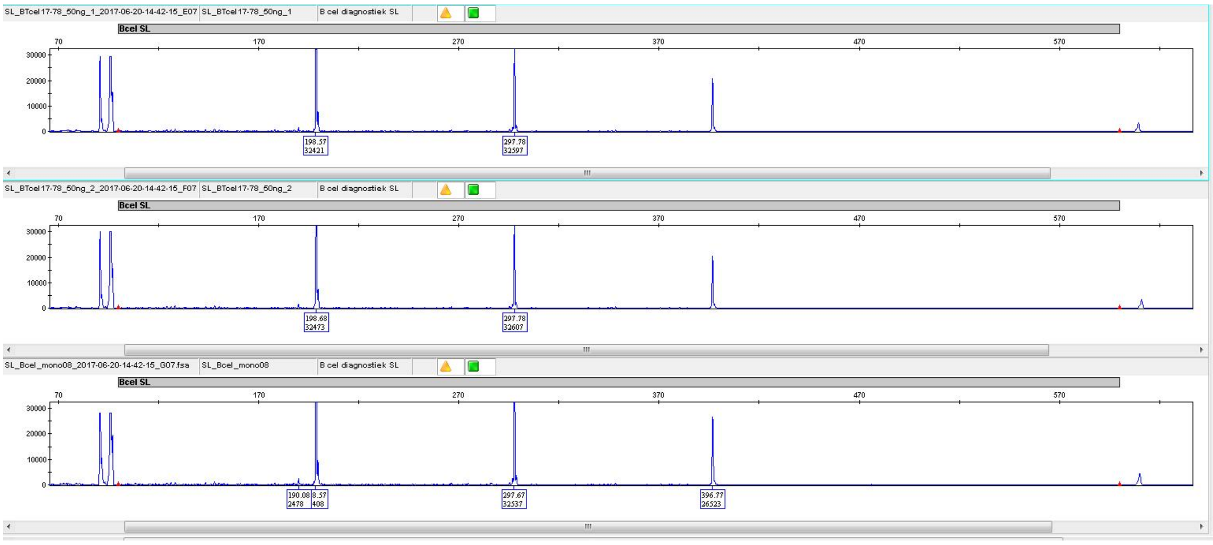


**Figure S06H**: **B-cell SL**

#### Figure S06: T-cell characterization at age 110 by flow cytometry

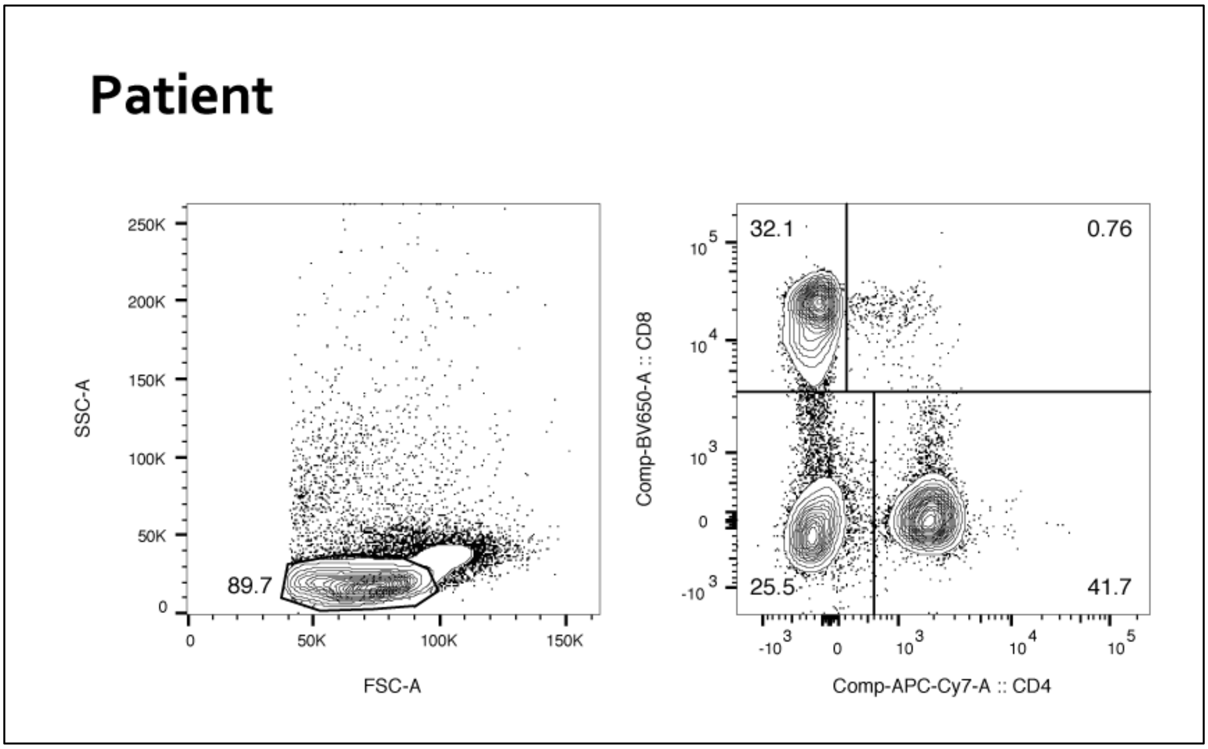


**Figure S07A**: **Lymphocyte gate in W110:** 44,330 cells enter (left), are gated to 39,771 cells to be further gated (right) to 16,591 (41.7%) CD4+ T-cells, and 12,574 (32.1%) CD8+ T-cells.

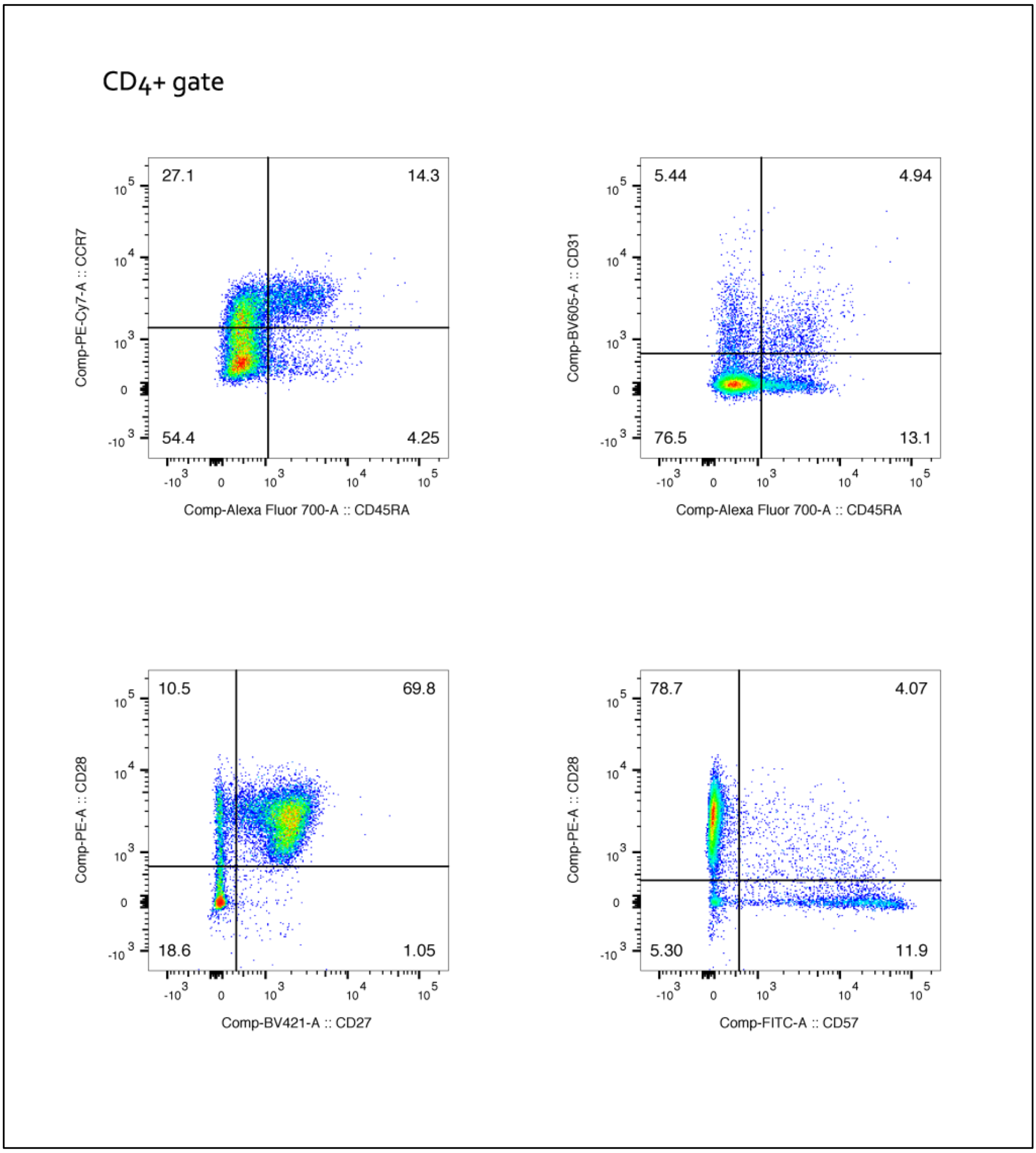


**Figure S07B: CD4+ Tcell gating in patient W110:** 16,591 CD4+ T-cells are analysed in each of these pairwise scatters; top left: CCR7-CD45RA+ (CD4+TEMRA); top right CD31+ CD45RA+ (recent thymic immigrants); bottom left: CD27+CD28+; bottom right: CD57+CD28- (senescent).

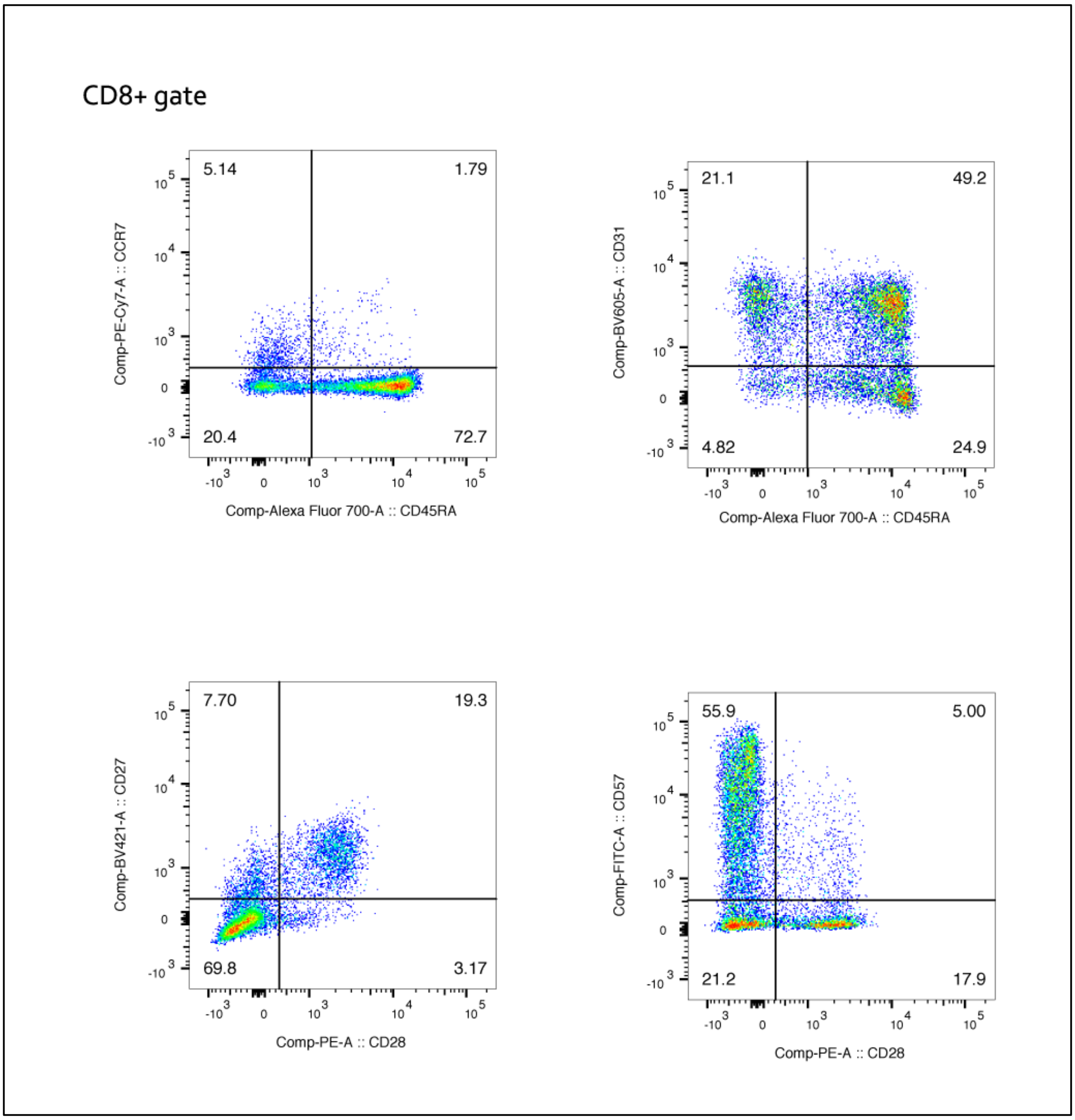


**Figure S07C: CD8+ Tcell gating in patient W110:** 12,574 CD8+ T-cells are analysed in each of these pairwise scatters; top left: CCR7-CD45RA+ (CD8+TEMRA); top right CD31+ CD45RA+ (done for consistency); bottom left: CD27+CD28+; bottom right: CD57+CD28- (senescent).

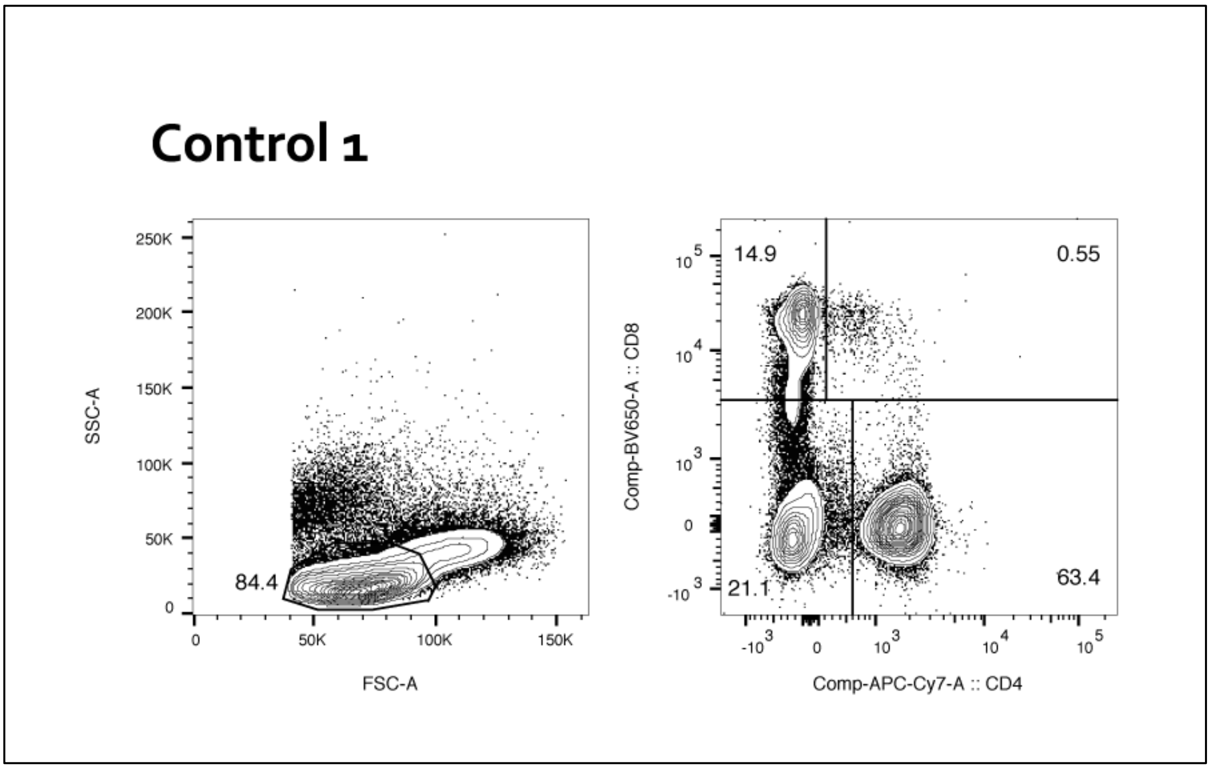


**Figure S07D**: **Lymphocyte gate in Control 1:** 117,956 cells enter (left), are gated to 99,514 cells to be further gated (right) to 63,105 (63.4%) CD4+ T-cells, and 14,862 (14,9%) CD8+ T-cells.

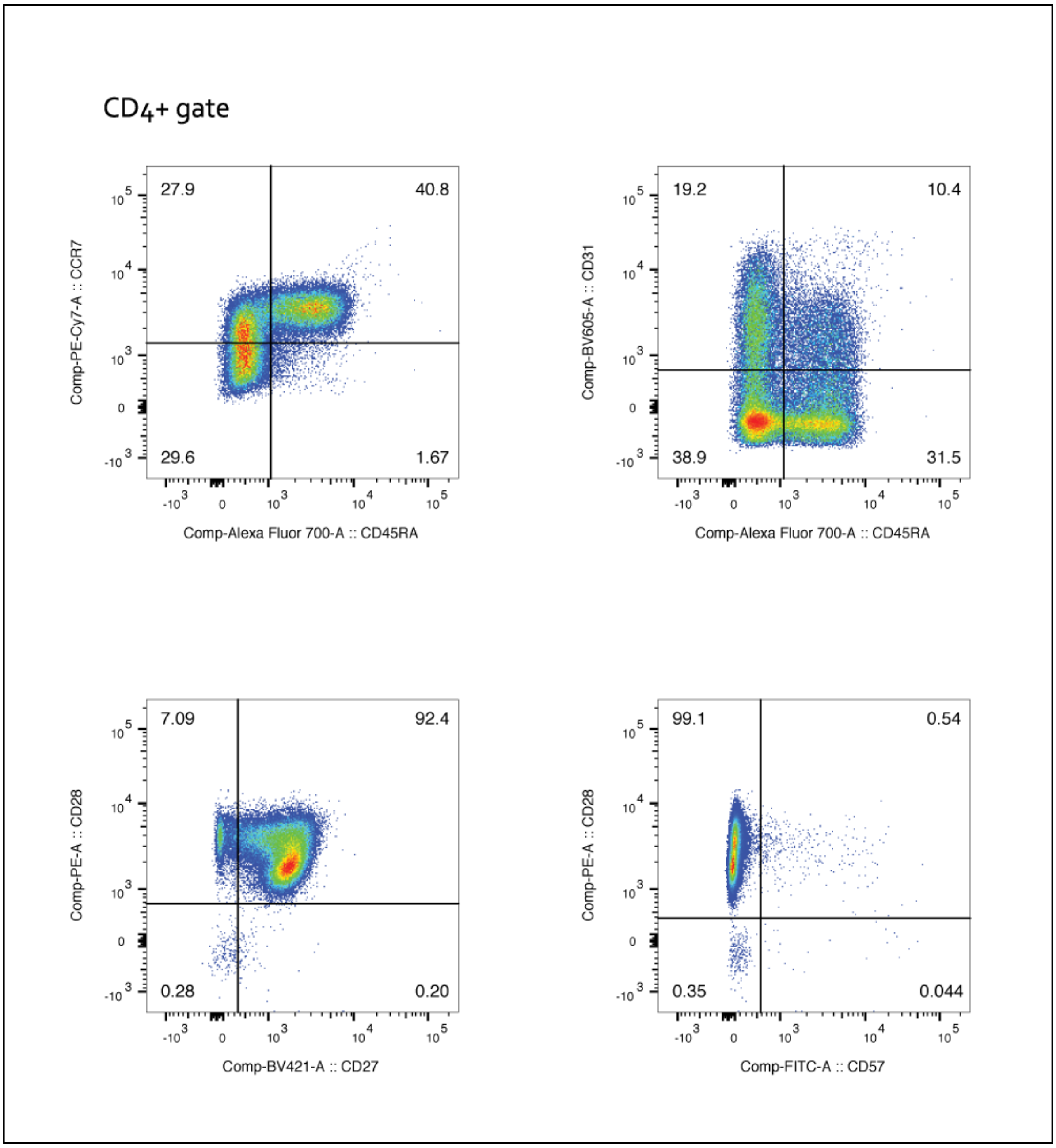


**Figure S07E: CD4+ Tcell gating in control 1:** 63,105 CD4+ T are analysed in each of these pairwise scatters; top left: CCR7-CD45RA+ (CD4+TEMRA); top right CD31+ CD45RA+ (recent thymic immigrants); bottom left: CD27+CD28+; bottom right: CD57+CD28- (senescent).

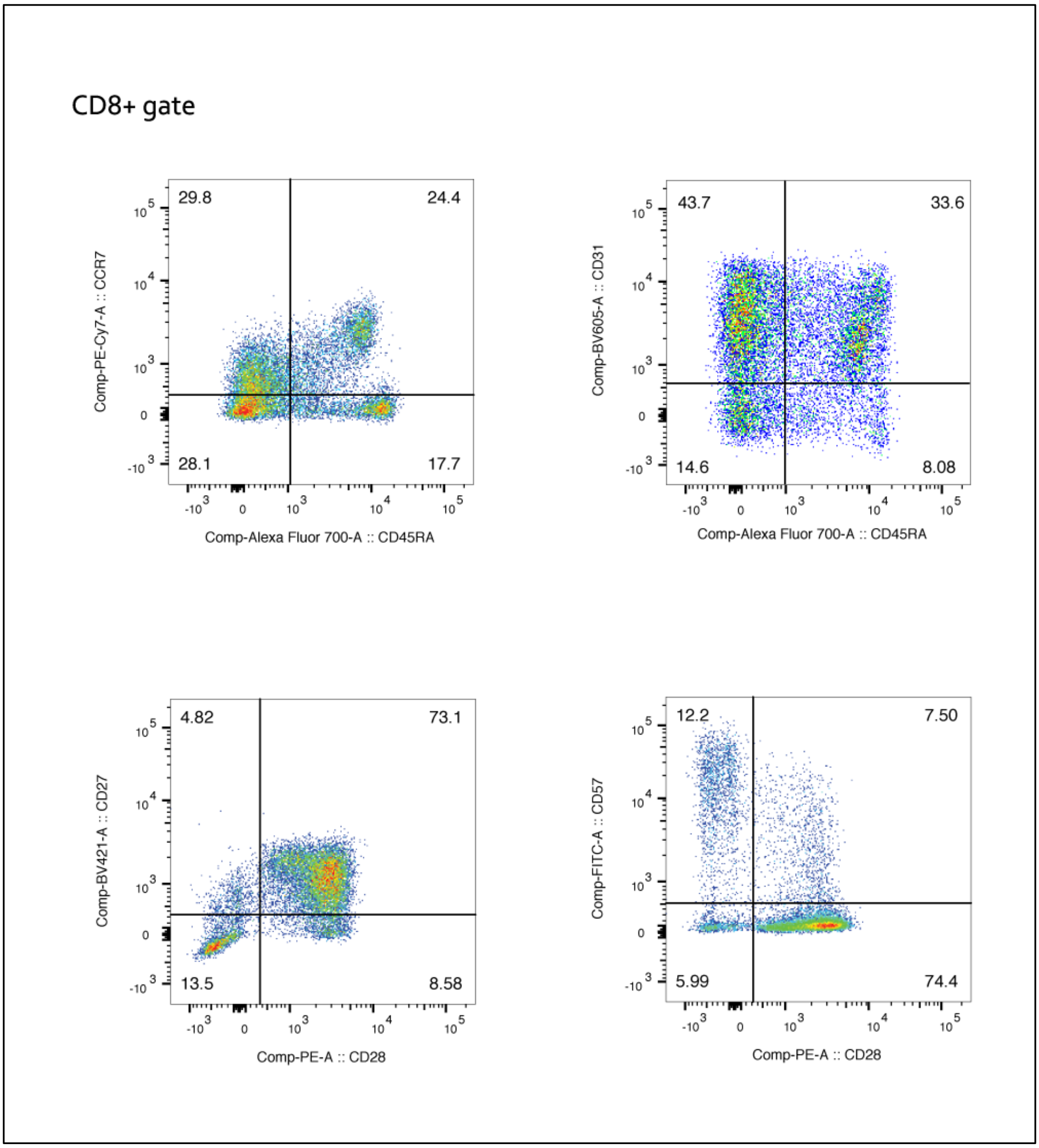


**Figure S07F: CD8+ Tcell gating in control 1:** 14,862 CD8+ T-cells are analysed in each of these pairwise scatters; top left: CCR7-CD45RA+ (CD8+TEMRA); top right CD31+ CD45RA+ (done for consistency); bottom left: CD27+CD28+; bottom right: CD57+CD28- (senescent).

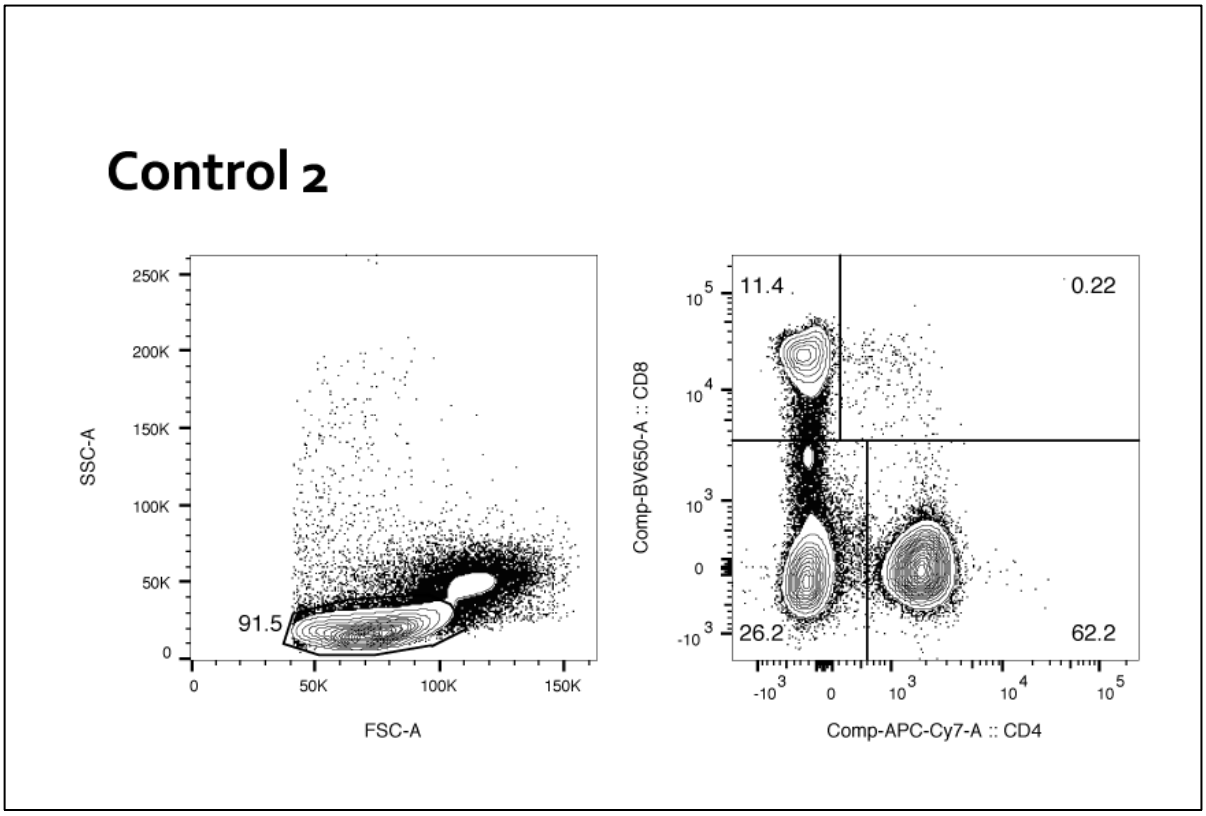


**Figure S07G**: **Lymphocyte gate in Control 2:** 114,564 cells enter (left), are gated to 104,799 cells to be further gated (right) to 65,196 (62.2%) CD4+ T-cells, and 11,968 (11,4%) CD8+ T-cells.

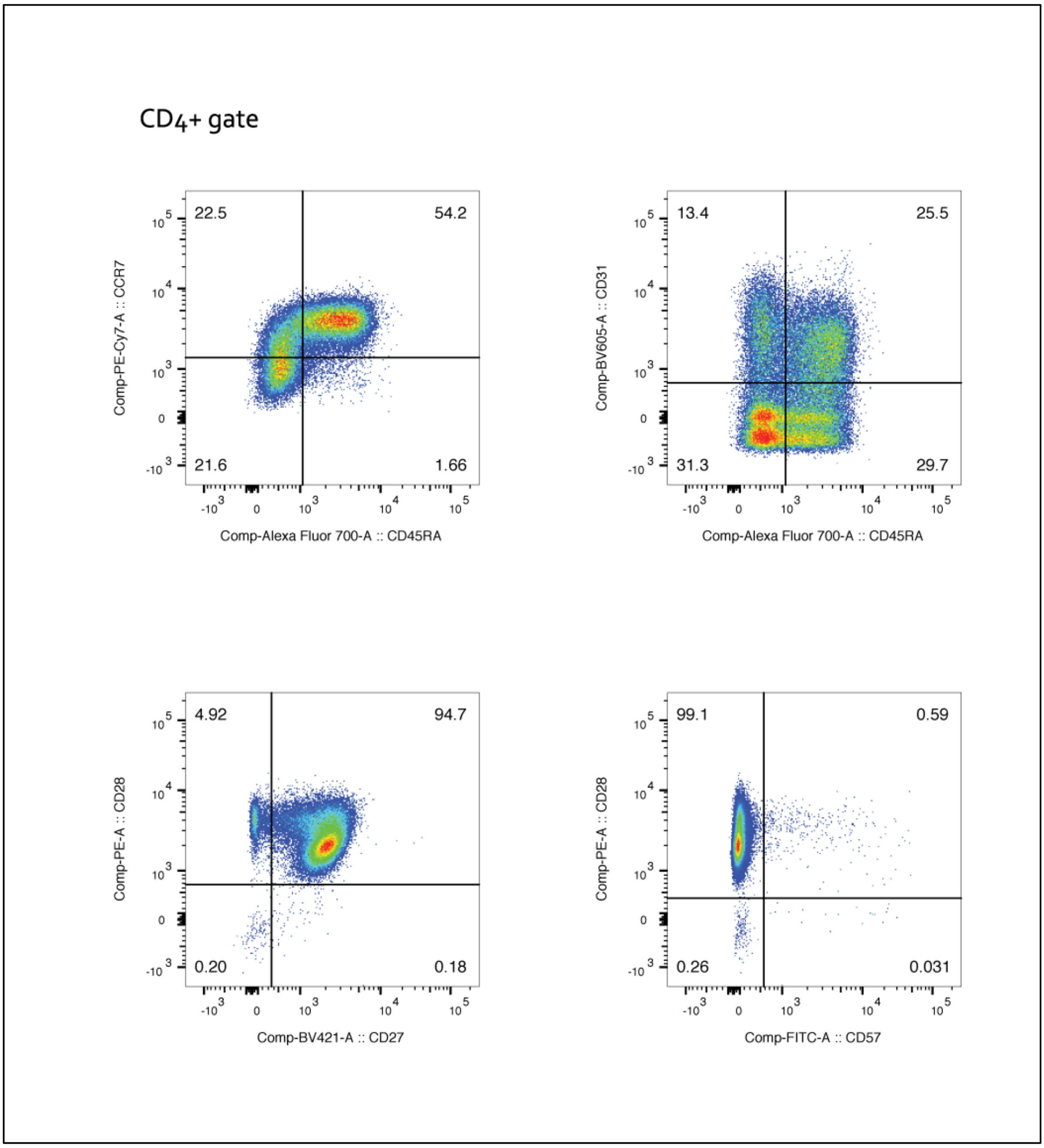


**Figure S07H: CD4+ Tcell gating in control 2:** 65,196 CD4+ T are analysed in each of these pairwise scatters; top left: CCR7-CD45RA+ (CD4+TEMRA); top right CD31+ CD45RA+ (recent thymic immigrants); bottom left: CD27+CD28+; bottom right: CD57+CD28- (senescent).

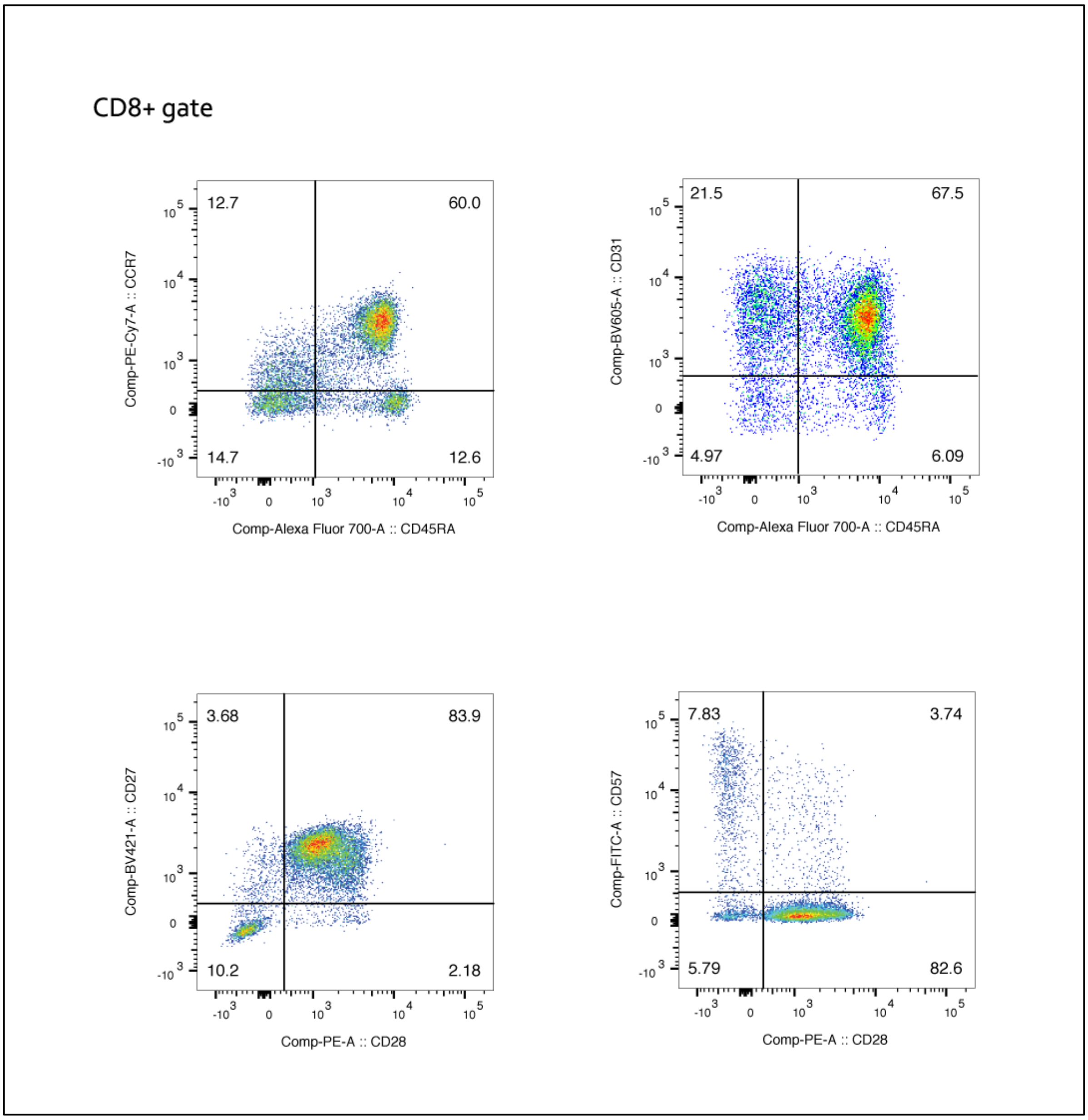


**Figure S07I: CD8+ Tcell gating in control 2:** 11,968 CD8+ T-cells are analysed in each of these pairwise scatters; top left: CCR7-CD45RA+ (CD8+TEMRA); top right CD31+ CD45RA+ (done for consistency); bottom left: CD27+CD28+; bottom right: CD57+CD28- (senescent).

#### Figure S07: T-cell characterization at age 111 by flow cytometry

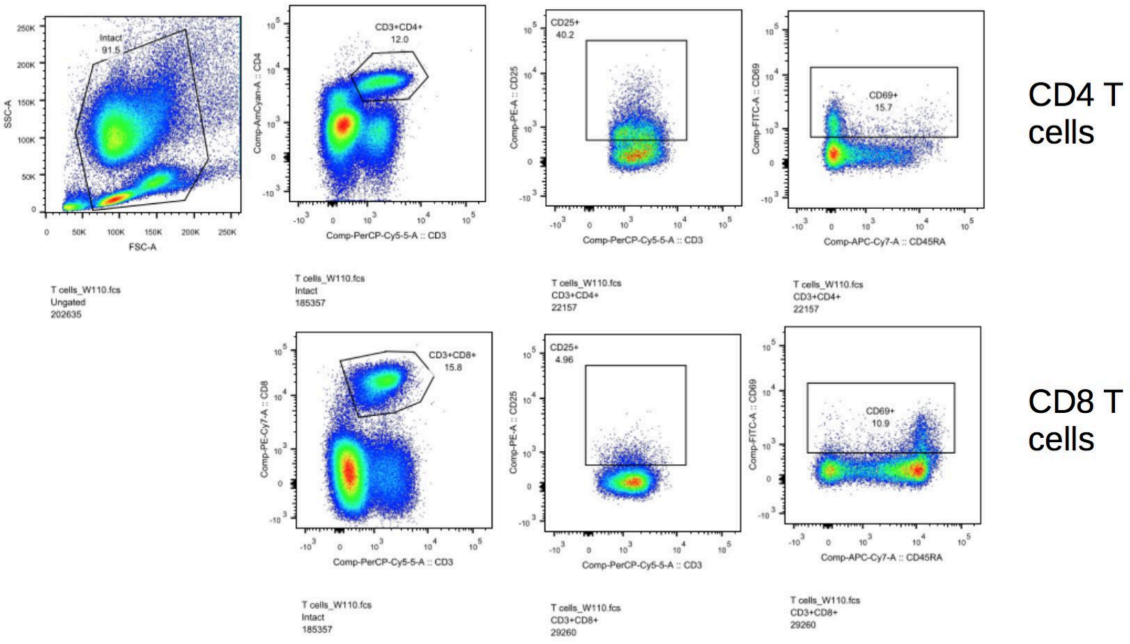

### Supplementary Tables

#### TABLE S1: 650 Putative Somatic Mutations

| gvarID | ref | alt | tier | VAF |
|---|---|---|---|---|
| chr17__66672231 | C | T | Blood_Tier1 | 0.467 |
| chr20__14934240 | C | T | Blood_Tier1 | 0.488 |
| chr11__123959954 | G | C | Blood_Tier1 | 0.468 |
| chr6__79616543 | A | G | Blood_Tier1 | 0.469 |
| chr6__156444677 | C | T | Blood_Tier1 | 0.438 |
| chr3__110499760 | A | G | Blood_Tier1 | 0.373 |
| chr2__164563143 | A | G | Blood_Tier1 | 0.417 |
| chr5__84789837 | G | A | Blood_Tier1 | 0.412 |
| chr3__192960239 | T | G | Blood_Tier1 | 0.338 |
| chr22__40724186 | G | A | Blood_Tier1 | 0.455 |
| chr2__15721915 | T | C | Blood_Tier1 | 0.369 |
| chr7__29217270 | A | G | Blood_Tier1 | 0.472 |
| chr1__79469387 | T | C | Blood_Tier1 | 0.425 |
| chrX__34276891 | T | G | Blood_Tier1 | 0.420 |
| chr6__150816248 | G | A | Blood_Tier1 | 0.330 |
| chr4__145441371 | A | T | Blood_Tier1 | 0.364 |
| chr10__71833394 | G | A | Blood_Tier1 | 0.333 |
| chr1__231042202 | C | T | Blood_Tier1 | 0.333 |
| chr2__228258480 | G | A | Blood_Tier1 | 0.458 |
| chr9__37342120 | C | T | Blood_Tier1 | 0.376 |
| chr4__24761759 | C | A | Blood_Tier1 | 0.367 |
| chr5__26329781 | G | C | Blood_Tier1 | 0.366 |
| chr13__105958180 | A | G | Blood_Tier1 | 0.500 |
| chr19__29093087 | C | G | Blood_Tier1 | 0.327 |
| chr6__152647273 | G | A | Blood_Tier1 | 0.354 |
| chr7__109719194 | A | G | Blood_Tier2 | 0.500 |
| chr20__25617183 | G | A | Blood_Tier1 | 0.432 |
| chr8__13478301 | T | C | Blood_Tier1 | 0.329 |
| chrX__129344146 | C | T | Blood_Tier1 | 0.389 |
| chr14__25504442 | T | C | Blood_Tier1 | 0.355 |
| chr15__101885527 | T | A | Blood_Tier1 | 0.444 |
| chr9__137633163 | G | A | Blood_Tier1 | 0.397 |
| chr14__64495445 | G | A | Blood_Tier1 | 0.395 |
| chr17__57689194 | T | C | Blood_Tier1 | 0.370 |
| chr2__141131460 | C | T | Blood_Tier1 | 0.393 |
| chr4__137409760 | T | C | Blood_Tier1 | 0.365 |
| chr4__55441256 | C | T | Blood_Tier1 | 0.383 |
| chr6__126113509 | T | C | Blood_Tier1 | 0.373 |
| chr15__77477740 | A | C | Blood_Tier1 | 0.431 |
| chr11__119595518 | C | T | Blood_Tier1 | 0.345 |
| chr9__19516480 | C | A | Blood_Tier1 | 0.299 |

| | | | | |
|---|---|---|---|---|
| chr6__44324229 | C | G | Blood_Tier1 | 0.266 |
| chr5__166531089 | C | T | Blood_Tier1 | 0.330 |
| chr7__82293864 | A | G | Blood_Tier1 | 0.333 |
| chr3__176430705 | G | A | Blood_Tier1 | 0.328 |
| chr1__170805330 | A | G | Blood_Tier1 | 0.348 |
| chr5__92280354 | C | A | Blood_Tier1 | 0.346 |
| chr20__39862424 | T | C | Blood_Tier1 | 0.315 |
| chr17__8280395 | G | A | Blood_Tier1 | 0.382 |
| chr4__171755715 | C | G | Blood_Tier1 | 0.325 |
| chr8__28291900 | G | A | Blood_Tier1 | 0.292 |
| chr6__81591019 | C | G | Blood_Tier1 | 0.348 |
| chr4__169089352 | T | A | Blood_Tier1 | 0.333 |
| chr14__26630906 | G | A | Blood_Tier1 | 0.295 |
| chr15__24051898 | C | A | Blood_Tier1 | 0.386 |
| chrX__84392105 | A | G | Blood_Tier1 | 0.342 |
| chr4__30984482 | G | T | Blood_Tier1 | 0.329 |
| chr6__51310525 | C | G | Blood_Tier1 | 0.271 |
| chr12__86416207 | G | A | Blood_Tier1 | 0.384 |
| chr7__122972688 | A | G | Blood_Tier1 | 0.333 |
| chr7__155804425 | A | T | Blood_Tier1 | 0.296 |
| chr11__18060193 | G | T | Blood_Tier1 | 0.435 |
| chr13__35027179 | A | T | Blood_Tier1 | 0.301 |
| chr8__126819256 | T | G | Blood_Tier1 | 0.340 |
| chr9__13732822 | T | A | Blood_Tier1 | 0.258 |
| chr6__69085881 | C | A | Blood_Tier1 | 0.340 |
| chr11__24024971 | G | A | Blood_Tier1 | 0.288 |
| chr8__17407813 | G | A | Blood_Tier1 | 0.286 |
| chr7__40599736 | G | A | Blood_Tier1 | 0.308 |
| chr4__11638973 | C | T | Blood_Tier1 | 0.323 |
| chr11__44526879 | T | C | Blood_Tier1 | 0.373 |
| chr16__53424032 | T | A | Blood_Tier1 | 0.371 |
| chr17__31969569 | C | A | Blood_Tier1 | 0.360 |
| chr3__71590489 | G | T | Blood_Tier1 | 0.255 |
| chr6__87120455 | T | C | Blood_Tier1 | 0.333 |
| chr10__99642604 | C | T | Blood_Tier1 | 0.356 |
| chr10__25106383 | A | C | Blood_Tier1 | 0.315 |
| chr1__237498066 | G | A | Blood_Tier1 | 0.257 |
| chr16__24450792 | G | A | Blood_Tier1 | 0.313 |
| chr8__14220884 | T | C | Blood_Tier1 | 0.342 |
| chr13__57257821 | A | T | Blood_Tier1 | 0.409 |
| chrX__66876515 | T | C | Blood_Tier1 | 0.290 |
| chr20__53722528 | G | A | Blood_Tier1 | 0.333 |
| chr2__38652340 | C | T | Blood_Tier2 | 0.458 |
| chr16__50955663 | T | C | Blood_Tier1 | 0.339 |
| chr1__216608363 | C | T | Blood_Tier1 | 0.296 |
| chr1__37164781 | T | A | Blood_Tier1 | 0.327 |

| | | | | |
|---|---|---|---|---|
| chr2__199917885 | A | G | Blood_Tier1 | 0.278 |
| chr1__113621193 | T | G | Blood_Tier1 | 0.355 |
| chr8__125435359 | A | G | Blood_Tier1 | 0.294 |
| chr20__36061512 | G | A | Blood_Tier1 | 0.308 |
| chrX__137615166 | A | T | Blood_Tier1 | 0.308 |
| chr17__51301725 | C | T | Blood_Tier1 | 0.373 |
| chr11__22783931 | T | C | Blood_Tier1 | 0.238 |
| chr10__84756078 | T | C | Blood_Tier2 | 0.429 |
| chr14__50105572 | A | G | Blood_Tier1 | 0.232 |
| chr5__175742143 | A | G | Blood_Tier1 | 0.364 |
| chr1__48839483 | T | G | Blood_Tier1 | 0.302 |
| chr5__42290327 | C | T | Blood_Tier1 | 0.257 |
| chr4__157045476 | T | A | Blood_Tier1 | 0.305 |
| chr4__17857259 | T | A | Blood_Tier1 | 0.277 |
| chr5__88243810 | C | G | Blood_Tier1 | 0.241 |
| chr18__9313391 | A | G | Blood_Tier2 | 0.321 |
| chr16__2645613 | C | T | Blood_Tier1 | 0.282 |
| chr2__200482779 | A | C | Blood_Tier1 | 0.341 |
| chr13__83427582 | G | C | Blood_Tier1 | 0.350 |
| chr10__125225251 | G | T | Blood_Tier1 | 0.235 |
| chr8__34502821 | G | C | Blood_Tier1 | 0.269 |
| chr2__227317709 | T | C | Blood_Tier2 | 0.375 |
| chr4__149877106 | G | A | Blood_Tier1 | 0.191 |
| chr18__1392644 | A | C | Blood_Tier1 | 0.300 |
| chr2__203150320 | T | G | Blood_Tier1 | 0.315 |
| chr2__99565440 | G | T | Blood_Tier2 | 0.238 |
| chr11__62037391 | T | C | Blood_Tier1 | 0.279 |
| chr3__50930603 | G | A | Blood_Tier1 | 0.264 |
| chr5__123025785 | A | G | Blood_Tier1 | 0.316 |
| chr5__140487053 | A | T | Blood_Tier1 | 0.250 |
| chr8__141389872 | T | C | Blood_Tier1 | 0.292 |
| chrX__18897311 | C | G | Blood_Tier1 | 0.224 |
| chr22__44385037 | C | G | Blood_Tier1 | 0.232 |
| chr14__21773358 | G | A | Blood_Tier1 | 0.311 |
| chr3__83880717 | C | A | Blood_Tier1 | 0.222 |
| chr2__198609762 | G | A | Blood_Tier1 | 0.274 |
| chr6__100169580 | C | T | Blood_Tier1 | 0.308 |
| chrX__132397792 | A | C | Blood_Tier1 | 0.237 |
| chr3__164622904 | A | T | Blood_Tier1 | 0.241 |
| chr17__68940187 | C | T | Blood_Tier1 | 0.333 |
| chr4__148542277 | C | A | Blood_Tier1 | 0.219 |
| chr18__60094786 | T | C | Blood_Tier2 | 0.182 |
| chr1__222054608 | G | A | Blood_Tier1 | 0.213 |
| chr15__94305134 | T | C | Blood_Tier1 | 0.265 |
| chr2__190340213 | T | C | Blood_Tier2 | 0.200 |
| chr4__169675257 | G | A | Blood_Tier1 | 0.147 |

| | | | | |
|---|---|---|---|---|
| chr12__79160128 | C | A | Blood_Tier1 | 0.313 |
| chr11__55523417 | C | A | Blood_Tier1 | 0.221 |
| chr16__7739039 | G | A | Blood_Tier1 | 0.200 |
| chr20__53456479 | G | A | Blood_Tier1 | 0.213 |
| chr7__73217590 | G | A | Blood_Tier1 | 0.128 |
| chr3__141002881 | A | G | Blood_Tier2 | 0.250 |
| chr3__77327697 | A | C | Blood_Tier2 | 0.474 |
| chr2__61401283 | A | G | Blood_Tier1 | 0.314 |
| chr2__227183785 | C | T | Blood_Tier1 | 0.391 |
| chr6__16390236 | T | C | Blood_Tier1 | 0.321 |
| chr2__136630897 | T | A | Blood_Tier1 | 0.250 |
| chrX__50069526 | T | C | Blood_Tier2 | 0.235 |
| chr9__100480485 | G | A | Blood_Tier1 | 0.277 |
| chr19__56255629 | A | G | Blood_Tier1 | 0.322 |
| chr18__75296947 | T | C | Blood_Tier1 | 0.167 |
| chr1__86284096 | C | T | Blood_Tier1 | 0.204 |
| chr12__59303326 | T | C | Blood_Tier1 | 0.258 |
| chr11__3766585 | A | G | Blood_Tier1 | 0.286 |
| chr12__81010387 | A | G | Blood_Tier2 | 0.200 |
| chr4__102280086 | T | C | Blood_Tier1 | 0.229 |
| chrX__114116246 | A | C | Blood_Tier1 | 0.305 |
| chr1__53590585 | C | T | Blood_Tier1 | 0.225 |
| chr2__176211302 | C | T | Blood_Tier1 | 0.317 |
| chr7__45276483 | G | T | Blood_Tier1 | 0.197 |
| chrX__33045447 | A | G | Blood_Tier1 | 0.289 |
| chr4__24092455 | C | T | Blood_Tier1 | 0.184 |
| chr15__36040233 | G | A | Blood_Tier1 | 0.292 |
| chr10__131873809 | G | C | Blood_Tier1 | 0.222 |
| chr2__123856347 | C | G | Blood_Tier1 | 0.489 |
| chr5__167871848 | A | G | Blood_Tier2 | 0.200 |
| chr3__147908859 | T | C | Blood_Tier1 | 0.333 |
| chr14__101629779 | A | G | Blood_Tier2 | 0.160 |
| chr12__80759750 | A | G | Blood_Tier1 | 0.246 |
| chr4__21814933 | T | C | Blood_Tier2 | 0.182 |
| chr9__137585155 | T | G | Blood_Tier1 | 0.238 |
| chr8__47359871 | C | T | Blood_Tier2 | 0.333 |
| chr2__28226905 | G | C | Blood_Tier1 | 0.294 |
| chr6__121901998 | G | A | Blood_Tier1 | 0.184 |
| chr7__103470996 | T | A | Blood_Tier2 | 0.444 |
| chr10__86358071 | C | T | Blood_Tier1 | 0.239 |
| chr15__26802929 | A | G | Blood_Tier2 | 0.304 |
| chr22__34167420 | C | T | Blood_Tier1 | 0.234 |
| chr9__8517886 | G | A | Blood_Tier1 | 0.198 |
| chr21__27807421 | T | C | Blood_Tier1 | 0.293 |
| chr16__26597017 | A | G | Blood_Tier1 | 0.162 |
| chr10__53125352 | A | T | Blood_Tier1 | 0.274 |

| | | | | |
|---|---|---|---|---|
| chr5__34012260 | A | G | Blood_Tier1 | 0.263 |
| chr5__151631087 | G | A | Blood_Tier1 | 0.167 |
| chr13__36278145 | C | T | Blood_Tier2 | 0.121 |
| chr11__85068516 | G | A | Blood_Tier1 | 0.357 |
| chr3__31655114 | T | C | Blood_Tier2 | 0.174 |
| chr22__46538743 | G | A | Blood_Tier2 | 0.372 |
| chr17__67216242 | T | C | Blood_Tier2 | 0.111 |
| chrX__34047538 | A | T | Blood_Tier1 | 0.280 |
| chr20__46878505 | C | A | Blood_Tier1 | 0.325 |
| chr9__84717849 | G | T | Blood_Tier1 | 0.272 |
| chr5__79447974 | C | T | Blood_Tier1 | 0.227 |
| chr5__116573374 | T | C | Blood_Tier2 | 0.133 |
| chr5__84711280 | A | T | Blood_Tier1 | 0.155 |
| chr14__35339308 | G | C | Blood_Tier1 | 0.188 |
| chr4__159243915 | A | G | Blood_Tier1 | 0.337 |
| chr7__131572995 | C | T | Blood_Tier1 | 0.216 |
| chr16__20829538 | T | C | Blood_Tier2 | 0.148 |
| chr21__31669351 | T | G | Blood_Tier2 | 0.242 |
| chrX__85300646 | T | C | Blood_Tier1 | 0.114 |
| chr10__95590751 | G | A | Blood_Tier1 | 0.138 |
| chr3__12400772 | C | T | Blood_Tier2 | 0.143 |
| chr6__160372588 | C | A | Blood_Tier1 | 0.316 |
| chr20__20453847 | G | A | Blood_Tier1 | 0.258 |
| chr16__18023205 | G | A | Blood_Tier1 | 0.253 |
| chrX__147904819 | C | G | Blood_Tier1 | 0.172 |
| chr15__86765022 | A | T | Blood_Tier1 | 0.103 |
| chr6__20794608 | A | G | Blood_Tier1 | 0.279 |
| chr1__220183809 | A | G | Blood_Tier2 | 0.108 |
| chr8__54468375 | T | C | Blood_Tier1 | 0.232 |
| chr1__186581699 | T | A | Blood_Tier1 | 0.216 |
| chr12__29141533 | C | T | Blood_Tier1 | 0.217 |
| chr8__41745949 | G | A | Blood_Tier1 | 0.286 |
| chr6__73053278 | T | C | Blood_Tier2 | 0.357 |
| chr11__134641812 | G | T | Blood_Tier1 | 0.218 |
| chr10__78249857 | T | C | Blood_Tier2 | 0.308 |
| chr11__4399767 | A | T | Blood_Tier1 | 0.184 |
| chr3__166901364 | G | A | Blood_Tier1 | 0.303 |
| chr11__55240858 | T | C | Blood_Tier2 | 0.121 |
| chr11__91653831 | C | T | Blood_Tier1 | 0.333 |
| chr19__52010257 | G | A | Blood_Tier1 | 0.281 |
| chr20__57909208 | C | T | Blood_Tier1 | 0.174 |
| chr5__41912087 | T | C | Blood_Tier3 | 0.500 |
| chr1__244390672 | T | C | Blood_Tier1 | 0.253 |
| chr4__107474761 | A | T | Blood_Tier2 | 0.118 |
| chr5__84069060 | C | T | Blood_Tier3 | 0.444 |
| chr7__42936683 | A | G | Blood_Tier2 | 0.200 |

| | | | | |
|---|---|---|---|---|
| chr18__75948056 | C | G | Blood_Tier2 | 0.160 |
| chr12__32994254 | G | A | Blood_Tier2 | 0.480 |
| chr7__293540 | C | T | Blood_Tier2 | 0.450 |
| chr7__103470997 | T | A | Blood_Tier3 | 0.500 |
| chr7__98695779 | A | G | Blood_Tier2 | 0.235 |
| chr8__27484960 | C | T | Blood_Tier1 | 0.160 |
| chr11__62926293 | G | A | Blood_Tier2 | 0.154 |
| chr15__73296442 | C | T | Blood_Tier1 | 0.147 |
| chr11__37183794 | A | G | Blood_Tier1 | 0.127 |
| chr6__166226981 | G | A | Blood_Tier1 | 0.116 |
| chr16__6613470 | G | A | Blood_Tier1 | 0.230 |
| chr10__109656568 | C | T | Blood_Tier1 | 0.242 |
| chr5__130022910 | G | A | Blood_Tier1 | 0.259 |
| chr7__126483539 | G | C | Blood_Tier2 | 0.235 |
| chr7__78431782 | C | T | Blood_Tier1 | 0.222 |
| chr21__31669352 | G | T | Blood_Tier2 | 0.229 |
| chr21__35721670 | G | A | Blood_Tier1 | 0.228 |
| chr1__201313101 | A | T | Blood_Tier1 | 0.085 |
| chr1__7358343 | T | C | Blood_Tier1 | 0.242 |
| chr11__133074959 | G | C | Blood_Tier2 | 0.176 |
| chr5__133385008 | C | A | Blood_Tier2 | 0.255 |
| chr19__20809490 | C | A | Blood_Tier2 | 0.484 |
| chr7__119749743 | G | T | Blood_Tier2 | 0.225 |
| chr3__20628725 | T | C | Blood_Tier3 | 0.500 |
| chr9__85325108 | C | G | Blood_Tier2 | 0.167 |
| chr4__115824013 | G | A | Blood_Tier2 | 0.136 |
| chr13__112600346 | G | T | Blood_Tier1 | 0.280 |
| chr3__130030623 | A | T | Blood_Tier2 | 0.286 |
| chr12__43843430 | A | G | Blood_Tier2 | 0.147 |
| chr16__52979969 | A | G | Blood_Tier2 | 0.190 |
| chr22__29259269 | C | T | Blood_Tier2 | 0.320 |
| chrX__14959214 | G | A | Blood_Tier1 | 0.078 |
| chr3__52315833 | T | C | Blood_Tier1 | 0.350 |
| chr2__8958069 | T | C | Blood_Tier2 | 0.100 |
| chr18__70491670 | A | G | Blood_Tier2 | 0.271 |
| chr2__149796259 | C | T | Blood_Tier1 | 0.111 |
| chr3__165184061 | T | C | Blood_Tier2 | 0.235 |
| chr10__20491854 | G | A | Blood_Tier1 | 0.119 |
| chr8__54188756 | A | G | Blood_Tier2 | 0.375 |
| chr5__120286007 | C | A | Blood_Tier2 | 0.328 |
| chr1__180644884 | G | A | Blood_Tier1 | 0.108 |
| chr19__56322749 | G | A | Blood_Tier1 | 0.164 |
| chr5__68689451 | A | G | Blood_Tier2 | 0.133 |
| chr14__82551829 | T | A | Blood_Tier1 | 0.254 |
| chr16__87305621 | G | A | Blood_Tier1 | 0.230 |
| chr8__116308909 | T | C | Blood_Tier2 | 0.114 |

| | | | | |
|---|---|---|---|---|
| chr12__90867035 | C | G | Blood_Tier2 | 0.200 |
| chr9__88013515 | G | T | Blood_Tier2 | 0.255 |
| chr19__48149785 | G | A | Blood_Tier1 | 0.075 |
| chr17__29741611 | G | A | Blood_Tier2 | 0.245 |
| chr3__21941024 | C | A | Blood_Tier1 | 0.244 |
| chr10__10034173 | C | T | Blood_Tier2 | 0.139 |
| chr6__146584713 | T | C | Blood_Tier2 | 0.154 |
| chr3__138121382 | G | C | Blood_Tier2 | 0.232 |
| chr11__133213386 | A | G | Blood_Tier1 | 0.169 |
| chr14__33343977 | T | A | Blood_Tier2 | 0.410 |
| chrX__109490016 | T | C | Blood_Tier1 | 0.228 |
| chr5__144949915 | C | T | Blood_Tier1 | 0.213 |
| chr2__63601769 | T | C | Blood_Tier2 | 0.133 |
| chr4__184500714 | T | G | Blood_Tier2 | 0.375 |
| chr8__69906311 | G | A | Blood_Tier1 | 0.108 |
| chr11__103189459 | A | G | Blood_Tier3 | 0.273 |
| chr20__38230941 | G | A | Blood_Tier1 | 0.337 |
| chr7__129772266 | G | A | Blood_Tier1 | 0.139 |
| chr8__138683067 | C | G | Blood_Tier2 | 0.182 |
| chr20__36457105 | G | A | Blood_Tier1 | 0.294 |
| chr1__233558620 | T | C | Blood_Tier2 | 0.125 |
| chr6__51587361 | G | A | Blood_Tier1 | 0.095 |
| chr14__31631966 | T | C | Blood_Tier2 | 0.159 |
| chr17__72851459 | C | T | Blood_Tier2 | 0.123 |
| chr7__119749744 | G | T | Blood_Tier2 | 0.195 |
| chr11__21892126 | A | G | Blood_Tier2 | 0.125 |
| chr1__179142889 | A | G | Blood_Tier2 | 0.212 |
| chr2__11641476 | G | A | Blood_Tier2 | 0.104 |
| chr11__115664774 | C | T | Blood_Tier2 | 0.328 |
| chr16__15603964 | G | C | Blood_Tier2 | 0.304 |
| chr7__142480345 | C | A | Blood_Tier2 | 0.108 |
| chr12__95625935 | T | C | Blood_Tier3 | 0.429 |
| chr21__47278674 | G | A | Blood_Tier2 | 0.194 |
| chr13__90075654 | A | G | Blood_Tier3 | 0.286 |
| chr7__83446721 | C | T | Blood_Tier1 | 0.265 |
| chr4__171153608 | A | G | Blood_Tier2 | 0.094 |
| chr4__171058550 | C | T | Blood_Tier1 | 0.194 |
| chr6__14860391 | G | T | Blood_Tier1 | 0.301 |
| chr2__116215885 | G | A | Blood_Tier2 | 0.143 |
| chr7__130645986 | G | A | Blood_Tier2 | 0.292 |
| chr20__18999983 | A | G | Blood_Tier2 | 0.118 |
| chr18__5952052 | A | G | Blood_Tier3 | 0.273 |
| chr11__44366993 | C | T | Blood_Tier1 | 0.240 |
| chr3__153752188 | C | G | Blood_Tier2 | 0.200 |
| chr2__97487365 | T | C | Blood_Tier2 | 0.109 |
| chr2__211063832 | T | C | Blood_Tier2 | 0.167 |

| | | | | |
|---|---|---|---|---|
| chr7__141023239 | A | T | Blood_Tier2 | 0.317 |
| chr7__158650954 | A | G | Blood_Tier2 | 0.167 |
| chr11__63759135 | G | A | Blood_Tier2 | 0.224 |
| chr15__40074575 | T | C | Blood_Tier2 | 0.229 |
| chr6__11618300 | G | A | Blood_Tier2 | 0.133 |
| chr12__87948328 | C | T | Blood_Tier2 | 0.085 |
| chr7__100913097 | G | T | Blood_Tier2 | 0.086 |
| chr4__33840364 | A | G | Blood_Tier2 | 0.143 |
| chr2__208166004 | A | G | Blood_Tier2 | 0.194 |
| chr1__31427695 | A | G | Blood_Tier2 | 0.102 |
| chr5__24947460 | T | C | Blood_Tier2 | 0.125 |
| chr3__138220877 | C | T | Blood_Tier2 | 0.160 |
| chr4__131585241 | A | G | Blood_Tier2 | 0.208 |
| chr7__100913094 | A | G | Blood_Tier2 | 0.093 |
| chr10__113806265 | G | A | Blood_Tier2 | 0.133 |
| chr2__185184871 | C | T | Blood_Tier2 | 0.279 |
| chr2__213904794 | A | G | Blood_Tier3 | 0.182 |
| chrX__102549956 | G | A | Blood_Tier3 | 0.250 |
| chr2__29007303 | T | C | Blood_Tier2 | 0.167 |
| chr17__63146103 | T | C | Blood_Tier2 | 0.314 |
| chr18__74348273 | T | C | Blood_Tier2 | 0.099 |
| chr2__183127666 | T | G | Blood_Tier2 | 0.143 |
| chr10__93546985 | G | A | Blood_Tier2 | 0.340 |
| chr21__27791896 | C | T | Blood_Tier2 | 0.222 |
| chrX__126808708 | C | A | Blood_Tier2 | 0.250 |
| chr2__148198348 | G | T | Blood_Tier2 | 0.279 |
| chr8__109617037 | T | C | Blood_Tier2 | 0.208 |
| chrX__34206854 | C | T | Blood_Tier2 | 0.154 |
| chr13__81353501 | G | T | Blood_Tier2 | 0.148 |
| chr10__58835229 | G | C | Blood_Tier2 | 0.190 |
| chr14__77817270 | G | A | Blood_Tier2 | 0.195 |
| chr1__77567898 | G | C | Blood_Tier2 | 0.286 |
| chr1__100321512 | A | G | Blood_Tier2 | 0.182 |
| chr8__99588635 | G | C | Blood_Tier2 | 0.233 |
| chr11__35956176 | G | A | Blood_Tier2 | 0.081 |
| chr10__55246987 | C | T | Blood_Tier2 | 0.429 |
| chr13__97517961 | C | T | Blood_Tier2 | 0.123 |
| chr2__28920772 | G | A | Blood_Tier2 | 0.103 |
| chr5__142792230 | C | G | Blood_Tier2 | 0.250 |
| chr2__42979897 | C | T | Blood_Tier2 | 0.333 |
| chr9__118893196 | A | C | Blood_Tier2 | 0.136 |
| chr2__164553954 | G | A | Blood_Tier2 | 0.244 |
| chr4__32354940 | A | G | Blood_Tier2 | 0.167 |
| chr6__147249464 | A | T | Blood_Tier2 | 0.083 |
| chrX__152227031 | G | A | Blood_Tier3 | 0.174 |
| chr7__153381781 | C | T | Blood_Tier2 | 0.119 |

| | | | | |
|---|---|---|---|---|
| chr5__145397325 | A | G | Blood_Tier2 | 0.237 |
| chr7__142480349 | A | T | Blood_Tier2 | 0.103 |
| chr17__52417510 | A | G | Blood_Tier2 | 0.127 |
| chr12__104001108 | C | A | Blood_Tier2 | 0.196 |
| chr2__123856244 | A | G | Blood_Tier2 | 0.281 |
| chr2__166298204 | T | C | Blood_Tier2 | 0.087 |
| chr15__74911715 | A | G | Blood_Tier2 | 0.125 |
| chr5__77992219 | C | T | Blood_Tier2 | 0.171 |
| chr18__5194040 | A | G | Blood_Tier2 | 0.255 |
| chr8__79944479 | T | C | Blood_Tier2 | 0.138 |
| chrX__126940482 | G | A | Blood_Tier2 | 0.233 |
| chr4__126361802 | T | A | Blood_Tier3 | 0.125 |
| chr2__183771999 | A | T | Blood_Tier2 | 0.231 |
| chr7__54540043 | C | T | Blood_Tier2 | 0.280 |
| chr13__75165964 | G | A | Blood_Tier3 | 0.133 |
| chr3__52977475 | G | A | Blood_Tier2 | 0.237 |
| chr3__158785637 | T | C | Blood_Tier2 | 0.186 |
| chr1__236402269 | A | G | Blood_Tier2 | 0.194 |
| chrX__14888241 | T | C | Blood_Tier2 | 0.232 |
| chr11__82829818 | C | T | Blood_Tier2 | 0.320 |
| chr12__59238155 | C | T | Blood_Tier2 | 0.354 |
| chr8__97577593 | G | A | Blood_Tier3 | 0.143 |
| chr12__132446865 | A | G | Blood_Tier3 | 0.125 |
| chr4__123669774 | C | A | Blood_Tier2 | 0.239 |
| chr2__183127665 | T | G | Blood_Tier2 | 0.162 |
| chr11__24685015 | T | C | Blood_Tier2 | 0.146 |
| chrX__143418060 | T | C | Blood_Tier3 | 0.118 |
| chr12__11210419 | A | C | Blood_Tier2 | 0.063 |
| chr12__8149812 | C | T | Blood_Tier2 | 0.222 |
| chr7__88148508 | A | G | Blood_Tier3 | 0.375 |
| chr12__11210420 | G | A | Blood_Tier2 | 0.063 |
| chrX__99060469 | G | A | Blood_Tier2 | 0.211 |
| chr16__65890985 | T | C | Blood_Tier3 | 0.333 |
| chr6__82175337 | A | G | Blood_Tier3 | 0.125 |
| chr12__33853021 | C | T | Blood_Tier2 | 0.178 |
| chrX__97811601 | T | A | Blood_Tier2 | 0.100 |
| chr14__71731010 | G | T | Blood_Tier2 | 0.089 |
| chrX__130195968 | C | A | Blood_Tier2 | 0.065 |
| chr6__32432565 | C | T | Blood_Tier2 | 0.102 |
| chr6__84964258 | T | C | Blood_Tier3 | 0.160 |
| chr2__123918052 | T | C | Blood_Tier3 | 0.194 |
| chr13__85569601 | T | G | Blood_Tier3 | 0.190 |
| chr9__120691738 | C | T | Blood_Tier2 | 0.155 |
| chr9__118123925 | C | T | Blood_Tier2 | 0.197 |
| chr10__126339325 | T | C | Blood_Tier2 | 0.310 |
| chr7__57634252 | C | G | Blood_Tier2 | 0.087 |

| | | | | |
|---|---|---|---|---|
| chrX__123798197 | A | T | Blood_Tier2 | 0.129 |
| chr1__63581312 | T | C | Blood_Tier2 | 0.115 |
| chr21__39764905 | T | C | Blood_Tier2 | 0.315 |
| chr13__20371060 | C | A | Blood_Tier2 | 0.089 |
| chr3__179937559 | T | C | Blood_Tier3 | 0.214 |
| chr6__70361621 | T | A | Blood_Tier2 | 0.106 |
| chr20__40341458 | C | A | Blood_Tier2 | 0.170 |
| chr1__180893060 | C | T | Blood_Tier2 | 0.281 |
| chr1__43017274 | A | G | Blood_Tier3 | 0.133 |
| chr16__88255777 | C | T | Blood_Tier2 | 0.214 |
| chr1__220238441 | C | T | Blood_Tier3 | 0.106 |
| chr12__14130513 | C | T | Blood_Tier3 | 0.098 |
| chr6__156071675 | G | A | Blood_Tier3 | 0.222 |
| chr3__38961324 | T | C | Blood_Tier2 | 0.214 |
| chr7__122029852 | A | G | Blood_Tier3 | 0.214 |
| chr2__95585053 | G | C | Blood_Tier2 | 0.105 |
| chr21__44826808 | A | G | Blood_Tier3 | 0.190 |
| chr4__175404073 | T | C | Blood_Tier3 | 0.235 |
| chr6__32432575 | T | C | Blood_Tier3 | 0.091 |
| chr11__94335617 | C | T | Blood_Tier3 | 0.176 |
| chr11__102729686 | C | T | Blood_Tier2 | 0.318 |
| chr8__61215776 | C | T | Blood_Tier3 | 0.286 |
| chr14__93904956 | T | C | Blood_Tier2 | 0.349 |
| chr1__35656873 | A | T | Blood_Tier2 | 0.124 |
| chr5__166164023 | G | A | Blood_Tier2 | 0.247 |
| chr5__38148040 | G | A | Blood_Tier2 | 0.307 |
| chr1__107865400 | C | T | Blood_Tier2 | 0.271 |
| chr4__45093115 | A | C | Blood_Tier3 | 0.121 |
| chr7__43402191 | C | T | Blood_Tier2 | 0.135 |
| chr5__11265045 | T | A | Blood_Tier3 | 0.212 |
| chr3__25263891 | T | A | Blood_Tier2 | 0.164 |
| chrX__136332201 | T | C | Blood_Tier2 | 0.152 |
| chr11__17058370 | G | C | Blood_Tier2 | 0.260 |
| chr2__222988298 | C | A | Blood_Tier2 | 0.141 |
| chr1__203505501 | C | A | Blood_Tier2 | 0.208 |
| chr9__127654540 | A | C | Blood_Tier2 | 0.339 |
| chrX__107726986 | T | C | Blood_Tier2 | 0.333 |
| chr2__160880663 | C | T | Blood_Tier3 | 0.125 |
| chr17__61953832 | T | C | Blood_Tier2 | 0.074 |
| chr18__11198650 | T | C | Blood_Tier2 | 0.281 |
| chr4__161063756 | T | C | Blood_Tier2 | 0.265 |
| chr2__90265077 | A | G | Blood_Tier2 | 0.093 |
| chr3__37745882 | G | A | Blood_Tier2 | 0.114 |
| chr7__142153278 | T | C | Blood_Tier2 | 0.091 |
| chr10__59279600 | T | C | Blood_Tier3 | 0.273 |
| chr21__41193413 | C | A | Blood_Tier2 | 0.313 |

| | | | | |
|---|---|---|---|---|
| chr7__29940225 | C | T | Blood_Tier3 | 0.394 |
| chr2__194613483 | A | G | Blood_Tier3 | 0.158 |
| chr13__99419311 | G | A | Blood_Tier2 | 0.100 |
| chr13__106148454 | A | G | Blood_Tier2 | 0.172 |
| chr2__237158632 | C | A | Blood_Tier2 | 0.089 |
| chr2__222988297 | C | A | Blood_Tier2 | 0.139 |
| chr2__151857888 | A | T | Blood_Tier3 | 0.219 |
| chr10__27239514 | G | A | Blood_Tier2 | 0.127 |
| chr4__91726950 | G | A | Blood_Tier3 | 0.114 |
| chrX__49972551 | C | T | Blood_Tier2 | 0.078 |
| chr3__94257342 | C | T | Blood_Tier2 | 0.233 |
| chr3__163911788 | A | T | Blood_Tier2 | 0.138 |
| chr20__33824956 | C | T | Blood_Tier2 | 0.234 |
| chrX__121955937 | C | T | Blood_Tier3 | 0.119 |
| chr1__91688375 | T | G | Blood_Tier2 | 0.082 |
| chr8__127828459 | A | G | Blood_Tier3 | 0.250 |
| chr19__20799880 | A | G | Blood_Tier2 | 0.092 |
| chr19__20799879 | G | A | Blood_Tier2 | 0.092 |
| chrX__49972548 | G | A | Blood_Tier2 | 0.094 |
| chr12__11176428 | G | C | Blood_Tier2 | 0.079 |
| chr8__21514835 | T | C | Blood_Tier3 | 0.214 |
| chr2__25469028 | C | T | Blood_Tier3 | 0.224 |
| chr1__72999818 | A | G | Blood_Tier3 | 0.375 |
| chr14__30593452 | A | C | Blood_Tier3 | 0.161 |
| chr8__138681711 | A | T | Blood_Tier3 | 0.263 |
| chrX__61884169 | G | A | Blood_Tier2 | 0.101 |
| chrX__65369257 | T | G | Blood_Tier3 | 0.159 |
| chr7__14288496 | G | A | Blood_Tier3 | 0.071 |
| chr12__48861370 | C | T | Blood_Tier3 | 0.214 |
| chr3__54633194 | C | T | Blood_Tier2 | 0.097 |
| chr2__96516676 | G | C | Blood_Tier2 | 0.194 |
| chr12__26365989 | T | C | Blood_Tier2 | 0.208 |
| chr8__138681710 | A | T | Blood_Tier3 | 0.263 |
| chr9__129576038 | G | T | Blood_Tier3 | 0.241 |
| chr5__174418500 | T | C | Blood_Tier2 | 0.068 |
| chr1__27210525 | C | T | Blood_Tier3 | 0.118 |
| chr4__184500715 | G | T | Blood_Tier3 | 0.357 |
| chr7__108229549 | T | A | Blood_Tier3 | 0.214 |
| chrX__121955915 | A | C | Blood_Tier3 | 0.100 |
| chr3__36839096 | C | T | Blood_Tier3 | 0.200 |
| chr3__192399169 | G | T | Blood_Tier2 | 0.217 |
| chr16__65320183 | G | T | Blood_Tier3 | 0.122 |
| chr18__52712073 | A | G | Blood_Tier3 | 0.097 |
| chr9__86052888 | T | A | Blood_Tier2 | 0.203 |
| chr15__26686717 | G | T | Blood_Tier3 | 0.316 |
| chr10__27241290 | A | C | Blood_Tier2 | 0.101 |

| | | | | |
|---|---|---|---|---|
| chr6__116473002 | T | C | Blood_Tier3 | 0.158 |
| chr11__97875462 | A | G | Blood_Tier3 | 0.333 |
| chr5__165415490 | G | A | Blood_Tier3 | 0.500 |
| chr11__128142166 | T | G | Blood_Tier3 | 0.120 |
| chr2__18299562 | C | T | Blood_Tier3 | 0.138 |
| chr7__146112371 | A | C | Blood_Tier3 | 0.111 |
| chr3__54979489 | C | T | Blood_Tier2 | 0.189 |
| chr10__37533452 | C | T | Blood_Tier2 | 0.132 |
| chr10__19005639 | C | T | Blood_Tier3 | 0.133 |
| chr12__19795500 | A | G | Blood_Tier3 | 0.123 |
| chr5__46351383 | A | T | Blood_Tier3 | 0.154 |
| chr8__109849541 | G | A | Blood_Tier3 | 0.188 |
| chr7__57708912 | C | T | Blood_Tier3 | 0.082 |
| chr12__17994764 | A | T | Blood_Tier3 | 0.139 |
| chr17__72097012 | G | A | Blood_Tier3 | 0.152 |
| chr7__142112777 | T | A | Blood_Tier3 | 0.065 |
| chr22__20734031 | A | C | Blood_Tier3 | 0.200 |
| chr12__94310061 | C | T | Blood_Tier3 | 0.231 |
| chr4__55890156 | A | G | Blood_Tier3 | 0.200 |
| chr12__59590041 | C | T | Blood_Tier3 | 0.077 |
| chr6__72812320 | T | A | Blood_Tier3 | 0.239 |
| chrX__10342784 | G | A | Blood_Tier3 | 0.087 |
| chr14__106773919 | C | T | Blood_Tier3 | 0.050 |
| chr8__48253994 | A | T | Blood_Tier3 | 0.182 |
| chr5__109463919 | C | T | Blood_Tier3 | 0.200 |
| chr2__53347865 | A | G | Blood_Tier3 | 0.182 |
| chr12__81828865 | T | A | Blood_Tier3 | 0.300 |
| chr2__181402658 | A | G | Blood_Tier3 | 0.148 |
| chr8__41646406 | T | C | Blood_Tier3 | 0.228 |
| chr14__30443917 | G | T | Blood_Tier3 | 0.114 |
| chr5__25439865 | C | T | Blood_Tier3 | 0.167 |
| chr12__75466133 | A | G | Blood_Tier3 | 0.333 |
| chr18__28026822 | T | A | Blood_Tier3 | 0.087 |
| chr16__84230139 | C | T | Blood_Tier3 | 0.235 |
| chr16__80313752 | G | T | Blood_Tier3 | 0.064 |
| chr12__11201928 | C | A | Blood_Tier3 | 0.111 |
| chr7__142170774 | T | C | Blood_Tier3 | 0.068 |
| chr7__70914724 | A | T | Blood_Tier3 | 0.103 |
| chr22__33479147 | A | T | Blood_Tier3 | 0.133 |
| chr7__57701642 | A | T | Blood_Tier3 | 0.111 |
| chr3__193363447 | T | C | Blood_Tier3 | 0.267 |
| chr3__64061172 | G | A | Blood_Tier3 | 0.130 |
| chr4__128738035 | A | G | Blood_Tier3 | 0.231 |
| chr12__11210413 | T | A | Blood_Tier3 | 0.071 |
| chr6__157861588 | G | A | Blood_Tier3 | 0.265 |
| chr9__34743193 | C | T | Blood_Tier3 | 0.154 |

| | | | | |
|---|---|---|---|---|
| chr9__85102481 | T | C | Blood_Tier3 | 0.500 |
| chr3__58299010 | A | G | Blood_Tier3 | 0.194 |
| chr2__208730681 | A | G | Blood_Tier3 | 0.182 |
| chr12__17164642 | C | A | Blood_Tier3 | 0.167 |
| chr11__102543946 | G | T | Blood_Tier3 | 0.143 |
| chr6__147163290 | T | C | Blood_Tier3 | 0.176 |
| chr7__142112781 | G | A | Blood_Tier3 | 0.069 |
| chr3__100305657 | A | T | Blood_Tier3 | 0.071 |
| chr5__66754035 | A | G | Blood_Tier3 | 0.176 |
| chr7__142112780 | A | C | Blood_Tier3 | 0.068 |
| chr5__148127300 | A | T | Blood_Tier3 | 0.113 |
| chr2__170983636 | T | C | Blood_Tier3 | 0.114 |
| chr22__44812734 | G | C | Blood_Tier3 | 0.200 |
| chr11__818321 | G | T | Blood_Tier3 | 0.172 |
| chr2__194109480 | A | C | Blood_Tier3 | 0.136 |
| chrX__22267527 | C | A | Blood_Tier3 | 0.231 |
| chr8__74400493 | T | C | Blood_Tier3 | 0.160 |
| chr2__194109479 | G | T | Blood_Tier3 | 0.109 |
| chr5__90035124 | A | G | Blood_Tier3 | 0.200 |
| chr13__71965717 | T | C | Blood_Tier3 | 0.400 |
| chr12__71818168 | T | C | Blood_Tier3 | 0.325 |
| chr13__105774795 | A | G | Blood_Tier3 | 0.194 |
| chr12__11208089 | T | C | Blood_Tier3 | 0.069 |
| chr8__126979801 | C | A | Blood_Tier3 | 0.160 |
| chr17__22217028 | A | G | Blood_Tier3 | 0.169 |
| chr16__85272383 | T | C | Blood_Tier3 | 0.207 |
| chr4__167260078 | T | A | Blood_Tier3 | 0.235 |
| chr14__78702011 | G | A | Blood_Tier3 | 0.194 |
| chr9__118893195 | A | C | Blood_Tier3 | 0.140 |
| chr1__26062022 | T | C | Blood_Tier3 | 0.375 |
| chr15__27169321 | T | C | Blood_Tier3 | 0.125 |
| chr3__159270357 | A | G | Blood_Tier3 | 0.214 |
| chr2__124948132 | A | T | Blood_Tier3 | 0.108 |
| chr12__95000294 | C | A | Blood_Tier3 | 0.348 |
| chr10__5854557 | G | A | Blood_Tier3 | 0.418 |
| chr2__89078968 | T | G | Blood_Tier3 | 0.130 |
| chr10__23074430 | T | C | Blood_Tier3 | 0.167 |
| chr8__41412171 | T | C | Blood_Tier3 | 0.133 |
| chr7__142174972 | G | A | Blood_Tier3 | 0.091 |
| chr4__177646963 | C | T | Blood_Tier3 | 0.231 |
| chr18__48949657 | A | G | Blood_Tier3 | 0.093 |
| chr11__41614000 | G | A | Blood_Tier3 | 0.318 |
| chr1__247287449 | T | C | Blood_Tier3 | 0.326 |
| chr6__1083453 | G | A | Blood_Tier3 | 0.148 |
| chr8__126968905 | T | C | Blood_Tier3 | 0.188 |
| chr6__107437250 | C | T | Blood_Tier3 | 0.250 |

| | | | | |
|---|---|---|---|---|
| chr5__125454760 | A | T | Blood_Tier3 | 0.184 |
| chr9__79491520 | G | T | Blood_Tier3 | 0.275 |
| chr5__3489699 | A | G | Blood_Tier3 | 0.091 |
| chr8__34584827 | T | C | Blood_Tier3 | 0.150 |
| chrX__111336821 | G | C | Blood_Tier3 | 0.246 |
| chrX__61950819 | T | A | Blood_Tier3 | 0.092 |
| chr2__95585054 | T | A | Blood_Tier3 | 0.108 |
| chr9__132197219 | A | T | Blood_Tier3 | 0.048 |
| chr16__12463267 | A | T | Blood_Tier3 | 0.355 |
| chr19__22493999 | T | C | Blood_Tier3 | 0.167 |
| chr7__9011291 | A | G | Blood_Tier3 | 0.174 |
| chr10__8961853 | G | A | Blood_Tier3 | 0.167 |
| chr6__32601760 | A | G | Blood_Tier3 | 0.085 |
| chr14__27688768 | G | A | Blood_Tier3 | 0.226 |
| chr12__56932190 | T | C | Blood_Tier3 | 0.207 |
| chr13__58595985 | G | A | Blood_Tier3 | 0.231 |
| chr6__166140601 | T | G | Blood_Tier3 | 0.179 |
| chrX__105747658 | C | G | Blood_Tier3 | 0.311 |
| chr14__29818566 | T | C | Blood_Tier3 | 0.167 |
| chr7__151243817 | G | A | Blood_Tier3 | 0.250 |
| chr11__30101778 | T | C | Blood_Tier3 | 0.188 |
| chr2__133007221 | C | T | Blood_Tier3 | 0.094 |
| chr14__88428978 | A | C | Blood_Tier3 | 0.088 |
| chr17__55146827 | C | A | Blood_Tier3 | 0.246 |
| chr10__125358902 | A | G | Blood_Tier3 | 0.133 |
| chr13__81777002 | A | G | Blood_Tier3 | 0.214 |
| chr21__30330594 | G | C | Blood_Tier3 | 0.245 |
| chr17__16808692 | G | T | Blood_Tier3 | 0.288 |
| chr7__85996958 | A | T | Blood_Tier3 | 0.214 |
| chr1__73751623 | T | C | Blood_Tier3 | 0.200 |
| chr8__58904251 | G | A | Blood_Tier3 | 0.070 |
| chr1__46105145 | T | G | Blood_Tier3 | 0.118 |
| chr6__81335455 | T | C | Blood_Tier3 | 0.214 |
| chr4__133649937 | A | C | Blood_Tier3 | 0.143 |
| chr8__140603870 | T | G | Blood_Tier3 | 0.043 |
| chr20__26135983 | T | A | Blood_Tier3 | 0.075 |
| chr6__166593308 | T | A | Blood_Tier3 | 0.222 |
| chr1__248907327 | G | A | Blood_Tier3 | 0.071 |
| chr6__128779460 | C | T | Blood_Tier3 | 0.344 |
| chr3__120233618 | G | T | Blood_Tier3 | 0.194 |
| chrX__130838278 | T | G | Blood_Tier3 | 0.121 |
| chr16__33969302 | C | A | Blood_Tier3 | 0.076 |
| chr5__106638979 | T | C | Blood_Tier3 | 0.076 |
| chr14__106902786 | C | T | Blood_Tier3 | 0.115 |
| chr7__67019719 | C | T | Blood_Tier3 | 0.290 |
| chrX__71907477 | G | T | Blood_Tier3 | 0.167 |

| | | | | |
|---|---|---|---|---|
| chr11__38061266 | A | G | Blood_Tier3 | 0.300 |
| chr11__16074445 | T | A | Blood_Tier3 | 0.181 |
| chr20__54408043 | C | A | Blood_Tier3 | 0.156 |
| chr18__57933503 | C | T | Blood_Tier3 | 0.340 |
| chr3__19741718 | T | C | Blood_Tier3 | 0.158 |
| chr4__72193370 | T | C | Blood_Tier3 | 0.211 |
| chr9__10009074 | A | G | Blood_Tier3 | 0.273 |
| chr12__7168848 | T | A | Blood_Tier3 | 0.105 |
| chr7__131727222 | A | C | Blood_Tier3 | 0.250 |
| chr1__196140095 | T | C | Blood_Tier3 | 0.167 |
| chr4__31280818 | G | A | Blood_Tier3 | 0.180 |

#### TABLE S2: Variant Allele Fractions of 307 Validated Somatic Mutations

| gvarID | PB0 | PB1 | PB2 | M1 | G1 | G2 | B1 | B2 | T1 | T4.2 | T8.2 | cluster |
|---|---|---|---|---|---|---|---|---|---|---|---|---|
| chr1__107865400 | 28.21 | 35.27 | 32.54 | 37.13 | 43.42 | 44.84 | 4 | 3.14 | 5.34 | 11.3 | 3.11 | 1 |
| chr1__113621193 | 31.34 | 35.99 | 36.29 | 42.39 | 46.41 | 46.99 | 4.28 | 3.06 | 6.53 | 11.74 | 4.19 | 1 |
| chr1__170805330 | 30.54 | 34.33 | 32.1 | 43.78 | 46.22 | 44.7 | 4.96 | 2.84 | 5.36 | 11.28 | 3.99 | 1 |
| chr1__179142889 | 29.6 | 36.1 | 33.77 | 37.7 | 45.95 | 43.76 | 4.23 | 4.01 | 6.74 | 12.09 | 4.27 | 1 |
| chr1__180644884 | 3.15 | 9.27 | 8.84 | 10.7 | 12.95 | 12.45 | 1.22 | 0.65 | 0.08 | 1.88 | 0.21 | 5 |
| chr1__180893060 | 30.51 | 34.84 | 31.6 | 43.76 | 43.46 | 44.62 | 4.12 | 3.25 | 5.55 | 11.48 | 3.15 | 1 |
| chr1__186581699 | 28.78 | 35.52 | 34.34 | 42.09 | 45.58 | 45.64 | 3.84 | 2.7 | 5.81 | 11.8 | 3.27 | 1 |
| chr1__201313101 | 1.97 | 7.02 | 7.39 | 9.06 | 9.74 | 10.6 | 1.17 | 0.83 | 0.15 | 1.46 | 0.1 | 5 |
| chr1__216608363 | 29.16 | 37.11 | 33.4 | 41.66 | 46.06 | 45.97 | 4.58 | 3.35 | 5.21 | 12.6 | 3.82 | 1 |
| chr1__222054608 | 19.99 | 25.98 | 26.63 | 31.72 | 35.93 | 34.97 | 2.92 | 2.63 | 2.96 | 7.28 | 1.4 | 2 |
| chr1__231042202 | 29.62 | 36.19 | 33.79 | 43.78 | 45.33 | 47.59 | 3.85 | 3.25 | 5.54 | 11.76 | 3.85 | 1 |
| chr1__237498066 | 28.34 | 33.46 | 31.94 | 38.2 | 44.08 | 44.34 | 3.75 | 2.81 | 4.51 | 10.98 | 3.1 | 1 |
| chr1__244390672 | 20.93 | 23.45 | 20.48 | 28.78 | 33.14 | 33.03 | 2.37 | 2.17 | 3.08 | 7.48 | 1.97 | 2 |
| chr1__35656873 | 9.53 | 9.44 | 8.76 | 12.07 | 13.78 | 11.55 | 0.7 | 1.14 | 0.42 | 1.92 | 0.56 | 4 |
| chr1__37164781 | 29.27 | 34.63 | 34.2 | 40.87 | 44.18 | 45.74 | 4.44 | 3.49 | 5.79 | 12.55 | 3.08 | 1 |
| chr1__48839483 | 28.86 | 31.94 | 30.64 | 38.08 | 44.58 | 43.04 | 3.68 | 2.77 | 5.27 | 10.46 | 3.25 | 1 |
| chr1__53590585 | 22.36 | 27.16 | 24.77 | 30.03 | 36.21 | 34.58 | 3.13 | 2.68 | 3.36 | 8.12 | 2.11 | 2 |
| chr1__7358343 | 29.16 | 34.35 | 33.19 | 37.68 | 45.04 | 43.57 | 4.29 | 3.83 | 5.61 | 10.86 | 3.51 | 1 |
| chr1__77567898 | 28.88 | 33.75 | 32.61 | 39.16 | 44.33 | 45.23 | 4.13 | 3.08 | 5.74 | 11.94 | 3.58 | 1 |
| chr1__79469387 | 31.69 | 32.6 | 32.05 | 37.89 | 43.79 | 43.38 | 4.32 | 2.91 | 5.84 | 10.24 | 3.36 | 1 |
| chr1__86284096 | 26.88 | 33.04 | 30.38 | 38.82 | 42.24 | 41.39 | 3.71 | 2.75 | 5.46 | 11.32 | 3.18 | 1 |
| chr10__109656568 | 30.09 | 33.73 | 33.18 | 41.62 | 45.61 | 45.75 | 5.26 | 2.59 | 5.89 | 11.79 | 3.69 | 1 |
| chr10__125225251 | 21.43 | 25.99 | 24.03 | 30.13 | 33.44 | 35.87 | 2.84 | 2.44 | 2.87 | 6.13 | 2.2 | 2 |
| chr10__126339325 | 27.18 | 28.21 | 25.73 | 32.38 | 36.45 | 37.93 | 2.3 | 3.63 | 4.45 | 9.32 | 2.94 | 2 |
| chr10__131873809 | 29.04 | 34.49 | 34.12 | 38.76 | 45.49 | 44.83 | 3.82 | 3.27 | 5.64 | 11.42 | 3.57 | 1 |
| chr10__20491854 | 2.66 | 7.45 | 8.59 | 10.58 | 10.82 | 12.57 | 0.74 | 0.59 | 0.15 | 1.26 | 0.41 | 5 |
| chr10__37533452 | 4.83 | 10.91 | 12.12 | 14.14 | 14.78 | 14.91 | 1.33 | 1.24 | 0.77 | 2.61 | 0.53 | 5 |
| chr10__55246987 | 27.81 | 33.1 | 31.01 | 38.72 | 42.41 | 41.44 | 3.54 | 2.68 | 5.49 | 8.8 | 3.51 | 1 |
| chr10__5854557 | 24.07 | 32.32 | 29.62 | 35 | 37.4 | 39.52 | 6.1 | 5.92 | 7.68 | 13.3 | 5.34 | 1 |
| chr10__58835229 | 28.79 | 35.62 | 29.78 | 38.67 | 43.9 | 43.73 | 3.18 | 3.79 | 5.71 | 10.84 | 3.43 | 1 |
| chr10__71833394 | 19.96 | 24.12 | 22.05 | 30.72 | 33.17 | 35.11 | 2.81 | 2.13 | 3.47 | 8.25 | 2.41 | 2 |

| | | | | | | | | | | | |
|---|---|---|---|---|---|---|---|---|---|---|---|
| chr10__84756078 | 28.56 | 33.89 | 32.63 | 40.51 | 46.65 | 45.07 | 3.48 | 2.81 | 6.41 | 12.22 | 2.61 | 1 |
| chr10__86358071 | 27.04 | 30.31 | 31.58 | 39.32 | 42.01 | 41.5 | 4.15 | 2.51 | 5.09 | 11.27 | 3.16 | 1 |
| chr10__93546985 | 25.7 | 31.3 | 29.29 | 37.52 | 41.81 | 40.67 | 3.17 | 2.82 | 4.37 | 9.85 | 2.83 | 1 |
| chr10__95590751 | 12.75 | 18.84 | 17.97 | 21.57 | 23.29 | 25.32 | 2.95 | 1.45 | 0.94 | 3.57 | 1.28 | 3 |
| chr11__102729686 | 29.74 | 36.25 | 32.97 | 40.88 | 43.64 | 42.88 | 4.01 | 3.45 | 5.43 | 11.34 | 3.36 | 1 |
| chr11__115664774 | 26.38 | 27.47 | 27.27 | 36.58 | 40.24 | 38.42 | 2.82 | 2.06 | 4.91 | 10.14 | 3.26 | 2 |
| chr11__119595518 | 28.13 | 34.83 | 33.12 | 39.53 | 43.97 | 44.29 | 4.1 | 3.11 | 5 | 12.1 | 3.15 | 1 |
| chr11__123959954 | 29.4 | 34.57 | 30.95 | 33.93 | 41.58 | 42.32 | 3.49 | 3.25 | 5.11 | 10.67 | 3.27 | 1 |
| chr11__133074959 | 27.4 | 31.87 | 29.03 | 35.28 | 40.79 | 41.82 | 3.6 | 2.64 | 4.82 | 10.14 | 2.38 | 1 |
| chr11__133213386 | 23.21 | 28.63 | 24.32 | 28.66 | 35.9 | 35.16 | 3.23 | 2.72 | 3.04 | 7.48 | 2.11 | 2 |
| chr11__134641812 | 21.06 | 25.85 | 26.24 | 27.72 | 34.02 | 34.4 | 3.35 | 2.93 | 3.31 | 7.37 | 2.64 | 2 |
| chr11__16074445 | 17.72 | 22.08 | 19.07 | 22.42 | 26.6 | 26.79 | 1.84 | 2.12 | 2 | 4.82 | 1.21 | 2 |
| chr11__17058370 | 30.35 | 34.81 | 33.52 | 41.09 | 44.48 | 47.15 | 3.77 | 3.87 | 5.3 | 12.23 | 4 | 1 |
| chr11__18060193 | 30.61 | 34.37 | 30.77 | 39.5 | 46.09 | 44.82 | 3.59 | 3.45 | 6.31 | 9.7 | 3.21 | 1 |
| chr11__24024971 | 29.46 | 35.36 | 29.5 | 38.02 | 44.33 | 46.11 | 3.28 | 3.71 | 4.86 | 11.91 | 3.12 | 1 |
| chr11__35956176 | 4.68 | 11.14 | 9.99 | 12.49 | 13.84 | 14.2 | 1.74 | 1.18 | 0.31 | 2 | 0.43 | 5 |
| chr11__37183794 | 3.47 | 9.92 | 9.67 | 12.88 | 11.33 | 12.99 | 0.97 | 0.6 | 0.07 | 1.35 | 0.15 | 5 |
| chr11__3766585 | 24.87 | 26.64 | 27.48 | 33.08 | 39.91 | 37.22 | 2.62 | 3.02 | 4.26 | 10.16 | 2.84 | 2 |
| chr11__41614000 | 16.21 | 21.2 | 17.18 | 25.84 | 28.54 | 26.92 | 2.91 | 1.43 | 3.07 | 6.56 | 1.03 | 3 |
| chr11__44366993 | 29.77 | 35.53 | 33.9 | 41.24 | 45.23 | 44.28 | 4 | 3.14 | 5.83 | 11.99 | 3.36 | 1 |
| chr11__55523417 | 24.31 | 29.54 | 28.43 | 32.52 | 37.82 | 38.97 | 3.42 | 2.48 | 5.23 | 8.89 | 2.77 | 2 |
| chr11__62037391 | 29.34 | 34.2 | 34.09 | 39.52 | 42.7 | 44.41 | 3.78 | 3.85 | 4.85 | 12.48 | 3.24 | 1 |
| chr11__82829818 | 28.28 | 30.89 | 31.2 | 37.51 | 42.65 | 42.95 | 3.82 | 3.95 | 6.14 | 12.6 | 3.36 | 1 |
| chr11__91653831 | 31.43 | 34.76 | 31.9 | 39.95 | 44.87 | 45.98 | 4.37 | 3.35 | 4.96 | 11.8 | 3.47 | 1 |
| chr11__94335617 | 28.11 | 37 | 30.99 | 39.33 | 44.87 | 44.14 | 3.94 | 2.79 | 5.93 | 10.92 | 3.83 | 1 |
| chr11__99367399 | 28.45 | 34.84 | 32.8 | 40.48 | 45.4 | 46.27 | 3.58 | 3.31 | 5.48 | 11.45 | 3.83 | 1 |
| chr12__14130513 | 2.91 | 8.88 | 8.92 | 11.35 | 11.43 | 13.34 | 0.81 | 0.67 | 0.16 | 1.33 | 0.33 | 5 |
| chr12__26365989 | 27.52 | 31.94 | 33.72 | 40.25 | 42.51 | 42.39 | 3.05 | 2.58 | 6.05 | 9.46 | 3.48 | 1 |
| chr12__29141533 | 28.64 | 34.47 | 32.11 | 41.18 | 45.01 | 45.4 | 3.71 | 2.32 | 5.59 | 11.52 | 3.71 | 1 |
| chr12__33853021 | 27.96 | 33.32 | 28.46 | 37.33 | 43.69 | 43.57 | 3.6 | 2.05 | 4.58 | 10.98 | 3.52 | 1 |
| chr12__48861370 | 13.25 | 15.74 | 13.36 | 18.24 | 21.86 | 19.19 | 0.91 | 0.87 | 2 | 4.32 | 1.04 | 3 |
| chr12__59238155 | 27.26 | 31.84 | 30.1 | 35.97 | 42.48 | 42.89 | 3.41 | 2.35 | 4.93 | 11.19 | 3.31 | 1 |
| chr12__59303326 | 27.42 | 36.52 | 34.14 | 39.73 | 46.63 | 45.16 | 2.96 | 2.41 | 5.83 | 11.76 | 3.33 | 1 |
| chr12__80759750 | 21.07 | 23.49 | 24.38 | 29.03 | 30.27 | 32.82 | 2.54 | 1.88 | 4.01 | 8.63 | 2.86 | 2 |
| chr12__86416207 | 30.29 | 34.23 | 33.21 | 38.28 | 45.07 | 45.09 | 3.46 | 2.12 | 5.97 | 11.89 | 3.36 | 1 |
| chr12__87948328 | 2.52 | 9.72 | 9.34 | 11.97 | 12.82 | 14.61 | 1 | 0.91 | 0.07 | 1.6 | 0.36 | 5 |
| chr12__90867035 | 23.6 | 27.43 | 26.34 | 32.95 | 35.47 | 36.68 | 2.78 | 2.24 | 3.05 | 7.65 | 2.18 | 2 |
| chr13__105958180 | 29.14 | 35.9 | 33.4 | 40.01 | 44.66 | 45.92 | 3.89 | 3.43 | 5.76 | 12.37 | 3.72 | 1 |
| chr13__112600346 | 23.47 | 28.57 | 29.2 | 32.59 | 39.32 | 38.64 | 2.21 | 2.49 | 1.8 | 5.56 | 2.44 | 2 |
| chr13__35027179 | 29.24 | 32.58 | 31.94 | 39.52 | 43.94 | 44.7 | 3.93 | 3.23 | 4.87 | 10.99 | 2.93 | 1 |
| chr13__57257821 | 28.06 | 35 | 31.86 | 39.49 | 42.29 | 43.45 | 3.03 | 2.5 | 5.1 | 10.69 | 3.83 | 1 |
| chr13__58595985 | 27.8 | 31.36 | 30.6 | 39.4 | 44.37 | 41.86 | 2.99 | 2.82 | 4.93 | 11.29 | 3.14 | 1 |
| chr13__81353501 | 3.04 | 8.47 | 9.04 | 10.49 | 10.74 | 12.7 | 0.73 | 0.69 | 0.17 | 1.37 | 0.18 | 5 |
| chr13__83427582 | 23.54 | 27.11 | 25.19 | 31.26 | 36.54 | 35.56 | 2.94 | 2.39 | 3.04 | 6.97 | 2.41 | 2 |
| chr13__97517961 | 24.14 | 27.96 | 25.71 | 32.4 | 37.25 | 38.39 | 2.71 | 2.35 | 2.7 | 7.78 | 2.18 | 2 |
| chr14__26630906 | 19.59 | 23.11 | 21.24 | 24.58 | 29.64 | 27.96 | 2.19 | 1.87 | 3.15 | 7.39 | 1.61 | 2 |

| | | | | | | | | | | | |
|---|---|---|---|---|---|---|---|---|---|---|---|
| chr14__31631966 | 28.13 | 33.01 | 31.84 | 37.83 | 45.13 | 42.02 | 3.54 | 2.98 | 5.53 | 11.27 | 4.09 | 1 |
| chr14__33343977 | 26.79 | 30.26 | 26.6 | 37.33 | 38.74 | 35.96 | 3.3 | 2.19 | 5.51 | 10.96 | 2.89 | 2 |
| chr14__35339308 | 20.94 | 25.77 | 24.86 | 31.85 | 35.01 | 36.01 | 3 | 2.24 | 2.79 | 7.9 | 2.32 | 2 |
| chr14__50105572 | 28.29 | 34.43 | 32.28 | 42.87 | 45.48 | 45.48 | 3.91 | 4.94 | 5.63 | 13.05 | 3.17 | 1 |
| chr14__64495445 | 29.31 | 34.59 | 34.06 | 36.96 | 43.37 | 47.11 | 3.96 | 3.99 | 5.48 | 11.8 | 3.6 | 1 |
| chr14__77817270 | 29.56 | 33 | 32.45 | 39.58 | 45.19 | 43.79 | 3.54 | 4 | 5.52 | 11.34 | 3.39 | 1 |
| chr14__78702011 | 27.15 | 31 | 29.71 | 38.14 | 40.72 | 42.55 | 4.05 | 2.97 | 4.75 | 10.37 | 2.99 | 1 |
| chr14__82551829 | 28.73 | 32.72 | 32.34 | 39.6 | 42.84 | 44.29 | 3.33 | 3.05 | 5.75 | 10.98 | 3.45 | 1 |
| chr14__93904956 | 27.63 | 34.2 | 32.82 | 40.18 | 44.13 | 44.6 | 3.99 | 3.65 | 5.25 | 11.12 | 3.68 | 1 |
| chr15__24051898 | 29.18 | 32.91 | 33.36 | 38.9 | 45.22 | 45.89 | 3.63 | 3.83 | 5.55 | 10.04 | 3.52 | 1 |
| chr15__36040233 | 19.86 | 25.15 | 25.03 | 30.68 | 36.92 | 33.62 | 2.48 | 2.05 | 3.31 | 6.6 | 2.28 | 2 |
| chr15__77477740 | 30.45 | 33.86 | 32.57 | 38.08 | 44.14 | 44.33 | 3.66 | 3.13 | 5.4 | 10.42 | 3.26 | 1 |
| chr15__86765022 | 2.62 | 8.99 | 7.98 | 11.73 | 12.47 | 13.32 | 1.19 | 0.63 | 0.19 | 1.66 | 0.29 | 5 |
| chr15__94305134 | 30.21 | 35.05 | 33.76 | 41.54 | 43.77 | 45.27 | 4.54 | 2.92 | 6.86 | 11.46 | 3.55 | 1 |
| chr16__15603964 | 29.61 | 35.53 | 32.67 | 40.33 | 45.05 | 43.48 | 3.63 | 3.42 | 5.36 | 11.99 | 3.61 | 1 |
| chr16__24450792 | 20.72 | 27.58 | 24.61 | 29.21 | 33.35 | 34.92 | 2.09 | 2.5 | 3.7 | 8.2 | 2.46 | 2 |
| chr16__26597017 | 3.08 | 8.99 | 8.12 | 10.38 | 11.65 | 13.7 | 0.84 | 0.57 | 0.17 | 1.18 | 0.32 | 5 |
| chr16__50955663 | 29.41 | 36.6 | 32.86 | 39.69 | 44.88 | 45.6 | 5.19 | 3.84 | 5.86 | 12.05 | 3.69 | 1 |
| chr16__53424032 | 33.14 | 40.97 | 38.09 | 45.76 | 49.52 | 49.04 | 3.62 | 3.59 | 6.01 | 13.69 | 3.12 | 1 |
| chr16__6613470 | 28.93 | 32.76 | 31.58 | 38.04 | 43.55 | 43.2 | 3.93 | 2.95 | 5.94 | 11.03 | 3.27 | 1 |
| chr16__7739039 | 13.87 | 21.03 | 22.03 | 25.2 | 27.92 | 29.58 | 2.27 | 1.38 | 1.04 | 4.67 | 1.02 | 3 |
| chr16__80313752 | 3.11 | 8.5 | 8.88 | 10.23 | 10.95 | 11.94 | 1.49 | 0.93 | 0.2 | 1.25 | 0.18 | 5 |
| chr16__87305621 | 29.06 | 33.6 | 33.99 | 39.53 | 45.56 | 44.68 | 4.46 | 4.02 | 6.15 | 12.69 | 4.24 | 1 |
| chr16__88255777 | 21.29 | 25.14 | 25.45 | 28.41 | 35.72 | 33.99 | 3.13 | 2.88 | 2.95 | 7.53 | 2.19 | 2 |
| chr17__31969569 | 33.04 | 39.89 | 37.43 | 45.68 | 51 | 48.65 | 4.27 | 2.73 | 5.92 | 12.25 | 3.41 | 1 |
| chr17__52417510 | 3.08 | 9.49 | 8.45 | 11.18 | 12.03 | 11.6 | 1.1 | 0.93 | 0.11 | 1.59 | 0.37 | 5 |
| chr17__57689194 | 25.27 | 30.43 | 29.95 | 34.74 | 42.71 | 39.49 | 3.1 | 2.48 | 4.75 | 8.88 | 2.87 | 2 |
| chr17__63146103 | 27.18 | 34.05 | 34.73 | 39.01 | 43.04 | 43.07 | 3.97 | 3.53 | 5.8 | 10.78 | 2.9 | 1 |
| chr17__66672231 | 28.86 | 35.83 | 32.28 | 37.83 | 43.85 | 43.15 | 3.67 | 3.4 | 5.91 | 11.47 | 2.77 | 1 |
| chr17__68940187 | 22.69 | 26.67 | 24.96 | 31.76 | 37.29 | 35.81 | 2.8 | 1.7 | 3.88 | 7.55 | 2.52 | 2 |
| chr17__72097012 | 14.42 | 21.99 | 20.68 | 26.55 | 30.05 | 28.31 | 2.92 | 1.5 | 1.73 | 5.99 | 0.87 | 3 |
| chr17__72851459 | 9.1 | 9.48 | 8.72 | 11.6 | 13.46 | 12.35 | 0.78 | 0.85 | 0.81 | 2.34 | 0.54 | 4 |
| chr17__8280395 | 29.8 | 35.18 | 32.57 | 39.54 | 44.75 | 43.13 | 3.62 | 3.34 | 5.84 | 13.53 | 3.32 | 1 |
| chr18__11198650 | 29.39 | 34.82 | 32.2 | 41.89 | 44.25 | 46.67 | 3.8 | 3.42 | 6.11 | 11.58 | 3.24 | 1 |
| chr18__1392644 | 28.64 | 35.02 | 34.26 | 42.68 | 45.2 | 43.07 | 4.26 | 3.3 | 5.5 | 12.46 | 3.75 | 1 |
| chr18__5194040 | 28.7 | 34.91 | 34.01 | 37.57 | 43.75 | 44.87 | 3.91 | 2.95 | 6.13 | 12.36 | 3.6 | 1 |
| chr18__70491670 | 29.5 | 34.47 | 32.94 | 40.68 | 43.7 | 45.31 | 3.84 | 3.28 | 5.11 | 11.95 | 3.85 | 1 |
| chr18__75948056 | 2.57 | 8.69 | 8.86 | 11.1 | 12.85 | 12.29 | 0.92 | 0.66 | 0.02 | 1.42 | 0.44 | 5 |
| chr19__29093087 | 30.33 | 35.23 | 35.03 | 40.76 | 46.43 | 46.77 | 4.11 | 3.09 | 5.97 | 11.94 | 3.98 | 1 |
| chr19__48149785 | 14.29 | 21.06 | 18.99 | 23.71 | 26.14 | 26.56 | 2.48 | 2.64 | 1.45 | 4.39 | 0.87 | 3 |
| chr19__56255629 | 30.07 | 31.64 | 30.55 | 40.17 | 43.71 | 46.46 | 3.5 | 3.02 | 4.86 | 11.11 | 3.4 | 1 |
| chr2__123856244 | 19.03 | 24.61 | 24.83 | 27.83 | 32.98 | 33.73 | 1.89 | 2.07 | 4.05 | 8.5 | 2.47 | 2 |
| chr2__141131460 | 29.71 | 35.01 | 33.15 | 40.13 | 45.17 | 45.93 | 3.87 | 2.91 | 6.02 | 11.13 | 2.93 | 1 |
| chr2__148198348 | 28.48 | 32.74 | 30.2 | 37.88 | 41.27 | 41.79 | 3.59 | 2.44 | 5.59 | 10.62 | 3.01 | 1 |
| chr2__157173187 | 28.62 | 33.08 | 32.89 | 40.11 | 44.73 | 45.92 | 3.53 | 3.11 | 4.62 | 12.42 | 3.3 | 1 |
| chr2__15721915 | 29.77 | 36.99 | 33.76 | 40.62 | 44.82 | 46.3 | 4.07 | 3.23 | 5.13 | 11.39 | 3.47 | 1 |

| | | | | | | | | | | | |
|---|---|---|---|---|---|---|---|---|---|---|---|
| chr2__164563143 | 29.53 | 33.8 | 31.9 | 39.81 | 43.6 | 44 | 3.12 | 3.35 | 4.89 | 10.96 | 3.75 | 1 |
| chr2__176211302 | 29.32 | 34.08 | 33 | 38.73 | 45.13 | 43.65 | 3.9 | 2.85 | 5.87 | 12.88 | 3.69 | 1 |
| chr2__183771999 | 31.32 | 35.92 | 33.39 | 41.66 | 46.91 | 45.96 | 4.13 | 3.29 | 6.53 | 11.88 | 4.1 | 1 |
| chr2__185184871 | 15.01 | 21.99 | 20.01 | 25.7 | 28.19 | 28.35 | 2.43 | 1.66 | 1.66 | 5.1 | 1.12 | 3 |
| chr2__194613483 | 27.72 | 34.44 | 32.51 | 38.8 | 44.26 | 45.29 | 4.49 | 2.62 | 5.87 | 11.18 | 3.36 | 1 |
| chr2__198609762 | 21.34 | 28.9 | 25.56 | 30.98 | 33.69 | 36.13 | 3.29 | 2.2 | 3.19 | 6.73 | 2.23 | 2 |
| chr2__199917885 | 29.25 | 35.05 | 32.41 | 40.36 | 47.22 | 45.68 | 4.74 | 3.04 | 6.27 | 11 | 3.85 | 1 |
| chr2__200482779 | 29.09 | 34.33 | 31.78 | 38.78 | 44.25 | 44.57 | 3.71 | 3.19 | 5.33 | 11.53 | 4.08 | 1 |
| chr2__203150320 | 31 | 34.02 | 33.94 | 42.51 | 45.81 | 45.27 | 3.16 | 3.55 | 5.56 | 12.72 | 2.97 | 1 |
| chr2__222988297 | 29.79 | 34.27 | 32.3 | 39.44 | 43.59 | 46.98 | 4.12 | 3.05 | 5.62 | 11.08 | 3.08 | 1 |
| chr2__227183785 | 25.83 | 30.86 | 28.82 | 37.07 | 40.85 | 40.62 | 3.43 | 3.09 | 4.37 | 8.75 | 2.44 | 1 |
| chr2__227317709 | 32.79 | 37.22 | 34.76 | 41.66 | 47.47 | 47.16 | 4.95 | 2.15 | 5.93 | 11.69 | 3.95 | 1 |
| chr2__228258480 | 31.29 | 37.06 | 35.72 | 39.71 | 46.56 | 46.37 | 4.42 | 3.54 | 5.45 | 11.62 | 4.56 | 1 |
| chr2__237158632 | 3.17 | 8.81 | 8.46 | 9.61 | 9.87 | 14.11 | 0.61 | 0.57 | 0.07 | 1.17 | 0.3 | 5 |
| chr2__28226905 | 30.62 | 33.84 | 28.53 | 36.42 | 40.54 | 38.6 | 3.23 | 2.75 | 6.22 | 11.48 | 2.76 | 1 |
| chr2__42979897 | 27.7 | 34.41 | 31.53 | 40.56 | 43.69 | 45.26 | 3.85 | 3.51 | 5.83 | 11.53 | 3.4 | 1 |
| chr2__61401283 | 31.83 | 35.52 | 33.48 | 42.15 | 45.38 | 47.6 | 3.67 | 2.88 | 5.63 | 11.39 | 3.97 | 1 |
| chr2__96516676 | 29.61 | 34.09 | 30.32 | 39.08 | 44.22 | 43.9 | 3.84 | 3.37 | 5.76 | 11.08 | 2.66 | 1 |
| chr2__99565440 | 27.33 | 33.33 | 32.56 | 38.3 | 44.08 | 44.26 | 3.63 | 2.26 | 4.75 | 12.12 | 2.7 | 1 |
| chr20__33824956 | 27.87 | 32.4 | 32.56 | 40.11 | 42.79 | 43.95 | 3 | 2.56 | 4.93 | 11.31 | 3.22 | 1 |
| chr20__36061512 | 30.95 | 35.53 | 35.38 | 43.45 | 47.53 | 47.69 | 4.63 | 3.25 | 6.09 | 13.99 | 3.85 | 1 |
| chr20__36457105 | 29.91 | 32.03 | 32.83 | 38.84 | 44.93 | 44.53 | 3.35 | 3.25 | 5.58 | 11.02 | 3.62 | 1 |
| chr20__38230941 | 27.93 | 32.43 | 30.4 | 38.32 | 41.14 | 44.62 | 3.21 | 3.21 | 4.94 | 10.57 | 3.27 | 1 |
| chr20__39862424 | 31.91 | 34.33 | 31.61 | 41.19 | 45.72 | 42.72 | 4.45 | 3.97 | 5.63 | 11.88 | 3.46 | 1 |
| chr20__40341458 | 14.75 | 20.06 | 19.52 | 24.81 | 28.19 | 26.87 | 2.09 | 1.59 | 1.09 | 4.73 | 1.14 | 3 |
| chr20__46878505 | 31.92 | 36.23 | 34.79 | 41.19 | 46.48 | 46.16 | 3.75 | 2.78 | 6.75 | 12.23 | 3.44 | 1 |
| chr20__53456479 | 29 | 33.18 | 28.86 | 40.98 | 42.97 | 44.8 | 3.39 | 2.44 | 6.28 | 10.78 | 3.27 | 1 |
| chr20__53722528 | 30.41 | 31.87 | 31.54 | 39.57 | 45.82 | 43.74 | 3.85 | 2.65 | 6.86 | 10.71 | 3.52 | 1 |
| chr20__54408043 | 28.38 | 35.75 | 33.39 | 38.37 | 43.88 | 46.41 | 4.57 | 3.42 | 5.59 | 11.96 | 3.53 | 1 |
| chr20__57909208 | 2.98 | 9.35 | 9.68 | 10.25 | 11.58 | 11.96 | 0.82 | 1.17 | 0.25 | 1.95 | 0.34 | 5 |
| chr21__27807421 | 28.74 | 33.06 | 33.12 | 39.42 | 42.55 | 43.35 | 4.53 | 3.58 | 5.58 | 11.87 | 3.75 | 1 |
| chr21__30330594 | 28.89 | 32.8 | 31.81 | 37.85 | 43.83 | 41.96 | 3.88 | 3.68 | 5.63 | 11.44 | 3.45 | 1 |
| chr21__35721670 | 30.15 | 33.71 | 33.78 | 41.01 | 43.76 | 44.36 | 4.28 | 3.17 | 5.62 | 11.65 | 3.71 | 1 |
| chr21__39764905 | 28.34 | 34.18 | 32.58 | 39.3 | 43.31 | 42.59 | 4.07 | 2.95 | 5.34 | 10.54 | 3.62 | 1 |
| chr21__41193413 | 22.97 | 29.03 | 28.32 | 32.63 | 39.49 | 37.12 | 4.45 | 3.27 | 4.73 | 9.92 | 3.32 | 2 |
| chr21__47278674 | 29.8 | 36.16 | 33.85 | 40.84 | 44.89 | 45.91 | 4.37 | 3.99 | 5.06 | 11.75 | 3.38 | 1 |
| chr22__34167420 | 30.13 | 34.8 | 32.33 | 42.24 | 45.2 | 45.24 | 4.12 | 2.81 | 6.37 | 11.9 | 2.91 | 1 |
| chr22__40724186 | 30.67 | 34.76 | 32.14 | 41.11 | 43.72 | 44.45 | 3.99 | 3.67 | 5.04 | 11.54 | 3.39 | 1 |
| chr22__44385037 | 31.46 | 36.29 | 32.41 | 41.89 | 47.97 | 47.43 | 3.43 | 5.06 | 5.96 | 11.48 | 3.44 | 1 |
| chr22__46538743 | 28.65 | 34.83 | 31.37 | 37.31 | 45.08 | 43.13 | 3.93 | 3.74 | 4.72 | 11.48 | 3.26 | 1 |
| chr3__130030623 | 30.26 | 34.7 | 32.4 | 41.25 | 46.08 | 43.96 | 3.69 | 3.22 | 6.17 | 11.46 | 3.87 | 1 |
| chr3__138121382 | 29.53 | 35.75 | 32.92 | 39.97 | 44.56 | 44.87 | 4.45 | 3.3 | 5.69 | 12.68 | 3.4 | 1 |
| chr3__147908859 | 29.2 | 33.75 | 32.27 | 40.88 | 43.01 | 44.56 | 3.99 | 3.1 | 5.51 | 10.83 | 3.82 | 1 |
| chr3__153752188 | 29.46 | 35.23 | 32.56 | 40.35 | 43.93 | 45.23 | 4.41 | 2.86 | 5.72 | 12.59 | 2.8 | 1 |
| chr3__158785637 | 14.34 | 22.11 | 20.72 | 24.98 | 27.04 | 29.21 | 2.12 | 1.51 | 1.68 | 5.6 | 0.96 | 3 |
| chr3__163911788 | 3.53 | 10.6 | 10.5 | 12.32 | 14.51 | 14.03 | 1.24 | 1.03 | 0.1 | 1.43 | 0.37 | 5 |

| | | | | | | | | | | | |
|---|---|---|---|---|---|---|---|---|---|---|---|
| chr3__176430705 | 30.56 | 35.45 | 35.24 | 40.5 | 47.04 | 46.91 | 4.19 | 2.95 | 5.84 | 12.5 | 3.09 | 1 |
| chr3__192399169 | 30.3 | 33.83 | 34.47 | 39.54 | 44.82 | 45.2 | 4.15 | 3.15 | 5.76 | 11.07 | 3.01 | 1 |
| chr3__192960239 | 29.7 | 34.3 | 31.21 | 40.77 | 45.19 | 44.98 | 3.38 | 2.92 | 5.69 | 12.05 | 3.28 | 1 |
| chr3__21941024 | 28.38 | 32.78 | 32.47 | 37.83 | 45.12 | 43.66 | 4.36 | 2.86 | 5.58 | 10.99 | 3.45 | 1 |
| chr3__37745882 | 2.29 | 7.28 | 6.94 | 7.52 | 9.18 | 9.64 | 0.76 | 0.69 | 0.05 | 0.79 | 0.25 | 5 |
| chr3__38961324 | 29.53 | 37 | 31.79 | 42.82 | 46.03 | 45.04 | 4 | 3.46 | 5.37 | 10.75 | 3.37 | 1 |
| chr3__50930603 | 20.94 | 25.12 | 24.99 | 28.31 | 33.72 | 33.95 | 3.12 | 2.31 | 2.06 | 6.54 | 2.11 | 2 |
| chr3__52315833 | 30.34 | 35.17 | 33.75 | 41.04 | 42.22 | 45.51 | 4.09 | 3.93 | 5.29 | 12.01 | 3.29 | 1 |
| chr3__52977475 | 27.19 | 33.45 | 31.29 | 40.26 | 43.97 | 43.04 | 3.34 | 4.15 | 5.79 | 11.43 | 3.19 | 1 |
| chr3__54633194 | 3.29 | 8.4 | 8.9 | 11.09 | 11.19 | 12.56 | 1.09 | 0.82 | 0.17 | 1.83 | 0.22 | 5 |
| chr3__54979489 | 12.66 | 18.81 | 19.18 | 23.13 | 25.43 | 25.93 | 1.43 | 1.36 | 1.12 | 3.59 | 0.86 | 3 |
| chr3__71590489 | 22.89 | 25.98 | 26.87 | 30.27 | 35.77 | 35.55 | 3.41 | 3.26 | 4.44 | 10.42 | 2.81 | 2 |
| chr3__94257342 | 28.83 | 32.16 | 32.05 | 38.16 | 41.96 | 43.41 | 4.07 | 3.21 | 4.63 | 10.1 | 3.19 | 1 |
| chr4__102280086 | 29.37 | 35.29 | 31.88 | 41.01 | 45.39 | 44.55 | 3.58 | 2.49 | 5.74 | 11.35 | 3.31 | 1 |
| chr4__115824013 | 7.43 | 7.53 | 6.74 | 9.04 | 10.15 | 10.14 | 0.75 | 0.42 | 0.77 | 1.63 | 0.6 | 4 |
| chr4__11638973 | 19.71 | 24.16 | 24.05 | 31.36 | 35.32 | 35.74 | 2.51 | 2.56 | 3.46 | 7.81 | 2.54 | 2 |
| chr4__123669774 | 29.68 | 37.29 | 32.31 | 42.82 | 45.1 | 42.06 | 3.97 | 2.36 | 5.66 | 12.53 | 3.82 | 1 |
| chr4__137409760 | 28.42 | 35.23 | 32.04 | 39.72 | 45.85 | 46.16 | 4.17 | 3.32 | 6.11 | 13.31 | 3.42 | 1 |
| chr4__145441371 | 29.22 | 33.45 | 33.41 | 41.12 | 45.02 | 43.01 | 4.34 | 3.98 | 4.9 | 11.48 | 3.1 | 1 |
| chr4__148542277 | 27.99 | 33.74 | 33.61 | 39.57 | 45.32 | 43.16 | 3.57 | 2.72 | 5.28 | 12.8 | 3.15 | 1 |
| chr4__149877106 | 28.61 | 34.18 | 30.71 | 39.38 | 43.32 | 44.77 | 3.44 | 3.22 | 6.01 | 11.12 | 3.39 | 1 |
| chr4__157045476 | 28.8 | 32.7 | 32.13 | 39.13 | 42.72 | 42.81 | 5.27 | 3.73 | 5.68 | 12.71 | 4.09 | 1 |
| chr4__159243915 | 28.61 | 33.54 | 30.3 | 41.21 | 42.35 | 43.33 | 4.07 | 3.04 | 5.13 | 10.12 | 3.09 | 1 |
| chr4__161063756 | 30.2 | 34.71 | 32.69 | 38.65 | 46.3 | 44.93 | 4.59 | 2.86 | 5.52 | 12.32 | 3.96 | 1 |
| chr4__169089352 | 28.44 | 33.4 | 32.78 | 37.89 | 45.36 | 43.06 | 4.01 | 3.19 | 5.29 | 11.8 | 2.76 | 1 |
| chr4__171058550 | 20.43 | 25.19 | 23.27 | 31.03 | 32.74 | 34.2 | 2.62 | 2.92 | 2.82 | 7.7 | 2.15 | 2 |
| chr4__17857259 | 29.81 | 35.1 | 33.04 | 38.35 | 43.31 | 43.78 | 4.06 | 3.39 | 4.96 | 11.28 | 2.96 | 1 |
| chr4__183386037 | 24.92 | 29.31 | 29.31 | 34.72 | 38.21 | 39.99 | 3.3 | 1.73 | 4.29 | 9.35 | 2.55 | 2 |
| chr4__24092455 | 15.28 | 22.94 | 20.58 | 24.68 | 27.82 | 29.29 | 1.94 | 1.89 | 1.73 | 5.17 | 0.83 | 3 |
| chr4__24761759 | 31.05 | 36.59 | 34.14 | 40.2 | 46.22 | 44.59 | 4.21 | 3.35 | 4.96 | 11.5 | 3.12 | 1 |
| chr4__30984482 | 28.94 | 33.92 | 30.57 | 39.76 | 43.02 | 42.95 | 3.48 | 2.83 | 5.03 | 11.22 | 3.73 | 1 |
| chr4__31280818 | 3.28 | 9.29 | 9.08 | 11.82 | 12.37 | 12.71 | 1.14 | 0.83 | 0.03 | 1.52 | 0.38 | 5 |
| chr4__55441256 | 27.42 | 34.59 | 32.69 | 35.99 | 43.43 | 43.23 | 3.29 | 2.5 | 5.5 | 11.21 | 3.91 | 1 |
| chr4__81241530 | 22.15 | 27.22 | 31.12 | 35.3 | 38.17 | 39.76 | 3.73 | 2.36 | 3.43 | 9.63 | 2.28 | 2 |
| chr5__123025785 | 30.08 | 34.73 | 34.2 | 40.84 | 45.25 | 45.7 | 4.33 | 3.84 | 4.5 | 12.52 | 3.49 | 1 |
| chr5__125454760 | 30.38 | 34.19 | 33.08 | 42.36 | 44.3 | 44.94 | 4.16 | 3.07 | 5.82 | 11.36 | 3.57 | 1 |
| chr5__130022910 | 30.26 | 36.17 | 35.84 | 40.98 | 45.23 | 47.36 | 4.14 | 3.63 | 5.76 | 12.7 | 3.12 | 1 |
| chr5__133385008 | 28.14 | 33.03 | 31.43 | 38.88 | 43.99 | 44.45 | 3.5 | 3.56 | 5.03 | 11.82 | 3.22 | 1 |
| chr5__142792230 | 28.18 | 36.47 | 32.09 | 41.78 | 44.17 | 45.55 | 4.66 | 2.95 | 6.66 | 11.62 | 3.59 | 1 |
| chr5__144949915 | 29.8 | 35.71 | 33.48 | 41.68 | 44.13 | 44.14 | 4.43 | 3.36 | 4.82 | 11.36 | 3.19 | 1 |
| chr5__148127300 | 3.02 | 8.44 | 7.53 | 11.53 | 11.57 | 13.41 | 1.06 | 0.87 | 0 | 1.53 | 0.38 | 5 |
| chr5__166164023 | 29.08 | 35.42 | 31.51 | 39.99 | 44.37 | 44.77 | 3.84 | 3.22 | 5.04 | 11.39 | 3.73 | 1 |
| chr5__166531089 | 33.08 | 38.34 | 35.12 | 45.36 | 46.89 | 48.41 | 4.11 | 3.34 | 5.98 | 11.76 | 3.99 | 1 |
| chr5__174418500 | 2.45 | 8.07 | 7.39 | 9.98 | 11.8 | 12.77 | 1.32 | 0.64 | 0.24 | 1.47 | 0.22 | 5 |
| chr5__175742143 | 30.77 | 35.34 | 34.23 | 42.81 | 46.39 | 46.36 | 4.2 | 3.95 | 5.94 | 12.49 | 3.3 | 1 |
| chr5__26329781 | 29.5 | 34.02 | 32.78 | 39.88 | 45.38 | 43.51 | 3.72 | 3.57 | 5.1 | 12.49 | 3.48 | 1 |

| | | | | | | | | | | | |
|---|---|---|---|---|---|---|---|---|---|---|---|
| chr5__34012260 | 20.5 | 25.33 | 23.68 | 29.43 | 35.22 | 33.09 | 2.49 | 2.23 | 2.95 | 6.65 | 2.42 | 2 |
| chr5__42290327 | 28.3 | 33.71 | 31.82 | 37.73 | 43.55 | 44.73 | 3.29 | 2.79 | 5.7 | 11.15 | 3.41 | 1 |
| chr5__84711280 | 2.75 | 8.04 | 7.93 | 10.95 | 11.74 | 10.86 | 1.51 | 0.76 | 0.07 | 1.41 | 0.5 | 5 |
| chr5__88243810 | 31.47 | 35.09 | 34.27 | 41.44 | 43.66 | 44.65 | 3.77 | 4.72 | 6.57 | 12.5 | 3.48 | 1 |
| chr6__100169580 | 28.57 | 33.28 | 31.18 | 40.63 | 43.94 | 44.81 | 4.16 | 2.89 | 5.34 | 12.57 | 3.07 | 1 |
| chr6__11618300 | 4.98 | 10.36 | 11.49 | 13.59 | 14.89 | 16.25 | 0.92 | 1.12 | 0.84 | 3.12 | 0.32 | 5 |
| chr6__121901998 | 3.2 | 8.9 | 9.21 | 11.39 | 12.51 | 13.18 | 0.88 | 1 | 0.14 | 1.28 | 0.27 | 5 |
| chr6__126113509 | 29.28 | 35.01 | 33.31 | 41 | 46.38 | 45.49 | 3.89 | 3.62 | 5.45 | 12.44 | 3.96 | 1 |
| chr6__128779460 | 28.4 | 32.06 | 31.25 | 39.42 | 43.4 | 44.96 | 3.46 | 3.04 | 5.05 | 11.57 | 3.32 | 1 |
| chr6__150816248 | 28.47 | 33.2 | 33.11 | 39.7 | 43.44 | 44 | 3.92 | 3.72 | 5.67 | 11.77 | 3.26 | 1 |
| chr6__152647273 | 28.01 | 32.09 | 32.73 | 37.74 | 43.53 | 42.43 | 3.65 | 3.64 | 6 | 10.34 | 3.08 | 1 |
| chr6__157861588 | 28.26 | 31.74 | 31.88 | 38.33 | 41.44 | 42.85 | 3.92 | 3.81 | 4.93 | 11.58 | 3.05 | 1 |
| chr6__160372588 | 30.28 | 34.47 | 32.34 | 39.9 | 44.57 | 44.6 | 3.89 | 3.95 | 5.45 | 13.05 | 3.16 | 1 |
| chr6__16390236 | 28.32 | 36.47 | 33.13 | 41.76 | 45.6 | 45.6 | 4.35 | 3.99 | 5.48 | 12.44 | 3.84 | 1 |
| chr6__166226981 | 9.22 | 21.15 | 22.68 | 26.39 | 28.55 | 30.58 | 2.1 | 2.5 | 1.11 | 4.56 | 0.6 | 3 |
| chr6__20794608 | 30.17 | 34.05 | 31.9 | 38.64 | 43.9 | 45.27 | 3.57 | 3.29 | 6.5 | 11.33 | 3.62 | 1 |
| chr6__28105508 | 32.37 | 38.31 | 38.32 | 41.86 | 44.8 | 48.2 | 4.31 | 4.25 | 4.63 | 13.85 | 3.34 | 1 |
| chr6__44324229 | 27.85 | 32.97 | 31.68 | 41.22 | 46.02 | 43.41 | 4.17 | 3.79 | 4.8 | 12.84 | 3.42 | 1 |
| chr6__51587361 | 10.3 | 10.57 | 9.17 | 11.44 | 14.03 | 13 | 0.88 | 1.07 | 0.79 | 2.19 | 0.56 | 4 |
| chr6__69085881 | 28.26 | 32.4 | 30.57 | 37.91 | 42.97 | 41.79 | 3.86 | 2.6 | 5.14 | 11.4 | 3.39 | 1 |
| chr6__72812320 | 27.34 | 31.88 | 33.15 | 39.3 | 40.2 | 42.97 | 4.07 | 2.67 | 5.15 | 10.75 | 3.18 | 1 |
| chr6__79616543 | 29.32 | 33.45 | 31.4 | 41.71 | 44.72 | 43.99 | 4.11 | 2.59 | 6.27 | 11.68 | 3.75 | 1 |
| chr6__81591019 | 14.27 | 16.73 | 18.85 | 23.89 | 25.16 | 24.95 | 2.08 | 0.99 | 2.82 | 5.6 | 1.98 | 3 |
| chr7__126483539 | 29.51 | 33.12 | 31.32 | 39.65 | 44.28 | 43.3 | 3.78 | 2.38 | 4.99 | 10.61 | 2.77 | 1 |
| chr7__129772266 | 4.18 | 9.57 | 10.31 | 11.88 | 13.73 | 13.98 | 1.91 | 1.56 | 0.89 | 3.06 | 1.16 | 5 |
| chr7__130645986 | 29.99 | 34.99 | 34.55 | 41.43 | 45.98 | 45.17 | 3.8 | 4.09 | 6.11 | 12.51 | 2.94 | 1 |
| chr7__131572995 | 30.5 | 35.56 | 33.52 | 40.71 | 44.7 | 44.87 | 4.59 | 2.22 | 5.78 | 11.52 | 3.23 | 1 |
| chr7__141023239 | 28.01 | 33.65 | 31.77 | 39.71 | 47.41 | 46.08 | 3.29 | 3.62 | 5.92 | 11.61 | 2.81 | 1 |
| chr7__14288496 | 6.35 | 7.82 | 6.89 | 9.54 | 8.35 | 9.8 | 1.25 | 1.06 | 2.47 | 4.32 | 0.94 | 4 |
| chr7__153381781 | 15.43 | 20.27 | 21 | 24.99 | 28.93 | 28.97 | 2.37 | 1.31 | 1.49 | 4.73 | 0.95 | 3 |
| chr7__24212517 | 29.09 | 34.4 | 31.32 | 40.36 | 44.23 | 44.52 | 3.27 | 2.79 | 5.38 | 11.6 | 3.64 | 1 |
| chr7__29217270 | 31.6 | 35.16 | 34.5 | 40.14 | 46.59 | 46.32 | 4.09 | 3.05 | 4.87 | 11.74 | 4.02 | 1 |
| chr7__40599736 | 29.98 | 34.71 | 32.42 | 40.21 | 44.5 | 46.04 | 3.96 | 2.91 | 5.37 | 11.07 | 3.89 | 1 |
| chr7__43402191 | 3.33 | 9.05 | 8.88 | 11.39 | 12.62 | 12.47 | 0.92 | 0.95 | 0.16 | 1.32 | 0.26 | 5 |
| chr7__54540043 | 27.53 | 33.49 | 31.97 | 39.73 | 44.52 | 43.01 | 4.15 | 2.46 | 5.3 | 11.14 | 3.82 | 1 |
| chr7__67019719 | 30.25 | 34.84 | 33.08 | 39.23 | 45.28 | 44.62 | 3.78 | 3.39 | 5.42 | 12.56 | 3.23 | 1 |
| chr7__78431782 | 25.98 | 29.1 | 29.51 | 34.19 | 39.09 | 40.4 | 3.25 | 2.96 | 4.24 | 10.56 | 3.93 | 2 |
| chr7__82293864 | 31.45 | 37.08 | 32.72 | 39.52 | 48.56 | 44.75 | 4.56 | 3.49 | 5.8 | 11.71 | 3.68 | 1 |
| chr7__83446721 | 29.57 | 34.73 | 32.45 | 39.98 | 45.09 | 43.9 | 4.39 | 3.32 | 5.49 | 10.74 | 3.81 | 1 |
| chr8__109617037 | 27.95 | 36.04 | 30.15 | 40.22 | 44.41 | 44.09 | 3.88 | 3.21 | 5.96 | 12.08 | 3.61 | 1 |
| chr8__11185286 | 28.45 | 34.25 | 32.98 | 42.16 | 43.46 | 44.47 | 3.71 | 4.33 | 5.44 | 11.85 | 3.19 | 1 |
| chr8__126819256 | 29.63 | 35.77 | 33.57 | 40.76 | 43.84 | 41.83 | 3.97 | 3.43 | 4.75 | 11.33 | 3.13 | 1 |
| chr8__13478301 | 29.27 | 34.61 | 32.9 | 41.34 | 43.89 | 44.55 | 3.9 | 3.23 | 4.94 | 11.8 | 3.67 | 1 |
| chr8__138529288 | 29.34 | 36.46 | 33.14 | 39.72 | 45.41 | 46.92 | 4.05 | 3.23 | 5.71 | 11.15 | 4.05 | 1 |
| chr8__141389872 | 27.16 | 33.22 | 29.86 | 39 | 42.36 | 43.67 | 4.09 | 3.2 | 4.92 | 11.47 | 3.66 | 1 |
| chr8__17407813 | 25.29 | 31.31 | 28.48 | 36.34 | 41.22 | 41.13 | 3.7 | 3.06 | 4.58 | 9.38 | 3.3 | 1 |

| | | | | | | | | | | | |
|---|---|---|---|---|---|---|---|---|---|---|---|
| chr8__27484960 | 21.49 | 27.91 | 23.38 | 31.03 | 33.55 | 33.22 | 2.96 | 2.6 | 2.83 | 7.16 | 2.33 | 2 |
| chr8__28291900 | 29.68 | 35.4 | 29.98 | 39.71 | 44.92 | 44.68 | 3.22 | 3.63 | 5.27 | 11.53 | 3.32 | 1 |
| chr8__34502821 | 30.46 | 32.15 | 34.93 | 42.06 | 45.31 | 46.57 | 3.8 | 3.25 | 5.35 | 10.43 | 3.46 | 1 |
| chr8__41646406 | 30.18 | 32.74 | 31.48 | 38.91 | 43.58 | 41.86 | 3.2 | 3.1 | 4.68 | 10.41 | 4.19 | 1 |
| chr8__41745949 | 27.51 | 32.81 | 30.72 | 39.82 | 43.88 | 42.68 | 4.47 | 3.77 | 5.81 | 11.83 | 4.06 | 1 |
| chr8__47359871 | 29.19 | 34.44 | 33.36 | 39 | 42.66 | 44.57 | 3.67 | 3.6 | 5.94 | 11.78 | 3.64 | 1 |
| chr8__54468375 | 29.9 | 34.19 | 32.15 | 38.59 | 45.44 | 44.3 | 3.99 | 3.09 | 5.96 | 11.58 | 3.25 | 1 |
| chr8__69906311 | 2.87 | 9.31 | 9.28 | 9.98 | 11.07 | 13.24 | 1.02 | 0.87 | 0.16 | 1.6 | 0.38 | 5 |
| chr8__99588635 | 29.17 | 35.46 | 33.83 | 41.41 | 42.21 | 44.24 | 4.19 | 2.94 | 5.56 | 12.2 | 3.62 | 1 |
| chr9__100480485 | 27.06 | 32.62 | 31.07 | 38.2 | 43.94 | 41.6 | 3.17 | 3.66 | 5.26 | 10.6 | 3.4 | 1 |
| chr9__118123925 | 27.66 | 34.04 | 30.96 | 38.54 | 43.46 | 46.14 | 4.31 | 2.94 | 4.91 | 11.01 | 3.17 | 1 |
| chr9__120691738 | 18.99 | 24.6 | 23.64 | 27.99 | 32.9 | 31.14 | 3.34 | 2.57 | 2.18 | 5.92 | 1.98 | 2 |
| chr9__125193632 | 29.24 | 32.34 | 33.31 | 38.85 | 45.62 | 42.9 | 3.46 | 3 | 5.6 | 12.78 | 4.1 | 1 |
| chr9__13732822 | 27.89 | 35.12 | 33.39 | 40.76 | 45.92 | 46.57 | 4.51 | 3.56 | 5.32 | 10.6 | 3.83 | 1 |
| chr9__137585155 | 29.11 | 35.31 | 30.56 | 40.61 | 41.81 | 43.49 | 5.18 | 3.06 | 4.15 | 10.94 | 3.39 | 1 |
| chr9__19516480 | 30.45 | 35.55 | 33.72 | 40.29 | 44.35 | 44.8 | 5.14 | 2.76 | 6.04 | 12.45 | 4.01 | 1 |
| chr9__37342120 | 33.77 | 40.88 | 39.3 | 45.85 | 51.98 | 52.99 | 5.22 | 4.21 | 6.7 | 14.77 | 4.68 | 1 |
| chr9__79491520 | 24.87 | 28.63 | 26.17 | 33.35 | 38.62 | 36.33 | 2.45 | 2.83 | 4.93 | 9.84 | 2.91 | 2 |
| chr9__84717849 | 29 | 34.62 | 32.5 | 40.13 | 44.1 | 44.15 | 3.47 | 3.44 | 5.8 | 12.87 | 3.99 | 1 |
| chr9__8517886 | 30.3 | 34.74 | 32.49 | 41.74 | 45.59 | 42.76 | 4.8 | 3.57 | 5.41 | 12.55 | 3.81 | 1 |
| chr9__86052888 | 30.95 | 36.37 | 33.87 | 41.12 | 44.97 | 45.7 | 3.82 | 3.77 | 4.42 | 10.64 | 3.41 | 1 |
| chr9__88013515 | 27.25 | 34.3 | 30.16 | 38.58 | 43.81 | 42.16 | 3.75 | 3.16 | 5.42 | 10.28 | 2.83 | 1 |
| chrX__105747658 | 30.04 | 34.85 | 34.45 | 41.13 | 43.36 | 45.22 | 4.74 | 2.7 | 6.49 | 12.83 | 4.23 | 1 |
| chrX__107726986 | 23.53 | 27.28 | 25.5 | 31.68 | 37.54 | 37.26 | 2.71 | 2.34 | 2.86 | 8.23 | 2.08 | 2 |
| chrX__109490016 | 31.68 | 35.39 | 32.47 | 42.72 | 48.07 | 44.21 | 4.06 | 3.28 | 6.48 | 12.58 | 3.92 | 1 |
| chrX__111336821 | 23.67 | 29.07 | 26.17 | 31.18 | 33.23 | 36.05 | 2.84 | 2.87 | 3.2 | 7.26 | 2.36 | 2 |
| chrX__114116246 | 30.17 | 35.06 | 34.54 | 41.08 | 47.36 | 47.08 | 3.86 | 3.19 | 6.57 | 12.03 | 3.96 | 1 |
| chrX__123798197 | 22.06 | 27.08 | 24.68 | 28.65 | 32.77 | 34.65 | 2.26 | 1.89 | 3.43 | 5.66 | 1.95 | 2 |
| chrX__126808708 | 27.48 | 34.95 | 32.34 | 37.14 | 42.89 | 42.36 | 3.95 | 2.55 | 5.78 | 10.84 | 3.37 | 1 |
| chrX__126940482 | 28.87 | 36.64 | 32.31 | 39.78 | 45.63 | 42.26 | 5.05 | 2.73 | 6.03 | 11.31 | 3.91 | 1 |
| chrX__129344146 | 29.96 | 35.69 | 31.49 | 40.74 | 44.56 | 42.95 | 3.55 | 4.05 | 5.89 | 12.86 | 3.81 | 1 |
| chrX__130195968 | 2.91 | 10.03 | 9.86 | 12.68 | 12.38 | 13.76 | 1.38 | 0.99 | 0.25 | 1.46 | 0.26 | 5 |
| chrX__132397792 | 22.8 | 27.37 | 24.69 | 32.74 | 34.81 | 36.66 | 3.7 | 2.51 | 4.09 | 8.63 | 1.77 | 2 |
| chrX__136332201 | 2.65 | 9.25 | 10.11 | 11.74 | 11.31 | 13.15 | 0.83 | 1.06 | 0.17 | 1.37 | 0.3 | 5 |
| chrX__137615166 | 31.61 | 35.32 | 31.87 | 41.42 | 43.82 | 45.35 | 4.02 | 3.97 | 7.36 | 12.42 | 2.56 | 1 |
| chrX__147904819 | 27.18 | 34.36 | 31.32 | 37.92 | 43.71 | 42.17 | 3.77 | 3.16 | 6.09 | 11.11 | 4.06 | 1 |
| chrX__14888241 | 30.35 | 36.6 | 32.98 | 39.57 | 42.66 | 43.77 | 4.34 | 2.47 | 5.89 | 12.85 | 3.57 | 1 |
| chrX__14959214 | 5.59 | 5.64 | 6.32 | 7.36 | 8.26 | 6.56 | 0.71 | 0.5 | 3 | 4.41 | 0.89 | 4 |
| chrX__24604702 | 29.3 | 35.99 | 33.19 | 40.53 | 43.55 | 43.55 | 3.66 | 3.69 | 5.52 | 12.36 | 3.44 | 1 |
| chrX__34047538 | 27.19 | 36.3 | 36.47 | 40.56 | 46.01 | 44.69 | 4.4 | 3.35 | 6.45 | 12.99 | 4.55 | 1 |
| chrX__34206854 | 2.41 | 7.89 | 7.64 | 8.82 | 9.43 | 11 | 0.6 | 0.44 | 0.05 | 1.29 | 0.09 | 5 |
| chrX__34276891 | 31.76 | 34.88 | 33.11 | 40.85 | 46 | 47.12 | 4.61 | 3.14 | 6.23 | 11.42 | 4.58 | 1 |
| chrX__66876515 | 29.25 | 35.92 | 31.33 | 39.6 | 44.36 | 43.2 | 3.19 | 3.18 | 6.35 | 13.1 | 4.07 | 1 |
| chrX__84392105 | 29.93 | 35.7 | 36.37 | 41.51 | 44.03 | 44.34 | 4.13 | 3.29 | 6.48 | 12.66 | 3.98 | 1 |
| chrX__97811601 | 3.64 | 9.38 | 9.47 | 11.44 | 11.9 | 13.1 | 1.11 | 0.9 | 0.21 | 1.56 | 0.26 | 5 |
| chrX__99060469 | 28.76 | 35.23 | 34.28 | 38.61 | 44.02 | 44.97 | 3.51 | 3.1 | 6.53 | 12.51 | 3.98 | 1 |

#### TABLE S3: Candidate driver definitions

<table>
<tr><th>GENE<br>[TRANSCIPT]</th><th>INCLUSION CRITERIA</th></tr>
<tr><td></td><td></td></tr>
<tr><td>DNMT3A<br>[NM_022552]</td><td>RTV: frameshift|nonsense|splice-site<br>KHV:<br>F290I|F290C|V296M|P307S|P307R|R326H|R326L|R326C|R326S|G332R|G332E|V339A|V339M|V339G|L344Q|L344P|R366P|R366H|R366G|A368T|A368V|R379H|R379C|I407T|I407N|I407S|F414L|F414S|F414C|A462V|K468R|C497G|C497Y|Q527H|Q527P|Y533C|S535F|C537G|C537R|G543A|G543S|G543C|L547H|L547P|L547F|M548I|M548K|G550R|W581R|W581G|W581C|R604Q|R604W|R635W|R635Q|S638F|G646V|G646E|L653W|L653F|I655N|V657A|V657M|R659H|Y660C|V665G|V665L|M674V|R676W|R676Q|G685R|G685E|G685A|D686Y|D686G|R688H|G699R|G699S|G699D|P700L|P700S|P700R|P700Q|P700T|P700A|D702N|D702Y|V704M|V704G|I705F|I705T|I705S|I705N|G707D|G707V|C710S|C710Y|S714C|V716D|V716F|V716I|N717S|N717I|P718L|R720H|R720G|K721R|K721T|Y724C|R729Q|R729W|R729G|F731C|F731L|F731Y|F731I|F732del|F732C|F732S|F732L|E733G|E733A|F734L|F734C|Y735C|Y735N|Y735S|R736H|R736C|R736P|L737H|L737V|L737F|L737R|A741V|P742P|P743R|P743L|R749C|R749L|R749H|R749G|F751L|F751C|F752del|F752C|F752L|F752I|F752V|W753G|W753C|W753R|L754P|L754R|L754H|F755S|F755I|F755L|M761I|M761V|G762C|V763I|S770L|S770W|S770P|R771Q|F772I|F772V|L773R|L773V|E774K|E774D|E774G|I780T|D781G|R792H|W795C|W795L|G796D|G796V|N797Y|N797H|N797S|P799S|P799R|P799H|R803S|R803W|P804L|P804S|K826R|S828N|K829R|T835M|N838D|K841Q|Q842E|P849L|D857N|W860R|E863D|F868S|G869S|G869V|M880V|S881R|S881I|R882H|R882P|R882C|R882G|A884P|A884V|Q886R|L889P|L889R|G890D|G890R|G890S|V895M|P896L|V897G|V897D|R899L|R899H|R899C|L901R|L901H|P904L|F909C|P904Q|A910P|C911R|C911Y</td></tr>
<tr><td>TET2<br>[NM_001127208]</td><td>RTV: frameshift|nonsense|splice-site<br>KHV: missense_mutations_in_catalytic_domains_(p.1104-1481_and_1843-2002)</td></tr>
<tr><td>ASXL1<br>[NM_015338]</td><td>RTV: frameshift|nonsense|splice-site (only in exon 11-12)<br>KHV: none</td></tr>
<tr><td>TP53<br>[NM_001126112]</td><td>RTV: frameshift|nonsense|splice-site<br>KHV:<br>S46F|G105C|G105R|G105D|G108S|G108C|R110L|R110C|T118A|T118R|T118I|S127F|S127Y|L130V|L130F|K132Q|K132E|K132W|K132R|K132M|K132N|F134V|F134L|F134S|C135W|C135S|C135F|C135G|C135Y|Q136K|Q136E|Q136P|Q136R|Q136L|Q136H|A138P|A138V|A138A|A138T|T140I|C141R|C141G|C141A|C141Y|C141S|C141F|C141W|V143M|V143A|V143E|L145Q|W146C|W146L|L145R|V147G|P151T|P151A|P151S|P151H|P151R|P152S|P152R|P152L|T155P|T155A|V157F|R158H|R158L|A159V|A159P|A159S|A159D|A161T|A161D|Y163N|Y163H|Y163D|Y163S|Y163C|K164E|K164M|K164N|K164P|H168Y|H168P|H168R|H168L|H168Q|M169I|M169T|M169V|E171K|E171Q|E171G|E171A|E171V|E171D|V172D|V173M|V173L|V173G|R174W|R175G|R175C|R175H|C176R|C176G|C176Y|C176F|C176S|P177R|P177R|P177L|H178D|H178P|H178Q|H179Y|H179R|H179Q|R181C|R181Y|D186G|G187S|P190L|P190T|H193N|H193P|H193L|H193R|L194F|L194R|I195F|I195N|I195T|R196P|V197L|G199V|Y205N|Y205C|Y205H|D208V|R213Q|R213P|R213L|R213Q|H214D|H214R|S215G|S215I|S215R|V216M|V217G|Y220N|Y220H|Y220S|Y220C|E224D|I232F|I232N|I232T|I232S|Y234N|Y234H|Y234S|Y234C|Y236N|Y236H|Y236C|M237V|M237K|M237I|C238R|C238G|C238Y|C238W|N239T|N239S|S241Y|S241C|S241F|C242G|C242Y|C242S|C242F|G244S|G244C|G244D|G245S|G245R|G245C|G245D|G245A|G245V|G245S|M246V|M246K|M246R|M246I|N247I|R248W|R248G|R248Q|R249G|R249W|R249T|R249M|P250L|I251N|L252P|I254S|I255F|I255N|I255S|L257Q|L257P|E258K|E258Q|D259Y|S261T|G262D|G262V|L265P|G266R|G266E|G266V|R267W|R267Q|R267P|E271K|V272M|V272L|R273S|R273G|R273C|R273H|R273P|R273L|V274F|V274D|V274A|V274G|V274L|C275Y|C275S|</td></tr>
</table>

| | |
|---|---|
| | C275F\|A276P\|C277F\|C277Y\|P278T\|P278A\|P278S\|P278H\|P278R\|P278L\|G279E\|R280G\|R280K\|R280T\|R280I\|R280S\|D281N\|D281H\|D281Y\|D281G\|D281E\|R282G\|R282W\|R282Q\|R282P\|E285K\|E285V\|E286G\|E286V\|E286K\|K320N\|L330R\|G334V\|R337C\|R337L\|A347T\|L348F\|T377P |
| *JAK2* [NM_004972] | **RTV:** none<br>**KHV:**<br>N533D\|N533Y\|N533S\|H538R\|K539E\|K539L\|I540T\|I540V\|V617F\|R683S\|R683G\|del/ins537-539L\|del/ins538-539L\|del/ins540-543MK\|del/ins540-544MK\|del/ins541-543K\|del542-543\|del543-544\|ins11546-547 3717 |
| *SF3B1* [NM_012433] | **RTV:** none<br>**KHV:**<br>G347V\|R387W\|R387Q\|E592K\|E622D\|Y623C\|R625L\|R625C\|R625G\|H662Q\|H662D\|T663I\|K666N\|K666T\|K666E\|K666R\|K700E\|V701F\|A708T\|G740R\|G740E\|A744P\|D781G\|E783K\|R831Q\|L833F\|E862K\|R957Q |
| *GNB1* [NM_002074] | **RTV:** none<br>**KHV**: K57N\|K57M\|K57E\|K57T\|I80T\|I80N |
| *CBL* [NM_005188] | **RTV:** none<br>**KHV**: RING_finger_missense_p.381_421 |
| *SFRS2* [NM_003016] | **RTV:** none<br>**KHV**: Y44H\|P95H\|P95L\|P95T\|P95R\|P95A\|P107H\|P95fs |
| *GNAS* [NM_016592] | **RTV:** none<br>**KHV**:<br>R201(844)S\|R201(844)C\|R201(844)H\|R201(844)L\|Q227(870)K\|Q227(870)R\|Q227(870)L\|Q227(870)H\|R374(1017)C |
| *BRCC3* [NM_024332] | **RTV**: frameshift\|nonsense\|splice-site<br>**KHV:** none |
| *CREBBP* [NM_004380] | **RTV**: frameshift\|nonsense\|splice-site<br>**KHV**:<br>D1435E\|R1446L\|R1446H\|R1446C\|Y1450C\|P1476R\|Y1482H\|H1487Y\|W1502C\|Y1503D\|Y1503H\|Y1503F\|S1680del |
| *NRAS* [NM_002524] | **RTV:** none<br>**KHV:**<br>G12S\|G12R\|G12C\|G12N\|G12P\|G12Y\|G12D\|G12A\|G12V\|G12E\|G13S\|G13R\|G13C\|G13N\|G13P\|G13Y\|G13D\|G13A\|G13V\|G13E\|G60E\|G60R\|Q61R\|Q61L\|Q61K\|Q61P\|Q61H\|Q61Q |
| *RAD21* [NM_006265] | **RTV**: frameshift\|nonsense\|splice-site<br>**KHV**: R65Q\|H208R\|Q474R |
| *U2AF1* [NM_006758] | **RTV**: none<br>**KHV**: D14G\|S34F\|S34Y\|R35L\|R156H\|R156Q\|Q157R\|Q157P |
| *PPM1D** [NM_003620] | **RTV**: frameshift\|nonsense\|splice-site (only in exon 5 or 6)<br>KHV: none |

**Table S3: Gene-specific variant inclusion criteria for 16 genes recurrently mutated in elderly patients:** Definitions are taken from Jaiswal *et al*. [Jaiswal 2017].
RTV: Rare Truncating Variant: a non-specific annotation indicating a type of truncating event, i.e.: 'frameshift', 'nonsense', 'splice-site'. Recurrent truncating events in hematological malignancies are annotated as KHV.
KHV: Known Hotspot Variant: Variants recurrently reported to be mutated in hematological malignancies.
